# Dissecting the structural and functional roles of a vicinal iron-binding site in encapsulated ferritins

**DOI:** 10.1101/785121

**Authors:** Cecilia Piergentili, Jennifer Ross, Didi He, Kelly J. Gallagher, Will A. Stanley, Laurène Adam, C. Logan Mackay, Kevin J. Waldron, David J. Clarke, Jon Marles-Wright

**Affiliations:** School of Natural and Environmental Sciences, Newcastle University. Newcastle upon Tyne, NE1 7RU; EaStCHEM School of Chemistry, The University of Edinburgh, Joseph Black Building, David Brewster Road, Edinburgh, Scotland, EH9 3FJ; Institute of Quantitative Biology, Biochemistry and Biotechnology, School of Biological Sciences, The University of Edinburgh, Max Born Crescent, Edinburgh, EH9 3BF; Biosciences Institute, Newcastle University, Newcastle upon Tyne, NE2 4HH

## Abstract

Encapsulated ferritins belong to the universally distributed ferritin superfamily, which function as iron detoxification and storage systems. Encapsulated ferritins have a distinct annular structure and must associate with an encapsulin nanocage to form a competent iron store that is capable of holding significantly more iron than classical ferritins. The catalytic mechanism of iron oxidation in the ferritin family is still an open question, due to differences in organization of the ferroxidase catalytic site and secondary metal binding sites vicinal to this. We have previously identified a metal binding site on the inner surface of the *Rhodospirillum rubrum* encapsulated ferritin at the interface between the two-helix subunits and proximal to the ferroxidase center. Here we present a comprehensive structural and functional study to investigate the functional relevance of this proposed iron entry site by means of enzymatic assays, mass-spectrometry, and X-ray crystallography. We show that catalysis occurs in the ferroxidase center and suggest a dual role for the secondary site, which both serves to attract metal ions to the ferroxidase center and acts as a flow-restricting valve to limit the activity of the ferroxidase center. Moreover, confinement of encapsulated ferritins within the encapsulin nanocage, while enhancing the ability of the encapsulated ferritin to undergo catalysis, does not influence the function of the secondary site.

## Introduction

The ferritin superfamily consists of a number of structurally and functionally distinct members, each of which is built around a four-helix bundle scaffold for an iron-binding di-iron ferroxidase active site (Andrews, 2010). The quaternary structure of these protein families shows a high degree of variation: from monomeric rubrerythrin proteins (Cardenas et al., 2016; Dillard et al., 2011); dimeric iron-mineralizing encapsulin-associated firmicute (IMEF) proteins (Giessen et al., 2019); dodecameric DNA Protection in Starved (DPS) cells proteins (Grant et al., 1998; Martinez and Kolter, 1997; Pesek et al., 2011); the 24-meric classical ferritins (Ftn) and bacterioferritins (Bfr) (Andrews, 1998; Frolow et al., 1994; Harrison and Arosio, 1996; Pfaffen et al., 2013); and the decameric encapsulated ferritins (EncFtn) (He et al., 2019, 2016). The rubrerythrin and DPS proteins play key roles in the protection of cells from oxidative stress, while DPS and the other members of the ferritin superfamily are vital iron stores (Almiron et al., 1992; Chiancone and Ceci, 2010; Choi et al., 2000; De Martino et al., 2016; Martinez and Kolter, 1997; Nair and Finkel, 2004; Ratnayake et al., 2000; Sztukowska et al., 2002). These latter proteins oxidize and convert free Fe(II) to inert Fe(III) oxyhydroxide or Fe(III) phosphate mineral forms that provide a reservoir of bioavailable iron (Chasteen and Harrison, 1999). The nanocages formed by the quaternary structure of DPS, Ftn, and Bfr allow iron oxidation and mineralization to occur within a biochemically privileged environment within the cell, which protects the host cell from iron-induced oxidative stress. Interestingly, the recently described IMEF and EncFtn proteins both have ferroxidase activity, but they do not form nanocage structures on their own, instead they must be localized to the lumen of encapsulin nanocages to form a competent iron-store (Giessen et al., 2019; He et al., 2019, 2016). EncFtn and IMEF are directed into encapsulins by a terminal localization sequence that binds to the interior wall of encapsulin. The EncFtn proteins form a distinct homodecameric annular assembly, with a pentamer of dimers topology (Figure 1A). One of the dimer interfaces forms a stable back-to-back conformation, exposing the metal ion binding residues (Figure 1B), which reconstitute a unique symmetrical ferroxidase center across a dimer interface when iron is bound (Figure 1C) (Ross et al., 2020).

**Figure 1.**
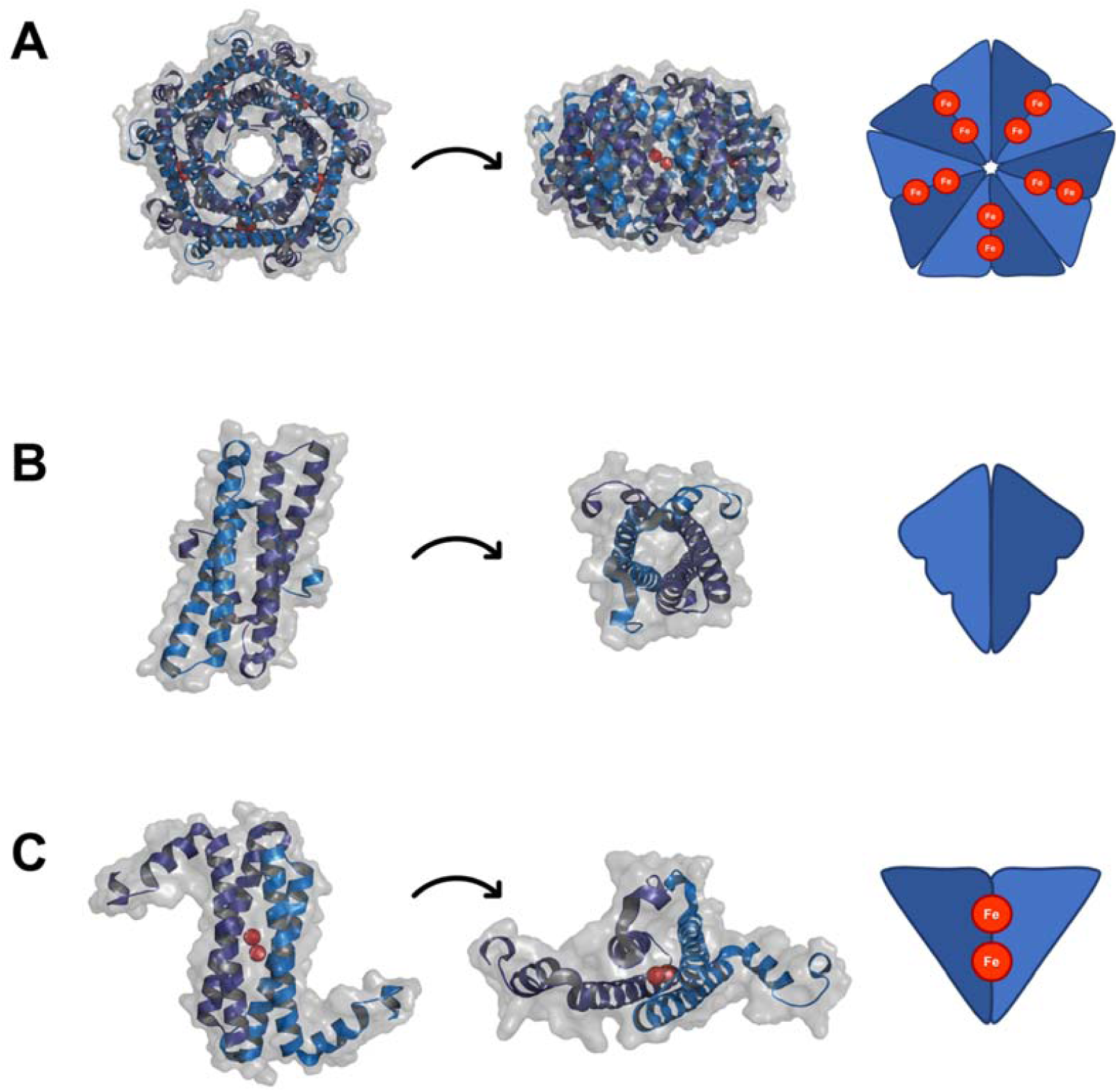
EncFtn subcomplex architecture. (**A**) Annular decamer structure of EncFtn shown with transparent solvent accessible surface rendered over cartoon secondary structure, with a simplified representation of topology. (**B**) The non-FOC dimer with cartoon secondary structure and transparent surface, and a simplified representation. (**C**) The non-FOC dimer in cartoon with two coordinated iron ions (shown as red spheres) and a simplified FOC cartoon is shown with two iron ions as red circles.

The catalytic route to iron mineralization in the different ferritin iron-stores is still a subject of vigorous debate (Baaghil et al., 2003; Chasteen and Harrison, 1999; Ebrahimi et al., 2009; Proulx-Curry and Chasteen, 1995; Xu and Chasteen, 1991; Yang et al., 1998). Given the lack of absolute conservation in structure of the ferroxidase center, it is likely that there is no single conserved mechanism for their activity (Baaghil et al., 2003; Chasteen and Harrison, 1999; Ebrahimi et al., 2009; Proulx-Curry and Chasteen, 1995; Xu and Chasteen, 1991; Yang et al., 1998). All of the ferritin family proteins that have been structurally characterized have additional iron-binding sites proximal to the ferroxidase center. In the case of the sites proximal to the ferroxidase center identified in DPS and ferritins, these have been suggested to be iron-entry, transit, and exit points from the ferroxidase center (Behera et al., 2015; Masuda et al., 2010; Theil et al., 2013), or the sites of electron transfer from Fe(II) substrates to the FOC (Yao et al., 2012). The subunit boundaries in the DPS and ferritin nanocages form a variety of symmetrical pores, which have variously been suggested or shown to be entry and exit sites to the lumen for iron, water and phosphate substrates (Andrews, 1998; Bellapadrona et al., 2009; Giessen et al., 2019; Liu et al., 2007; Treffry and Harrison, 1978). Whatever their function, these secondary metal sites are essential to the correct functioning of the ferritin iron-stores (Ebrahimi et al., 2016; W. Hagen et al., 2017; Treffry et al., 1998).

In our previous work on the EncFtn family we identified a number of conserved secondary metal-binding sites (He et al., 2019, 2016). A partially occupied metal ion site was present on the outer surface (proposed exit site) of the EncFtn decamer (Figure 1A in (He et al., 2016)), and a highly occupied metal ion site was present on the inner face (proposed entry site), which is proximal to the ferroxidase center (Figure 1B in (He et al., 2016) (Figure 2A and B). Both of these sites are found at dimer interfaces between the two-helix EncFtn subunits and are formed when the four-helical ferritin fold is reconstituted by the iron-mediated assembly of the EncFtn ferroxidase center (Ross et al., 2020). A comparison of these sites to secondary metal binding sites in other ferritin family members highlights the fact that ‘C’ site, first identified in the *E. coli* ferritin (Hempstead et al., 1994; Stillman et al., 2001), is topologically similar to the proposed iron-exit site identified in our previous work, whereas the entry site is unique to the EncFtn family (He et al., 2016). In our structure of the *R. rubrum* EncFtn protein, the entry site coordinates a calcium ion from the crystallization solution in a symmetrical arrangement of glutamic acid residues (E31/E34) and a tryptophan residue (W38) contributed by each chain from the ferroxidase center dimer (Ross et al., 2020) (Figure 2C). The metal coordination distances between the side chain carboxylic acid oxygens and calcium ion are 2.1 Å Ca – Glu31-Ox, 2.7 Å Glu34 – Ox, which are similar to the Fe – Glu – Ox coordination distances seen in the FOC of 2.2-2.6 Å.

**Figure 2.**
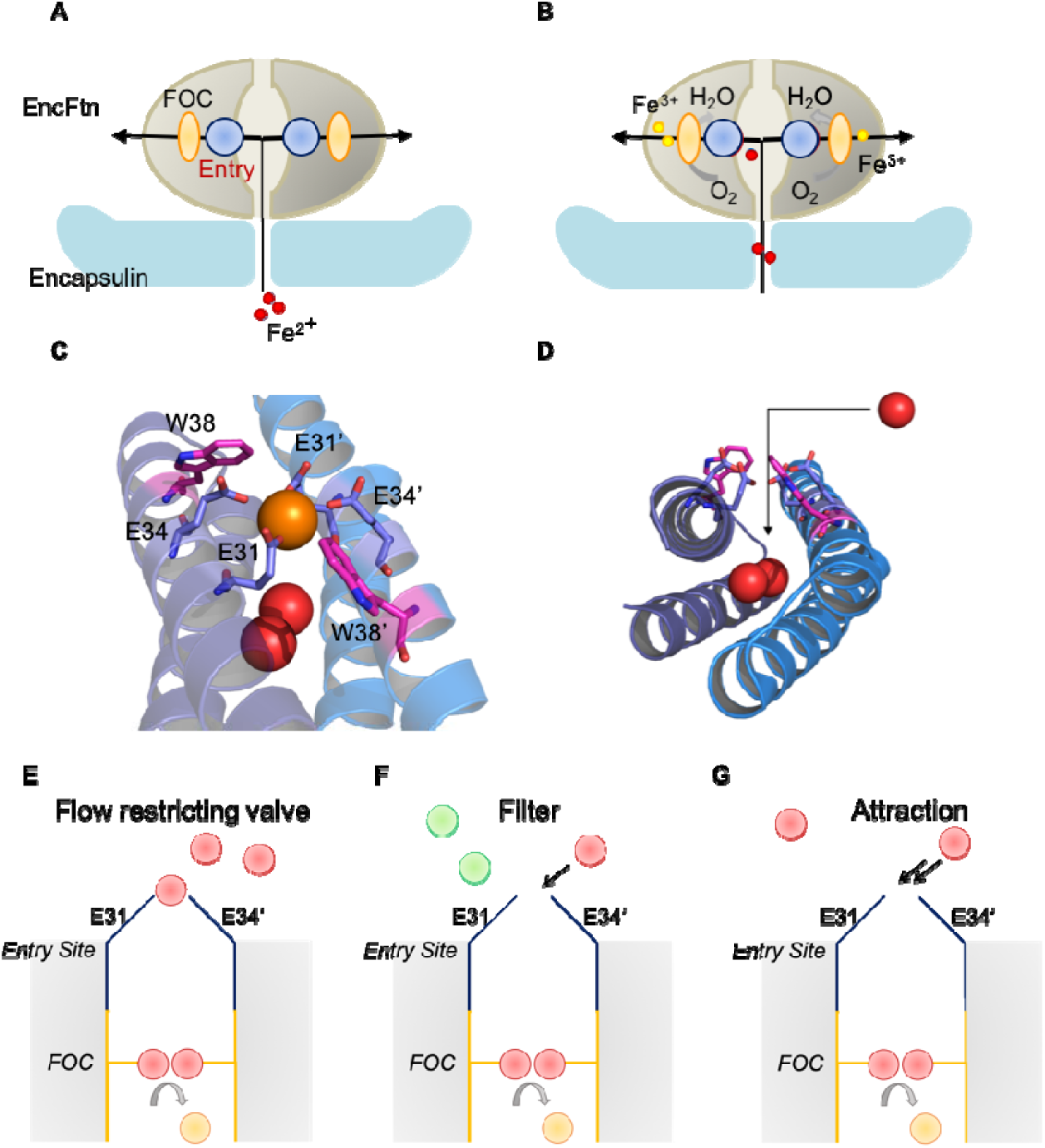
Hypotheses for the functional role of the entry site in EncFtn. (**A, B**) Schematic view of the proposed pathway for incoming ferrous ions moving towards the ferroxidase center (FOC). EncFtn is represented by a grey oval, where the entry sites and ferroxidase centers are shown in blue and yellow, respectively. A portion of the encapsulin is displayed in pale blue, with emphasis given to the 5-fold pore used by ferrous ions to reach the encapsulated ferritin. Ferrous ions (red circles) are shown to be converted into ferric ions (yellow circles) in the ferroxidase center with the concomitant reduction of molecular oxygen. (**C**) Coordination of calcium (orange sphere) within the dimer interface by Glu31 and Glu34 from the two subunits. Iron ions coordinated in the ferroxidase center (FOC) are depicted as red spheres. Trp38 side chains (pink) from both chains are also shown. (**D**) Structure of the proposed iron entry site. The black arrow indicates movement of ferrous ions (red spheres) pathway to FOC through the EncFtn dimer interface. Glu31, Glu34 and Trp38 side chains from both subunits are shown. Several hypotheses for the role played by the residues contributing to the entry site are shown in **E**-**G** (flow restriction valve (**E**), selectivity filter (**F**), and attraction (**G**). Ferrous and ferric ions are depicted as red and yellow circles, respectively, while non-cognate species are in green. The entry site and ferroxidase centrum locations are represented with blue and yellow lines, respectively.

We hypothesize that this site acts as an entry site for iron ions approaching the ferroxidase center, and acts in some way to control the access of metal ions to the active site of the protein (Figure 2D). The glutamic acid residues in the entry site are arranged in such a way as to capture and coordinate an Fe(II) ion in an octahedral geometry before passing it to the FOC. The positioning of W38 adjacent to this site suggests that the indole ring sterically constrains the position of the E31 side chain to an optimal position for metal coordination. We have considered several hypotheses regarding the functional role of the entry site, including acting as a flow restrictor valve to limit the flow of metal ions to the FOC (“valve” function); an electrostatic funnel to direct metal ions to the FOC (“attraction” function); a discriminatory filter to avoid mis-metalation and consequent inhibition of the active site (“filter” function) (Figure 2E/F/G); or for protein stability and assembly (“stabilizing” function). It is also possible that these functions may overlap, and the entry site may play a number of key roles in the stability and activity of the encapsulated ferritins.

In order to understand the role of the entry site, a number of single-point variants of the *R. rubrum* EncFtn protein were produced by site-directed mutagenesis of heterologous expression plasmids to replace the residue of interest with non-metal binding residues. These variants were produced in the isolated *R. rubrum* EncFtn protein alone and in an *R. rubrum* Encapsulin:EncFtn (Enc:EncFtn) co-expression system. We produced and purified recombinant E31A, E34A and W38G/A variants of EncFtn and the Enc:EncFtn complex and analyzed the solution and gas phase behavior of these proteins to understand the influence of these variants on the stability of the EncFtn complex. We obtained X-ray crystal structures of the E31A and E34A variants of the EncFtn protein, finding that their structures display altered entry site geometry, with the abrogation of metal binding in this site in the E34A variant. The functional role of the entry site was explored biochemically; we show that changes of metal coordination in this site enhance the ferroxidase activity of the EncFtn protein, while increasing its susceptibility to inhibition by competing zinc ions. Taken together, these results indicate that the entry site plays a minor role in stabilizing the oligomeric state of the EncFtn decamer, and has a dual role as a site of metal ion attraction and as a flow restrictor valve to limit metal access to the ferroxidase center and thus slow the oxidation of ferrous ions to ferric ions.

## Results

### Purification of variant encapsulated ferritins

In order to explore the functional roles of the residues in the entry site of the *R. rubrum* EncFtn, a number of single point variants were produced by site-directed mutagenesis, removing each metal binding residue in turn and substituting each with alanine or glycine. The resulting set of variants were also generated in association with the encapsulin (Enc) to produce Enc:EncFtn complexes. All proteins were produced by heterologous expression in *Escherichia coli.*

Our previous work demonstrated that changes to the ferroxidase center affect both the assembly and activity of the *R. rubrum* EncFtn (EncFtn-WT) (He et al., 2016). To investigate whether changes to the entry site influence the quaternary structure of the EncFtn protein, each variant was subjected to size-exclusion chromatography using a HiLoad 16/60 Superdex 200 (S200) column calibrated with standards of known molecular mass (Figure 3A and B). The average molecular masses of the EncFtn proteins were confirmed by liquid chromatography-mass spectrometry (LC-MS, Table 1). Each of the variant EncFtn proteins eluted with a similar profile, with a major peak at 60 ml retention volume, a second peak at 75 ml, and the final peak at 82 ml (Figure 3A, Figure 6-figure supplement 1A, Table 2). The E34A variant had significantly more protein in the 82 ml peak when compared to the EncFtn-WT and E31A variant, suggesting that this change affects the stability of the protein complex. Based on our prior work on the assembly pathway of the *Haliangium ochraceum* EncFtn protein (Ross et al., 2020) and native mass spectrometry of other representative members of the EncFtn family (He et al., 2019) these peaks would correspond to decamer, tetramer, and dimer species, respectively. The anomalous size estimation based on calibration standards can be attributed to the fact that the EncFtn proteins used in this study were full length proteins complete with their flexible encapsulation sequence, which would increase the hydrodynamic radius of the oligomers considerably.

**Table 1.** Average molecular masses of encapsulated ferritins obtained by LC-MS.

| Protein | Observed Mass (Da) | Assignment | Theoretical Mass (Da) |
| --- | --- | --- | --- |
| EncFtn-WT | 15060.71 ± 0.14 | monomer without Met | 15061.55 |
|  | 30120.78 ± 0.48 | dimer without Met | 30123.10 |
| EncFtn-E31A | 15003.78 ± 0.03 | monomer without Met | 15003.51 |
| EncFtn-E34A | 15003.29 ± 0.07 | monomer without Met | 15003.51 |
|  | 30006.20 ± 0.84 | dimer without Met | 30007.02 |
| EncFtn-W38A | 14946.09 ± 0.08 | monomer without Met | 14946.42 |
| EncFtn-W38G | 14932.33 ± 0.06 | monomer without Met | 14932.39 |
| Enc:EncFtn-WT | 29733.06 ± 0.16 | Enc monomer with Met | 29732.99 |
|  | 15061.59 ± 0.15 | EncFtn monomer without Met | 15061.55 |
|  | 30121.88 ± 0.78 | EncFtn dimer without Met | 30123.10 |
| Enc:EncFtn-E31A | 29732.42 ± 0.09 | Enc monomer with Met | 29732.99 |
|  | 15003.59 ± 0.14 | E31A monomer without Met | 15003.51 |
| Enc:EncFtn-E34A | 29732.80 ± 0.22 | Enc monomer with Met | 29732.99 |
|  | 15003.05 ± 0.25 | E34A monomer without Met | 15003.51 |
|  | 30005.48 ± 0.95 | E34A dimer without Met | 30007.02 |
| Enc:EncFtn-W38A | 29732.87 ± 0.23 | Enc monomer with Met | 29732.99 |
|  | 14946.25 ± 0.04 | W38A monomer without Met | 14946.42 |
| Enc:EncFtn-W38G | 29732.37 ± 0.46 | Enc monomer with Met | 29732.99 |
|  | 14932.23 ± 0.06 | W38G monomer without Met | 14932.39 |
| *ID*-EncFtn | 12404.67 ± 0.06 | monomer without Met | 12404.46 |

The starting Methionine residue is not always retained, and this has been indicated in “Assignment”. Error values generated from MassLynx v4.1.

**Table 2.** Elution volumes and oligomerization states of EncFtn and Enc:EncFtn from size-exclusion chromatography.

| Protein | Theoretical Mass without Met1 (Da) | Peak 1 |  | Peak 2 |  | Peak 3 |  |
| --- | --- | --- | --- | --- | --- | --- | --- |
| EncFtn | 15061.55 | $V_e =$<br>59.89 ml | 292 kDa,<br>19-mer | $V_e =$<br>75.68 ml | 81 kDa<br>5-mer | $V_e =$<br>82.38 ml | 47 kDa<br>3-mer |
| EncFtn-E31A | 15003.51 | $V_e =$<br>60.77 ml | 272 kDa<br>18-mer | $V_e =$<br>76.15 ml | 78 kDa<br>5-mer | $V_e =$<br>82.53 ml | 46 kDa<br>3-mer |
| EncFtn-E34A | 15003.51 | $V_e =$<br>59.42 ml | 304 kDa<br>20-mer | $V_e =$<br>76.01 ml | 79 kDa<br>5-mer | $V_e =$<br>81.63 ml | 50 kDa<br>3-mer |
| EncFtn-W38A | 14946.42 | $V_e =$<br>59.94 ml | 291 kDa<br>19-mer | $V_e =$<br>76.48 ml | 76 kDa<br>5-mer | | |
| EncFtn-W38G | 14932.39 | $V_e =$<br>60.07 ml | 288 kDa<br>19-mer | $V_e =$<br>75.45 ml | 82 kDa<br>5-mer | $V_e =$<br>82.59 ml | 46 kDa<br>3-mer |
| Enc-EncFtn | | $V_e =$<br>45.18 ml | 966 kDa | | | | |
| Enc:EncFtn-E31A | | $V_e =$<br>45.32 ml | 955 kDa | | | | |
| Enc:EncFtn-E34A | | $V_e =$<br>46.91 ml | 839 kDa | | | | |
| Enc:EncFtn-W38A | | $V_e =$<br>45.42 ml | 947 kDa | | | | |
| Enc:EncFtn-W38G | | $V_e =$<br>46.92 ml | 839 kDa | | | | |

**Figure 3.**
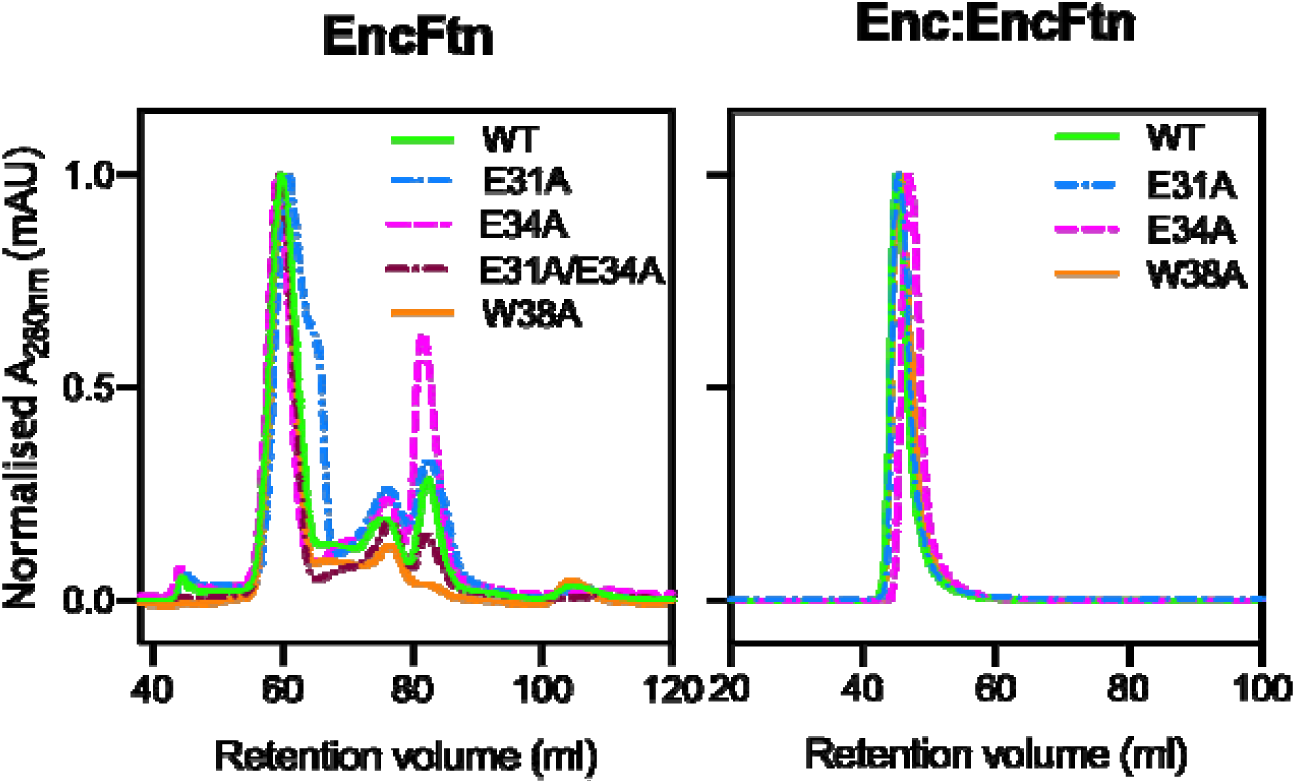
Purification of recombinant EncFtn and Enc:EncFtn protein complexes. Recombinant EncFtn (**A**) and Enc:EncFtn **(B)** proteins were purified by anion exchange chromatography (Hi-Trap Q-sepharose FF, GE Healthcare) and then subjected to size-exclusion chromatography using a Superdex 200 16/60 column (GE Healthcare) previously equilibrated with 50 mM Tris-HCl, pH 8.0, 150 mM NaCl. The elution profiles in **A** reveal that all EncFtn proteins show a main peak at around 60 ml, diagnostic of oligomerization states close to 10-mer, and smaller peaks at ∼ 76 ml and 82 ml (indicative of smaller assembly states, Table 2). Enc:EncFtn proteins elute in a single peak around 46 ml, suggesting that EncFtn is compartmentalized within the Encapsulin shell. An SDS-PAGE gel of (B) is shown in Figure 3–figure supplement 1, while transmission electron micrographs of Enc:EncFtn proteins are presented in Figure 3-supplement 2. Comparison of elution profiles from gel-filtration purification step of EncFtn-W38A and W38G are shown in Figure 3-figure supplement 3A/B. doi. 10.6084/m9.figshare.9885557

In contrast to the EncFtn variants, the Enc:EncFtn protein complexes all eluted as a single peak within the void volume of the S200 column, indicating the formation of protein complexes larger than 600,000 Da (separation range M_r_ for this column is 10,000-600,000 Da), hence not able to enter the matrix pores (Figure 3B, Figure 3-figure supplement 3B, Table 2). SDS-PAGE analysis of peak fractions confirmed the presence of both the EncFtn and encapsulin proteins (Figure 3–figure supplement 1). To determine if these complexes had formed correctly assembled encapsulin nanocages, we performed transmission electron microscopy on uranyl acetate-stained samples (Figure 3-figure supplement 2, Figure 3-figure supplement 3D/E). The encapsulins purified with wild type and variant EncFtn all showed characteristic 25 nm encapsulin cages, with visible internal density for the EncFtn proteins. This is consistent with our previous observations for the Enc:EncFtn wild-type complex (He et al., 2016).

### X-ray crystal structures of EncFtn variants

Given the observation that the E31A and E34A EncFtn variants do not display any significant differences in terms of oligomeric state and stability in solution, we subjected truncated variants of these proteins with the encapsulation sequence removed and a C-terminal hexa-histidine (EncFtn-sH) to crystallization screening to determine any changes at the metal binding sites. We obtained crystals of both the E31A and E34A variants in similar conditions to EncFtn-WT. We were unable to obtain crystals of either W38G or W38A variants of the EncFtn protein, suggesting that removing this tryptophan destabilizes the quaternary structure of the protein sufficiently to present a barrier to crystallization.

The structures of the *R. rubrum* EncFtn-sH E31A and E34A variants were determined by X-ray crystallography to 2.66 and 2.19 Å resolution respectively. They adopt the same overall quaternary fold as EncFtn-WT, with root-mean-square deviation (RMSD) Cα differences of 0.2 A^2^ over 91 Cα and 0.17 Å^2^ over 90 Cα for the E31A and E34A variants respectively, and an RMSD Cα of 0.23 Å^2^ over 90 Cα between the two variants. This indicates that these amino acid substitutions do not significantly alter the structure of the polypeptide backbone n the E31A and E34A variants. The position of side chains in the FOC and their metal coordination geometry are conserved for both variants, indicating that changes to the entry site do not fundamentally change the structure of the FOC.

The entry site of the E31A variant appears to coordinate a metal ion via the Glu34 side chain. Inspection of anomalous maps allowed us to assign this as a calcium ion; however, the coordination distances are much longer than in EncFtn-WT, at 3.8 Å compared to 2.8 Å (Figure 4, Figure 4 figure supplement 1). Loss of the coordinating Glu31 side chain ligand does not appear to cause gross structural rearrangements in this site, and the change in metal ion binding when compared to the Wt protein can be ascribed to the change in the coordination environment around this site. In contrast, the entry site of the E34A variant does not appear to coordinate any metal ions and the side chain of Glu31 is flipped away by 120° from this site towards Arg42 (Figure 4, Figure 4 figure supplement 2), and the Trp38 indole ring is also moved away from this site when compared to EncFtn-WT. Together, these local structural changes imparted by the E31A substitution greatly disrupt the metal coordination in this site when compated to the WT protein.

**Figure 4.**
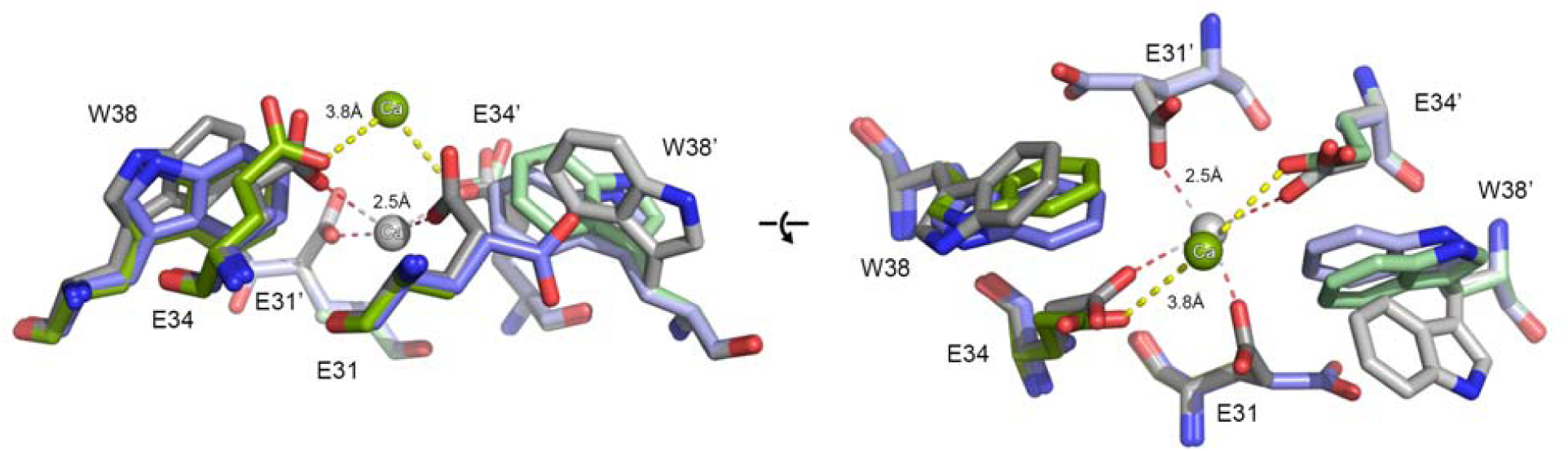
Comparison of metal ion binding in EncFtn entry site variant crystal. Metal ion entry sites for the X-ray crystal structures of the E31A and E34A EncFtn variants are shown with the EncFtn-sH wild-type for comparison. Residues in the entry site are shown as stick representations with spheres for coordinated calcium ions. Residues in the entry site are labelled, with residues from the second monomer indicated with prime symbols. WT is shown in grey, with the coordinated calcium ion in grey with coordination distances shown. E31A is shown in green with coordinated calcium ion in green; the metal ion is further from the site in this variant, with coordination distances to Glu34 of 3.8 Å. The E34A variant is shown in blue; no metal is coordinated in this site, and in the absence of a metal ion Glu31 is shifted by 117° when compared to the WT. Experimental electron density maps for the entry site residues of the E31A and E34 variants are shown in Figure-4-figure supplements 1 and 2 respectively.

### Native mass spectrometry of EncFtn variants

To build on the crystal structures of the EncFtn variants, we next used native mass spectrometry to study their structural dynamics and conformational heterogeneity(Ross et al., 2020). In our initial studies we found that EncFtn-W38A was not amenable to native mass spectrometry analysis; consequently, native MS was performed on EncFtn-WT, EncFtn-E31A, EncFtn-E34A, and EncFtn-W38G.

To interrogate the stability and oligomeric states of the EncFtn variants, we performed native MS and ion mobility MS experiments on samples purified by size-exclusion chromatography (Figure 5) (He et al., 2019). In native MS analysis, EncFtn-E31A behaves similarly to EncFtn-WT and presents solely as a decameric charge state distribution centered around the 25+ charge state (Figure 5, pink triangles). In contrast, under the same instrumental conditions, EncFtn-E34A and EncFtn-W38G displayed both decameric and monomeric charge state distributions (Figure 5, highlighted with pink triangles and blue circles, respectively). The presence of monomeric high charge states (8+ to 16+ for EncFtn-E34A, and 8+ to 15+ for EncFtn-W38G) suggests a degree of gas phase dissociation of the decameric complex during native MS analysis. Even with careful control of instrument conditions, this monomeric dissociation product could not be completely eliminated, suggesting that complexes of the EncFtn-E34A and EncFtn-W38G are less stable than EncFtn-WT. Thus, these initial observations suggest that loss of either E34 or W38 results in gas phase destabilization of the decamer assembly, although we cannot rule out the observed difference in stability being due to different iron content within the FOC in each of these protein preparations. We also note that the decameric charge state distributions of EncFtn-WT, EncFtn-E34A and EncFtn-W38G are extended and include several additional higher charge, low abundance charge states (Figure 5, highlighted with *) when compared to our previous EncFtn native MS analysis of a truncated EncFtn-WT protein (He et al., 2016). We attribute their presence to the ability of the solvent-exposed encapsulation sequence to readily protonate in solution. The difference in gas phase stability of the EncFtn variants was also evident when increasing the sampling cone voltage and/or activating the trap voltage of the mass spectrometer, which activates the complex and causes dissociation (Figure 5–figure supplement 1). EncFtn-W38G fully dissociates into its monomer charge state distribution at lower activation conditions than the other variants, further highlighting the instability of its decamer form (Figure 5-figure supplement 1D). The EncFtn-E31A, EncFtn-E34A and EncFtn-W38G variants (Figure 5-figure supplement 1, B, C and D) dissociate solely into their monomer charge state distribution with higher activation conditions, whereas a dimer species is also observed for EncFtn-WT; an observation which may indicate a weakened dimer interface in the variants.

**Figure 5.**
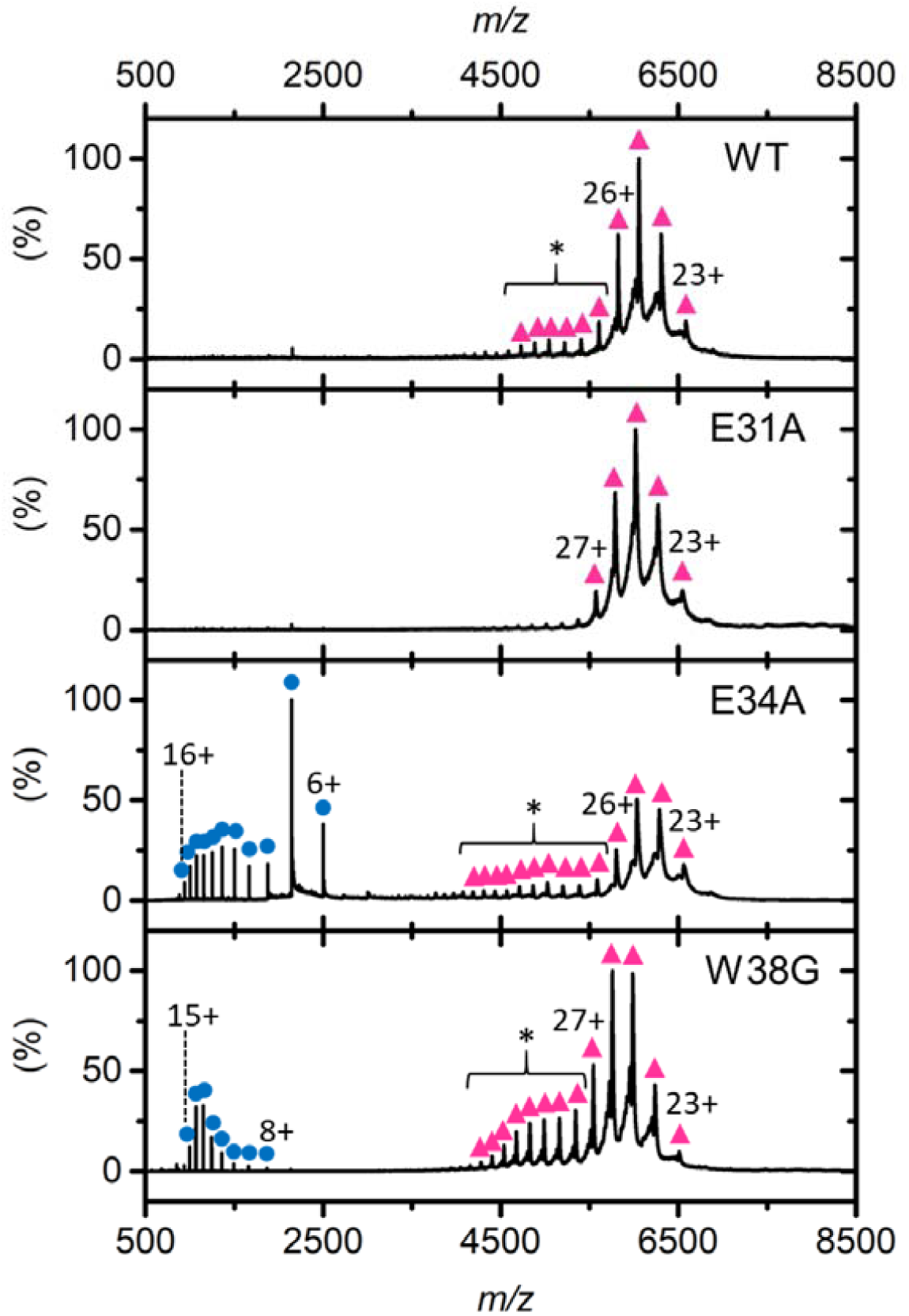
Native MS Analysis of EncFtn variants. Native nESI spectra of EncFtn variants displaying decameric (pink triangles) and monomeric (blue circles) charge state distributions. The elongated decameric charge state distributions of WT and E34A are highlighted by an asterisk. Gas phase dissociation of each variant is shown in Figure 7– figure supplement 1. Collision induced unfolding (CIU) MS experiments are presented as heat maps in Figure 7-figure supplement 2, with comparison of the relative ratio of compact to extended forms of each variant in Figure 7-figure supplement 3.

To further investigate the gas phase structure and stability of the variants, ion mobility collision induced unfolding (CIU) experiments were performed (Eschweiler et al., 2015). Each gas phase decamer protein complex was activated in a stepwise manner by incremental changes of the trap voltage prior to analysis of the conformation of the native protein ions using ion mobility mass spectrometry. The resulting ion mobility profile (arrival time distributions) for an individual protein ion is then plotted as a function of activation voltage to produce a heat map to reveal discrete conformations and gas phase unfolding transitions (Figure 5-figure supplement 2). It is clear from these experiments that two discrete conformations of each EncFtn variant are observed during CIU; a major conformation with a compact structure and a minor conformation with a more extended, unfolded structure (Figure 5-figure supplement 2 labelled ‘C’ and ‘E’ respectively, and Figure 5– figure supplement 3). At minimal activation, EncFtn-WT, EncFtn-E34A and EncFtn-E31A all exist in a single, compact decameric conformation with a drift time of between 10 and 11 ms, suggesting the same overall structure. As activation is increased, all three variants undergo a discrete transition to a more extended conformation (characterized by a drift time of 12.5 ms) at around 30 V activation, followed by a more complex transition to higher drift time conformations with activation above approximately 40V, before dissociation of the complex to release monomeric subunits. The close similarity of these CIU profiles strongly suggests similar gas phase structures and stability for the EncFtn-WT and the E31A and E34A variants (Figure 5-figure supplement 2 and Figure 5-figure supplement 3). In contrast, EncFtn-W38G occurs in the more extended conformation (12.5 ms) throughout the CIU experiment (Figure 5-figure supplement 2, W38G ‘E’), with the compact conformation only observed as a minor species at low activations. This is in accord with the native MS studies and suggests that although the W38G assembly shares similar overall structure to the WT and glutamate variants, the tryptophan substitution has a significant destabilizing effect on the decameric assembly. This increased structural/conformational heterogeneity in the tryptophan variant of EncFtn may account for our inability to obtain crystals in standard crystal screens.

By calibration of the ion mobility data, the collision cross sections (CCS) of both the discrete and extended conformations of the EncFtn variant decamers were determined (Table 3). The compact conformation of EncFtn-WT has a CCS of 71.40 nm^2^, whereas the more extended conformation, observed at higher activation, has a CCS of 80.62 nm^2^ (Table 3). We note that the CCS of the compact conformer is significantly larger than the CCS calculated from the EncFtn-sHis in the crystal structure of 58.7 nm^2^, and the 58.2 nm^2^ CCS observed in previously reported IM-MS experiments (He et al., 2016). However, prior studies were performed on truncated EncFtn-WT protein, which has the C-terminal encapsulation sequence removed. The higher observed CCS for EncFtn-WT suggests that the encapsulation sequence is on the surface of the decamer as it has an increased cross-sectional area. This is in accord with our observations in the size-exclusion experiments, where the EncFtn variants eluted with an apparently higher hydrodynamic radius than the expected oligomeric states. Thus, apart from the significant destabilization effect on higher order complex formation imparted by the tryptophan substitution, native MS and IM-MS experiments suggest that there is little overall structural change between the variants and EncFtn-WT.

**Table 3.**
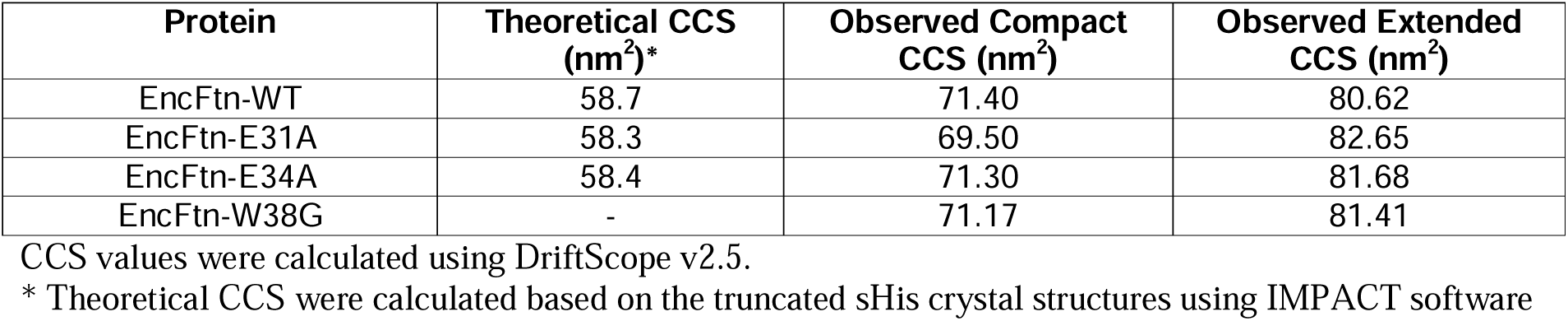
Collision cross sections (CCS) of the compact and extended conformations of the EncFtn variants.

| Protein | Theoretical CCS (nm<sup>2</sup>)* | Observed Compact CCS (nm<sup>2</sup>) | Observed Extended CCS (nm<sup>2</sup>) |
| --- | --- | --- | --- |
| EncFtn-WT | 58.7 | 71.40 | 80.62 |
| EncFtn-E31A | 58.3 | 69.50 | 82.65 |
| EncFtn-E34A | 58.4 | 71.30 | 81.68 |
| EncFtn-W38G | - | 71.17 | 81.41 |

CCS values were calculated using DriftScope v2.5.
\* Theoretical CCS were calculated based on the truncated sHis crystal structures using IMPACT software

### Ferroxidase activity of EncFtn variants

The changes in metal ion coordination observed in the entry sites in the crystal structures of the E31A and E34A variants suggest that these residues play a direct role in controlling the accessibility of the ferroxidase center to metal ions. To test our hypotheses about the functional role of this site (Figure 2), we investigated the ferroxidase activity of all of the EncFtn variants produced, both in isolation, and in complex with the encapsulin protein.

The ferroxidase activity of the wild-type EncFtn protein was confirmed to be comparable to that previously determined for the EncFtn-strepII variant (He et al., 2019). Surprisingly, all three variants displayed higher catalytic activity than the wild-type protein (Figure 6). The EncFtn-E34A variant had a 2-fold higher activity, while both the W38A and E31A mutations oxidize Fe(II) to Fe(III) around 5-fold faster, and their progress curves present a distinct shape in contrast to the wild-type protein (Figure 6). In order to test the contribution of Trp38 to the encapsulated ferritin catalytic activity we tested the W38A variant, allowing for a direct comparison with the other variants. We also verified that the W38G and W38A variants had comparable activities in complex with the Encapsulin protein (Figure 3-figure supplement 3C).

**Figure 6.**
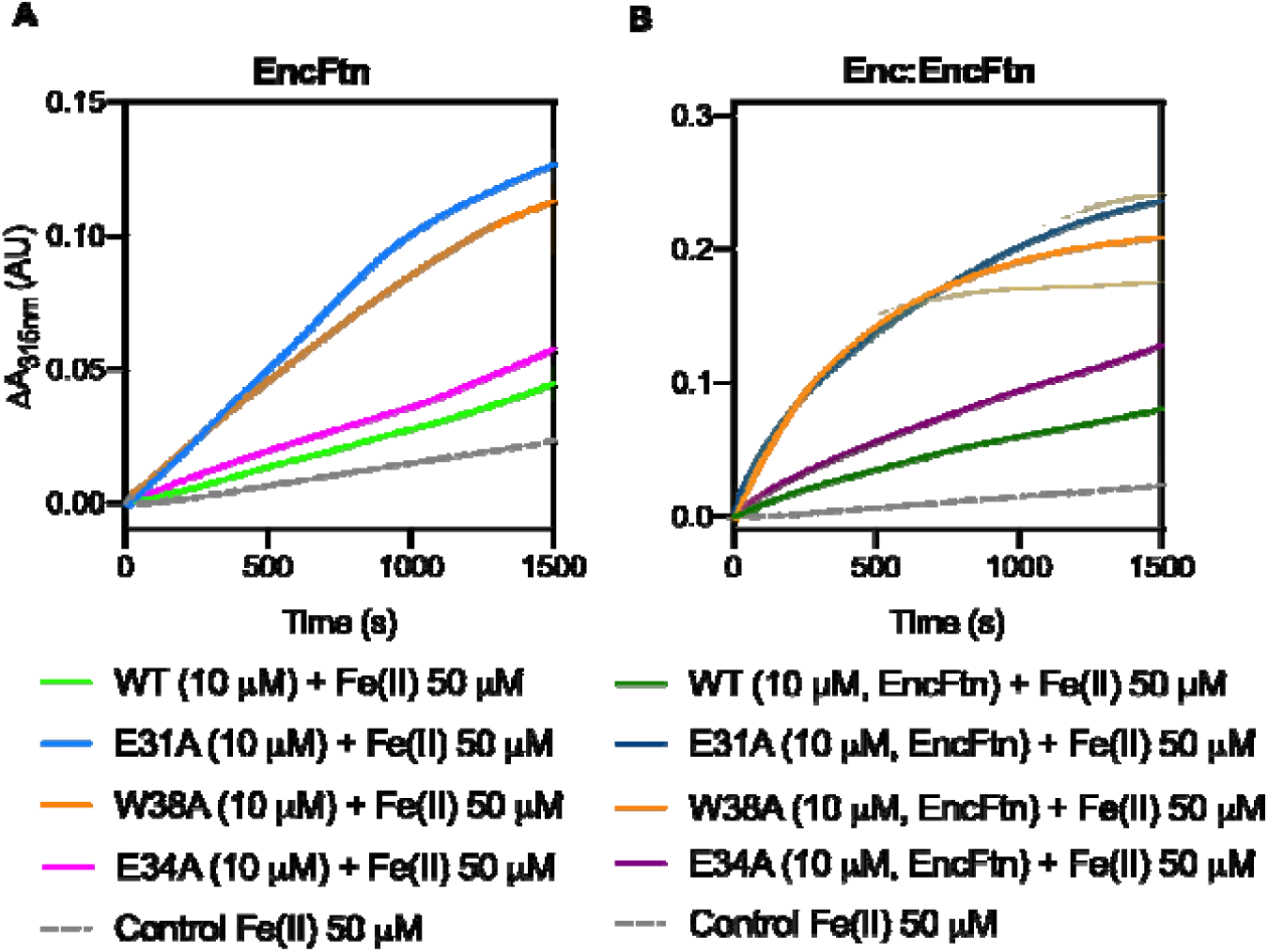
Ferroxidase activity of EncFtn and Enc:EncFtn complexes. (**A**) EncFtn wild-type (green line), EncFtn-E34A (pink line), EncFtn-E31A (blue line), and EncFtn-W38A (orange line), (10 μM, monomer) were incubated with 50 μM FeSO_4_.7H_2_O (10 times molar equivalent Fe(II) per FOC) and progress curves of the oxidation of Fe(II) to Fe(III) was monitored at 315 nm at room temperature. The background oxidation of iron at 50 μM in enzyme-free control is shown for reference (dotted grey line). Solid lines represent the average (n = 3) of technical replicates, shaded areas represent standard deviation from the mean. Protein and iron samples were prepared anaerobically in Buffer H (10 mM HEPES pH 8.0, 150 mM NaCl), and 0.1 % (v/v) HCl, respectively. (**B**) Enc:EncFtn-WT (dark green line), Enc:EncFtn-E34A (dark pink line), Enc:EncFtn-E31A (dark blue line), and Enc:EncFtn-W38A (orange line) (25 μM Enc:EncFtn, corresponding to 10 μM EncFtn monomer) were incubated with 50 μM FeSO_4_.7H_2_O and progress curves of the oxidation of Fe(II) to Fe(III) was monitored at 315 nm at room temperature. The background oxidation of iron at 50 μM in enzyme-free control is shown for reference (dotted grey line). Solid lines represent the average (n = 3) of technical replicates, shaded areas represent standard deviation from the mean. Protein and iron samples were prepared anaerobically in Buffer H (10 mM HEPES pH 8.0, 150 mM NaCl), and 0.1 % (v/v) HCl, respectively. Comparison of data collected for W38A and W38G variants are shown in Figure 3-figure supplement 3, including elution profiles from gel-filtration purification steps (A/B), ferroxidase activities of Enc:EncFtn-W38A and W38G (C) and Enc:EncFtn-W38A and W38G transmission electron micrographs (D/E). Data shown in A and B were re-plotted together in Figure 6-figure supplement 1 to emphasize the increase in activity when EncFtn are encapsulated. Linear regression on first 200s of ferroxidase assays with EncFtn and Enc:EncFtn proteins is shown in Figure 6-figure supplement 2, while Figure 6-figure supplement 3 presents comparison between EncFtn-E34A and E31A/E34A catalytic activities. doi.10.6084/m9.figshare.9885575

The increase in activity observed by all of these variants suggest that the entry site plays a role in restricting the flow of substrate Fe(II) ions to the catalytic FOC. This is supported by the structural data available for the E31A and E34A variants. When the differences in enzymatic activity are considered in the context of the X-ray crystal structures, it is clear that the disruption of metal coordination in the entry site has a significant impact on the catalytic activity of the EncFtn protein. The EncFtn-E31A structure shows that, as a consequence of changes in metal ion coordination, the restriction in metal ion flow provided by this variant is partially removed when compared to EncFtn-WT. However, the residual glutamic acid, Glu34, is still able to coordinate metal ions, albeit less closely than the wild type enzyme, given the increased distance between these residues and lack of additional coordination from Glu31 (Figure 4).

The crystal structure of the EncFtn-E34A variant shows that the mutation of Glu34 to Ala results in a reorganization of the entry site, with the side chain of Glu31 reoriented in such a way that it is no longer correctly positioned for metal coordination, essentially removing both the electrostatic attraction and flow restriction for this site (Figure 4). A ferroxidase assay carried out with an EncFtn-E31A/E34A variant shows that its activity is comparable to EncFtn-E34A (Figure 6-figure supplement 3). These data corroborate our hypothesis that in EncFtn-E34A, the flow restricting function is completely abolished, and no attraction is conveyed by the residual Glu31. As no flow restriction is provided at this level, we would expect to observe a higher activity, however the restructuring of this site results in a loss of any attracting forces, and consequently only a modest increase in activity is observed when compared to EncFtn-WT. The difference in activity between EncFtn-E31A and EncFtn-E34A shows how critical Glu34 is, and more broadly the importance of this electrostatic attraction, for the activity of the protein.

As we were unable to produce crystals of the W38A variant, we could not determine the effect of removing this tryptophan residue on the structure of the entry site. However, analysis of the crystal structure of the EncFtn-WT shows that this tryptophan’s indole ring appears to sterically constrain the Glu31 side chain in the correct position for metal-coordination (Figure 1, (He et al., 2016). We propose that the removal of Trp38 results in the Glu31 side chain being able to move away from this site. Our ferroxidase assay data show this variant as having a comparable activity profile to the E31A variant; we therefore propose that the two variants possess similar entry site structures. The broader destabilizing effects of the loss of the tryptophan side chain are harder to model, but the loss of a side chain of the size of the tryptophan indole may lead to further structural rearrangements around the internal dimer interfaces engaged by this residue.

The same ferroxidase assay was performed with the Enc:EncFtn complex, with wild-type and EncFtn variants (Figure 6B, Figure 6-figure supplement 1). The deviations from wild-type behavior were compared to those observed in the absence of the encapsulin nanocompartment (Figure 6A, Figure 6-figure supplement 1). Enzymatic reaction initial rates (*v*_0_) were calculated for each of the variants (in the first 200 s of the assay) and then divided by the initial rate of the corresponding wild-type protein (EncFtn-WT or Enc:EncFtn-WT) (Figure 6-figure supplement 1, Tables 4, 5 and 6). This allowed comparison of the assay data across the two contexts for the EncFtn enzyme (encapsulated vs non-encapsulated). Our data show that the E31A and W38A variants have a greater effect on activity within both systems (about 5-fold faster when compared to wild-type initial rate) while the E34A variant displays a 2-fold increase in the initial velocity *v*_0_ (Figure 6-figure supplement 3, Table 6). The EncFtn variants and wildtype retain the same relative differences in their activities whether they are associated to the encapsulin shell, or not. This observation suggests that the attraction and flow restricting properties are exerted by the entry site and are not changed by the sequestration of the EncFtn within the encapsulin cage.

**Table 4.** Linear regression fit of EncFtn ferroxidase assay progress curves.

|  | **EncFtn-WT** | **EncFtn-E31A** | **EncFtn-W38A** | **EncFtn-E34A** |
| --- | --- | --- | --- | --- |
| Equation | $Y = 2.322e-005 \cdot X - 0.0001751$ | $Y = 0.0001012 \cdot X - 0.001880$ | $Y = 9.463e-005 \cdot X + 0.0004715$ | $Y = 4.378e-005 \cdot X - 0.0002987$ |
| Slope $m$ | $2.322e-005 \pm 2.510e-007$ | $0.0001012 \pm 3.109e-007$ | $9.463e-005 \pm 1.575e-007$ | $4.378e-005 \pm 1.339e-007$ |
| R square | 0.9773 | 0.9981 | 0.9994 | 0.9981 |
| Number of X values | 201 | 201 | 201 | 201 |

**Table 5.** Linear regression fit data of Enc:EncFtn ferroxidase assay progress curves.

|  | **Enc:EncFtn-WT** | **Enc:EncFtn-E31A** | **Enc:EncFtn-W38A** | **Enc:EncFtn-E34A** |
| --- | --- | --- | --- | --- |
| Equation | $Y = 8.658e-005 \cdot X - 0.0004506$ | $Y = 0.0003718 \cdot X + 0.007998$ | $Y = 0.0004038 \cdot X - 0.0009754$ | $Y = 0.0001323 \cdot X + 0.002270$ |
| Slope $m$ | $8.658e-005 \pm 3.470e-007$ | $0.0003718 \pm 3.233e-006$ | $0.0004038 \pm 1.581e-006$ | $0.0001323 \pm 5.404e-007$ |
| R square | 0.9968 | 0.9852 | 0.997 | 0.9967 |
| Number of X values | 201 | 201 | 201 | 201 |

**Table 6.** EncFtn variants initial enzymatic rate ratios.

|  | **E31A** | **W38A** | **E34A** |
| --- | --- | --- | --- |
| EncFtn<br>$v_0^{\text{variant}} / v_0^{\text{wild-type}}$ | 4.36 | 4.08 | 1.89 |
| Enc:EncFtn<br>$v_0^{\text{variant}} / v_0^{\text{wild-type}}$ | 4.30 | 4.66 | 1.53 |

The initial enzymatic rates of the variants ( $v_0^{\text{variant}}$ ) were divided by the initial rate $v_0^{\text{wild-type}}$ . Initial rates were calculated on data from the first 200 s of ferroxidase assays (Tables 4 and 5).
The resulting values allow comparison between enzymatic kinetics of EncFtn and Enc:EncFtn proteins.
doi10.6084/m9.figshare.9885575

### Zinc inhibition of ferroxidase activity of EncFtn

In order to test the hypothesis that the entry site acts as a selectivity filter and is able to discriminate between metal ions approaching the FOC, ferroxidase assays were performed with the addition of zinc as a competitive inhibitor of ferroxidase activity (Le Brun et al., 1995). The activity of EncFtn wild-type and variant proteins (10 μM, monomer) with 50 μM Fe(II) was assayed in the presence of 34 μM Zn(II) (Figure 7A). The selected zinc concentration was previously determined to be the IC_50_ for wild type EncFtn-strepII (He et al., 2019). This concentration was chosen to allow identification of the impact of inhibition across the set of variant proteins. In line with our previous observations, EncFtn-WT showed 54% inhibition (He et al., 2019) (Figure 7A and B). The activity of the three variant proteins (E31, E34, W38) showed a markedly increased relative inhibition when compared to their activity in the absence of zinc (88%, 66%, and 69%, respectively) (Figures 7A and B). The increased inhibition seen in the variants can be explained by the changes in metal coordination at the entry site. With all the variants, the abrogation of metal coordination at this site permits greater access to the FOC for competing metal ions, as they are no longer effectively captured in the entry site. While these data show that the entry site restricts the general accessibility of the FOC to competing metal ions, they do not directly address the question of whether the site is acting as a selectivity filter.

**Figure 7.**
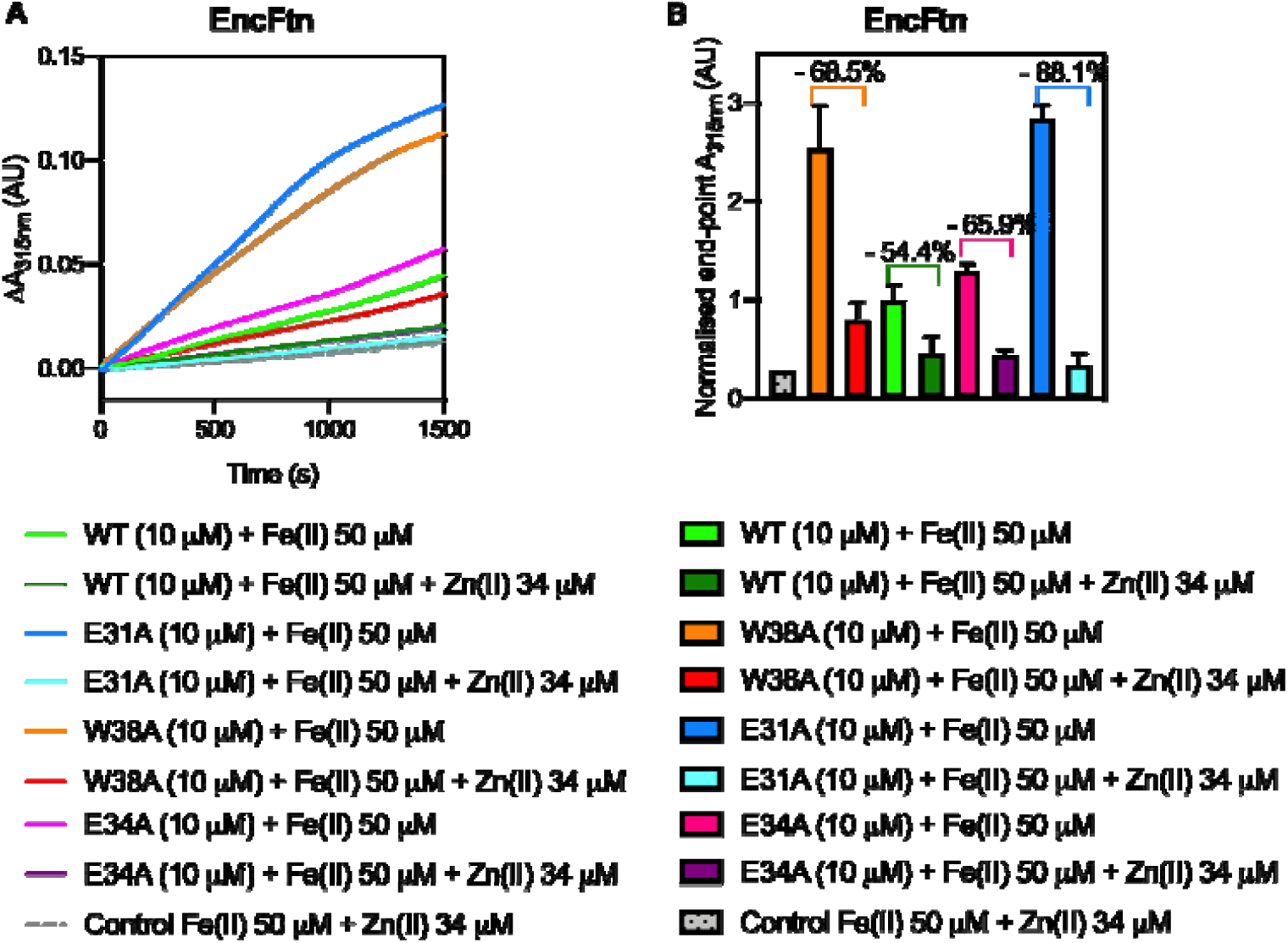
Ferroxidase activity of EncFtn wild-type and variants is inhibited by zinc. (**A**) Comparison between data shown in Figure 6A, collected in the absence of competing metals, are here plotted with curves from ferroxidase assays of EncFtn wild-type and variants carried out using [protein] = 10 µM and [Fe(II)] = 50 µM in the presence of [Zn(II)] = 34 µM. The zinc concentration corresponds to a response inhibited by 50% for EncFtn-Strep under the same experimental conditions (He et al., 2019). EncFtn wild-type is represented by a dark green line, EncFtn-E34A by a purple line, EncFtn-E31A by a pale blue line, and EncFtn-W38A by a red line. As a control, a mix of Fe(II) and Zn(II) salts (grey dotted line) were assayed in the absence of the enzyme. (**B**) End-point data presented in Figure 7A were plotted as columns to compare inhibition by zinc. Percentage of inhibition is shown above corresponding columns. Color-coding is consistent with Figure 7A. doi.10.6084/m9.figshare.9885575

### Steady state fluorescence emission of EncFtn in the presence of Zn(II)

To determine if zinc ions directly interact with the entry site and thus compete with iron for access to the FOC, intrinsic tryptophan fluorescence was exploited to study the Trp38 microenvironment in the presence and absence of zinc sulfate. Tryptophan fluorescence is the dominant source of protein intrinsic fluorescence with a maximum UV absorbance at ∼ 280 nm and emission at ∼ 320-350 nm (Teale and Weber, 1957). In addition to Trp38, EncFtn-WT possesses two more tryptophan residues (W72 and W80), one phenylalanine (F89), and two tyrosine residues (Y39 and Y87). Tyr39 is located in close proximity to the FOC and Glu31/Glu34, while the other aromatic residues are > 10 Å from both metal-coordination sites. Therefore, any observed change in the protein fluorescence emission should be due to changes in the entry site, and hence to metal binding at this site. This hypothesis was tested by monitoring steady state fluorescence emission of EncFtn proteins over time and upon addition of increasing sub-stoichiometric concentrations of Zn(II). The control experiment with EncFtn-W38A showed an hypsochromic shift of 8 nm in the emission maximum (λ_EM_ ∼ 322 nm compared to λ_EM_ ∼ 330 nm of EncFtn-WT) (Figure 8–figure supplement 1), suggesting a contribution to intrinsic fluorescence from more buried residues in the absence of Trp38. Although this variant still possesses both glutamic acid residues in the entry site, no specific trend was observed upon zinc addition (Figure 8, Figure 8–figure supplement 2).

**Figure 8.**
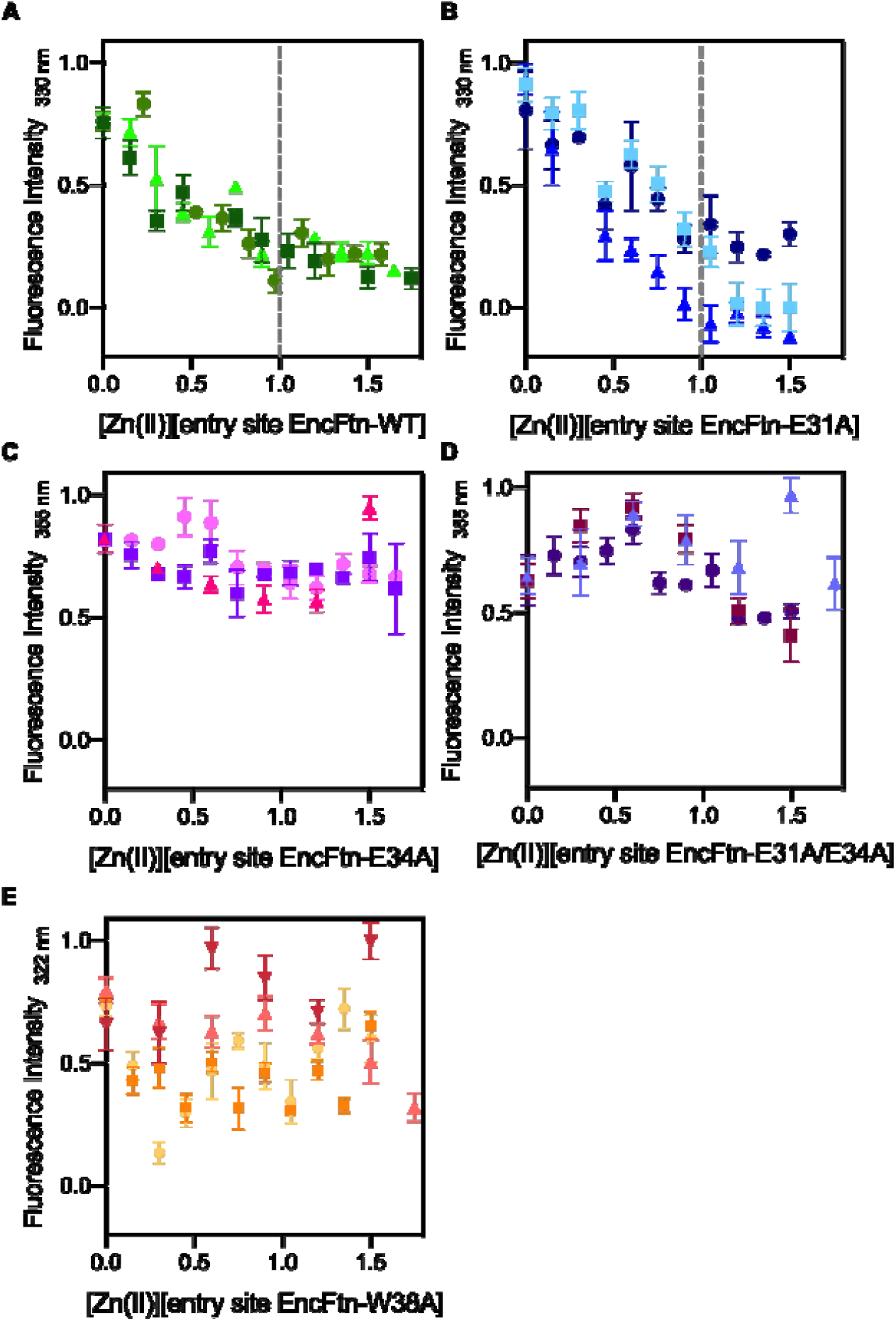
Intrinsic fluorescence of EncFtn wild-type and variants titrated with Zn(II). Fluorescence emission of EncFtn proteins (20 µM, corresponding to [entry-site] = 10 µM) at various wavelength (330 nm for EncFtn-WT and -E31A, 355 nm for EncFtn-E34A and E31A/E34A, and 322 nm for EncFtn-W38A) with excitation at 280 nm, and following titration with ZnSO_4_ .7H_2_O. (**A/B**) Fluorescence is quenched upon zinc addition, with inflection around 1 molar equivalents, suggesting perturbation of environment around Trp38. (**C/D**) In the absence of entry site residues available for metal coordination no quenching was observed. (**E**) EncFtn-W38A was tested as control. Symbols represent average and standard deviation of experimental replicates, collected at equilibrium either by Scan (n >3) or Kinetic (n = 10) options on the Cary Eclipse software package. Dotted grey lines were added at 1 molar equivalent of Zn(II) when inflection was observed. Intrinsic tryptophan fluorescence emission spectra in the absence of Zn(II) of wild-type and variants are shown in Figure 8-figure supplement 1 to display maximum emission values. Figure 8-figure supplement 2 shows fluorescence spectra in the presence and absence of 1.5 molar equivalents of Zn(II). doi.10.6084/m9.figshare.11920512

Only wild-type EncFtn and the E31A variant showed quenching of their fluorescence emission on addition of zinc, with an inflection at ∼1 molar equivalent of Zn(II) per entry site (Figure 8, Figure 8–figure supplement 2). Variants with abrogated metal coordination (E34A and E31A/E34A) did not show any specific trend, or inflection upon zinc addition (Figure 8, Figure 8–figure supplement 2). Moreover, tryptophan fluorescence is highly influenced by the solvent polarity of the surrounding environment and fluorescence emission spectra from both E34A and E31A/E34A variants exhibited a bathochromic shift of 25 nm (λ_EM_ ∼ 355 nm), implying an increasing polarity in Trp38 surroundings (Figure 8–figure supplement 1). These data suggest that upon metal coordination, the Trp38 indole ring is restrained towards the gateway of the channel leading to FOC.

These data suggest that metal ions can indeed interact with the entry site when it is intact (wild type), or partially present (E31A). In the absence of the entry site, Zn(II) will likely reach the FOC, an event that is not detectable with this assay, but shown by ferroxidase analysis in the presence of Zn(II) where increased inhibition of all variants was observed.

### Native FT-ICR mass spectrometry of ID-EncFtn

Given the observation in the intrinsic tryptophan fluorescence experiments that zinc directly interacts with the entry site, we performed high-resolution native FT-ICR MS to further probe whether this site was acting as a selectivity filter to restrict the passage of competing ions to the FOC. The high mass resolution of FT-ICR MS affords isotopic resolution of native protein complexes and, when applied to the study of metalloproteins, allows assignment of coordinated metals with high confidence. To further aid this analysis, we also employed an isotope depletion strategy, recently developed in our laboratory (Gallagher et al., 2020). The depletion of ^13^C and ^15^N stable isotopes from the EncFtn protein sample results in a dramatically reduced isotope distribution in mass spectrometry analysis. This simplified signal results in increased sensitivity and, most importantly for native MS analysis, reduces the occurrence of overlapping signals from proteoforms of similar mass, such as complete separation of sodiated and potassiated protein ions (a common observation in native protein MS). Thus, this strategy simplifies data interpretation and allows more confident and precise assignment of metal bound species. Due to the quadrupole range of the FT-ICR MS instrument used, a truncated and strep(II)-tagged version of EncFtn-WT was used as the depleted species, herein known as *ID*-EncFtn.

FT-ICR analysis of apo-*ID*-EncFtn reveals that it exists exclusively as a monomer (Figure 9A), as seen for EncFtn expressed in minimal media (He et al., 2016). Isotopic resolution confirms there is no metal bound within the purified *ID*-EncFtn monomer; which is as predicted, due to the limited bioavailability of metals in the minimal media required for the production of isotopically depleted proteins (Figure 9–figure supplement 1). Titration with Fe(II) prior to MS analysis results in the disappearance of the monomeric species and the appearance of a decameric charge state distribution (Figure 9B, pink triangles). This result is consistent with the Fe(II)-dependent assembly of EncFtn (He et al., 2016). Interestingly, titration with Zn(II) prior to native MS reveals a similar result with decamer being formed (Figure 14C, pink triangles), supporting our previous solution-phase observations that metal-mediated higher order assembly is not specific for iron (He et al., 2016). Loading *ID*-EncFtn with Fe(II) followed by the addition of Zn(II) results in a decameric charge distribution similar to that of the Zn(II)-mediated decamer (Figure 9D). Isotopic resolution cannot be achieved for the decameric species due to heterogenous metal loading, but there is sufficient mass resolving power to allow approximate assignment of metal-loading (Figure 9–figure supplement 2). Titrations with Fe(II) reveal a decameric complex consistent with between 10 and 20 Fe(II) ions bound (Figure 9–figure supplement 2A). This is more than seen previously (He et al., 2016), but could be due to coordination of additional surface iron ions, or other metal adducts (such as sodium or potassium, which are commonly observed in native MS). There are between 10 and 30 Zn(II) ions associated with the decameric species during Zn(II) titrations. This suggests that Zn(II) is coordinating to *ID*-EncFtn in more of the metal binding sites than iron (Figure 9–figure supplement 2B). The decameric *ID*-EncFtn species observed after the addition of both iron and zinc has a higher mass than either individually (Figure 9–figure supplement 2C) - including the upper range of 30 Zn(II) ions seen in zinc titrations - suggesting both metals may be associated to the complex simultaneously. No peak is observed corresponding to 10 Fe(II) ions (iron-loaded FOC) and 5 Zn(II) ions (Zn(II)-loaded entry site).

**Figure 9.**
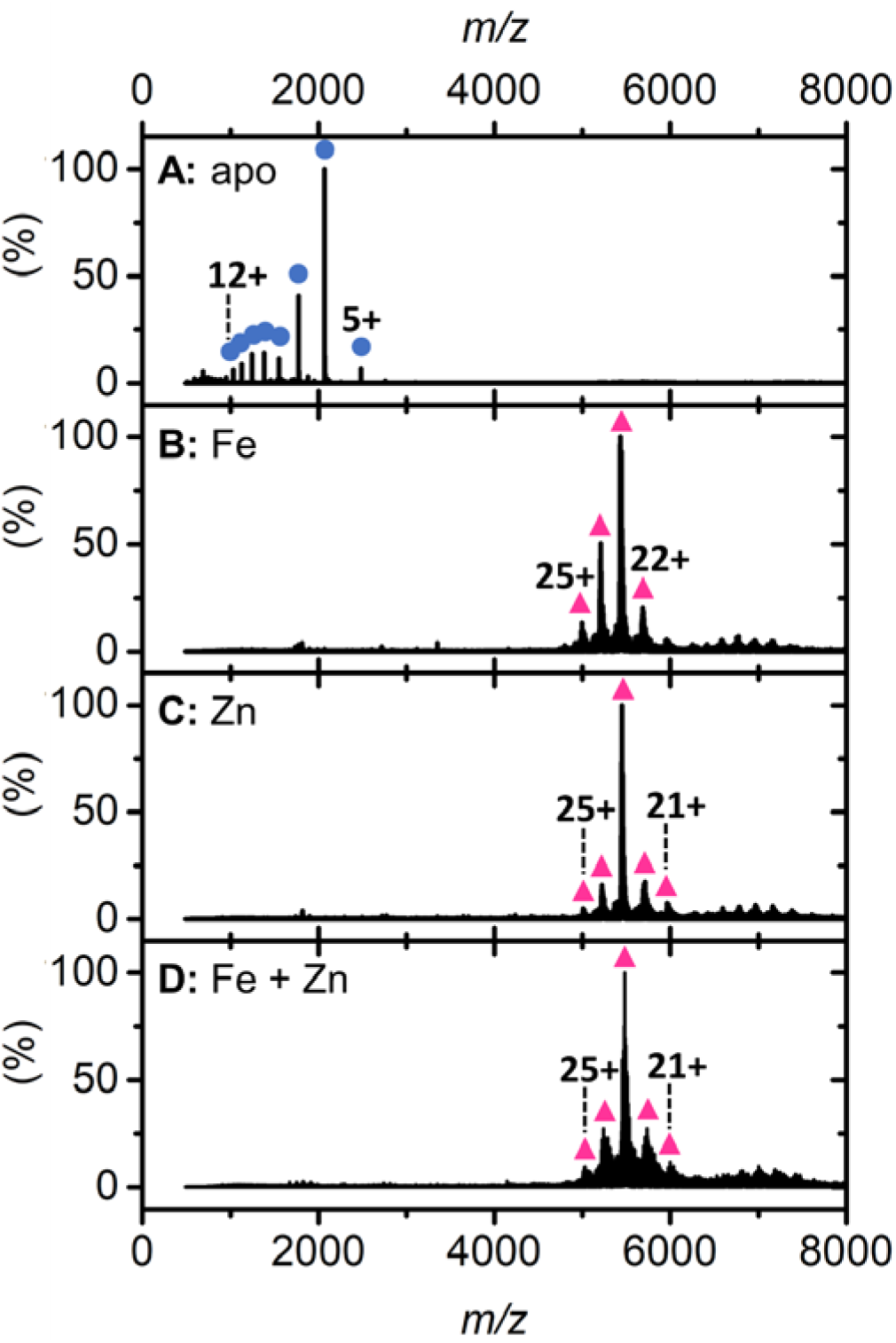
Native FT-ICR mass spectrum of *ID*-EncFtn. Native nESI mass spectra of: apo-*ID*-EncFtn (**A**); *ID*-EncFtn titrated with Fe(II) (**B**); *ID*-EncFtn titrated with Zn(II) (**C**); and *ID*-EncFtn titrated with Fe(II) followed by Zn(II) (**D**). Oligomerization states are stressed with colored shapes, monomer in blue circles and decamer in pink triangles. Metal loading of (**A**) is shown in Figure 9–figure supplement 1, and metal association of **B**-**D** is shown in Figure 9–figure supplement 2. B, C and D experience different dissociation pathways during CID shown in Figure 9–figure supplement 3, Figure 9–figure supplement 4 and Figure 9– figure supplement 5 respectively.

We used collision-induced dissociation (CID) to gain additional understanding of metal loading in *ID*-EncFtn. Under the correct experimental conditions, CID can lead to dissociation of a protein complex whilst retaining ligand interactions and specific subcomplex assembles, and thus can provide information on the topology of the original complex. Iron-loaded *ID*-EncFtn dissociates predominantly into a high-charge monomer species, the so-called ‘typical’ dissociation product of protein complexes (Hall et al., 2013). However, upon closer inspection, evidence of higher order collision induced subcomplexes is present - a minor tetramer species is observed at *m/z* 3552 (charge 14+) (Figure 9-figure supplement 3A, blue circles and purple diamond respectively). This species has low signal-to-noise ratio (S/N); however, its depleted isotope distribution is similar to that of four *ID*-EncFtn monomers with two Fe(III) ions bound (Figure 9-figure supplement 3B). We assign this species as of two non-FOC dimers bound together by a di-iron containing FOC, this dissociation pathway is consistent with the proposed assembly of EncFtn (Ross et al., 2020) (Figure 9-figure supplement 3C).

In stark contrast, *ID*-EncFtn with coordinated Zn(II) ions dissociates into monomer and dimer species (Figure 9–figure supplement 4A, blue circles and green squares respectively). Isotopic resolution was achieved for the dimer species and reveals this species has a mass consistent with two *ID*-EncFtn monomers associated two Zn(II) ions (Figure 9–figure supplement 4B). The presence of two metal ions bound within the dimer suggests that this species is the FOC dimer with the FOC occupied by two Zn(II) ions. There is no peak observed consistent with an apo-dimer, nor a dimer with one, or three, Zn(II) ions bound. This ‘atypical’ dissociation pathway was previously observed for EncFtn (He et al., 2016) (Figure 9–figure supplement 4C). The differing dissociation pathways of zinc-coordinated *ID*-EncFtn and iron-coordinated *ID*-EncFtn suggests that zinc stabilizes the FOC interface and binds with a higher affinity than iron in the FOC catalytic center; this higher affinity is as predicted from the Irving-Williams series. This supports our observations from the ferroxidase assay that show zinc inhibiting the FOC. Analysis of *ID*-EncFtn first loaded with iron and then subsequently challenged with zinc reveals a similar dissociation to that titrated solely with zinc (Figure 9–figure supplement 5A). The dimer species observed is *ID*-EncFtn_2_Zn(II)_2_, the same dimer species seen when titrated with zinc alone (Figure 9–figure supplement 5B). This suggests that in EncFtn-WT, zinc can efficiently displace iron in the FOC, that zinc inhibition acts at the catalytic ferroxidase center, and that zinc is not held within or discriminated against at the entry site (Figure 9–figure supplement 5C).

In summary, we have shown that substitution of residues in the entry site triad (E31, E34, W38) with non-metal binding residues results in the enhancement of enzymatic activity (Figure 6A). Our crystal structures support the conclusion that this is a consequence of the creation of a wider and more loosely coordinating entry site, which allows the faster passage of metal ions to the FOC due to a reduced energy barrier at this site (Figure 4). Furthermore, when these key residues are removed, the protein is more susceptible to inhibition by Zn(II).

## Discussion

Our work presents a comprehensive structure/function study investigating the role of the entry site proximal to the ferroxidase center (FOC) in encapsulated ferritin-like proteins. To investigate the role of this site we produced variant proteins with amino-acid substitutions in residues in this site to replace metal coordinating residues in the *R. rubrum* EncFtn protein with alanine (E31A, E34A) and to remove the large tryptophan side chain proximal to this site (W38A/G).

Solution and gas-phase analyses of variant proteins showed that, in contrast to the FOC, removal of the metal coordinating ligands (E31A/E34A) did not have a significant effect on protein oligomerization; however, removal of the Trp38 side chain had a significant destabilizing effect on the quaternary structure of the protein. X-ray crystal structures of the E31A and E34A variants highlighted changes in metal coordination at the entry site. The effect of these changes on the activity of the protein was tested using ferroxidase assays and zinc inhibition experiments. Removal of the Glu31 and Trp38 side chains significantly enhanced the activity of the enzyme; while removal of metal coordinating residues increased the susceptibility of the protein to inhibition by zinc. Investigation of the EncFtn intrinsic fluorescence emission properties upon zinc binding revealed coordination in the entry site when this is intact or partially formed (WT and E31A, respectively), confirming that this site acts as an additional metal binding site prior to the FOC.

High resolution MS analysis of *ID*-EncFtn metal binding reveals that *ID*-EncFtn decamer assembly can be induced by iron and zinc ions. Furthermore, zinc appears to bind within the FOC more tightly than iron and stabilizes the FOC dimer. When *ID*-EncFtn is titrated with iron and subsequently zinc, only zinc is observed bound in *ID*-EncFtn subcomplexes, indicating that zinc displaces iron from the FOC.

Taken together, these results imply that the entry site has a lower affinity for metal ions than the ferroxidase center and that in accord with the Irving-Williams series the FOC forms more stable complexes with zinc than iron. However, bacterial cells buffer metal concentrations via a polydisperse buffer in the order of the Irving-William series (Foster et al., 2014; Frausto da Silva and Williams, 1991; Reyes-Caballero et al., 2011; Tottey et al., 2008), resulting in highly competitive Zn(II) ions being buffered to lower available levels than Fe(II), thus presumably preventing the mis-metalation of ferritin sites *in vivo*.

These observations support the hypothesis that the entry site has a dual function, in attracting metal ions to the catalytic site, and as a flow restrictor to limit the movement of metal ions to the FOC and thus slow the rate of iron oxidation (Figure 2 E/G). Our results confirm that the ferroxidase center is indeed the active site of this enzyme, as the loss of the entry site enhances enzyme activity rather than abrogates it. Why the EncFtn protein has evolved a mechanism to effectively slow its activity is still an open question. There may be a trade-off between substrate turnover and the availability of a sink for the electrons liberated from the iron in the ferroxidase center. The catalytic mechanism and electron sink for ferritins is still a subject of vigorous debate. The EncFtn family presents a unique structural arrangement of the active site, which will no doubt present a distinct mechanism to other members of the ferritin family when probed further (W. R. Hagen et al., 2017). The entry site may well provide a brake to ensure productive coupling of iron oxidation and mineralization.

Encapsulation of the EncFtn protein within the encapsulin nanocage enhances the activity of all of the EncFtn variants (Figure 6), but does not change the way in which modifications to the entry site affect the catalytic activity, indicating that the function of the entry site is not changed through association with the encapsulin shell. These results suggest that the encapsulin shell acts to guide productive iron ion interactions with the EncFtn protein; indeed, our previous work showed that the encapsulin shell interacts with a significant amount of iron in the absence of the EncFtn protein. This warrants future investigation into the ability of the encapsulin shell to conduct metal ions to the EncFtn active site and the role of the encapsulin shell pores in the access of substrates to the encapsulated enzyme.

## Materials and Methods

| **Key Resources Table** |  |  |  |  |
| --- | --- | --- | --- | --- |
| **Reagent type (species) or resource** | **Designation** | **Source or reference** | **Identifiers** | **Additional information** |
| strain, strain background (*Escherichia coli*) | BL21(DE3) | Cambridge Biosciences | 60401 | chemically competent |
| recombinant DNA reagent | pACYCDuet-1 | Novagen | 71147 | vector for protein expression in *E. coli* |
| sequence-based reagent | Primers used in this study | IDT |  | See Table 8 |
| commercial assay or kit | Pierce™ BCA Assay kit | Thermo Fisher Scientific | 23225 |  |
| commercial assay or kit | Stratagene QuikChange kit | Agilent | 200523 |  |
| chemical compound, drug | Lysozyme | Sigma-Aldrich | L6876 |  |
| chemical compound, drug | Benzonase | Sigma-Aldrich | E1014 |  |
| chemical compound, drug | PEG 3350 | Sigma-Aldrich | 1546547 |  |
| chemical compound, drug | Calcium acetate | Sigma-Aldrich | 21056 |  |
| chemical compound, drug | Iron sulfate | Sigma-Aldrich | 1270355 |  |
| chemical compound, drug | PEG 200 | Sigma-Aldrich | 807483 | Cryoprotectant |
| chemical compound, drug | FeSO<sub>4</sub>·7H<sub>2</sub>O; Fe(II) in “ferroxidase assay” sections | Sigma-Aldrich | 1270355 |  |
| chemical compound, drug | ZnSO<sub>4</sub>·7H<sub>2</sub>O; Zn(II) in “zinc inhibition of ferroxidase assays” and “steady state fluorescence” sections | Sigma-Aldrich | 204986 |  |
| chemical compound, drug | <sup>14</sup>N (99.99%)-ammonium sulfate | ISOTEC | 485411 |  |
| chemical compound, drug | <sup>12</sup>C (99.9%)–glucose | Cambridge Isotope Laboratories | CLM-4819-1 |  |
| chemical compound, drug | Iron(II) chloride tetrahydrate; Fe(II) in the “Native FT-ICR mass spectrometry” section | Sigma-Aldrich | F2130 |  |
| chemical compound, drug | Zinc acetate; Zn(II) in the “Native FT-ICR mass spectrometry” section | Sigma-Aldrich | 383317 |  |
| software, algorithm | MassLynx | Waters Corporation |  |  |
| software, algorithm | DriftScope | Waters Corporation |  |  |
| software, algorithm | Spyder CIUsuite | Dixit et al., 2018 |  |  |
| software, algorithm | IMPACT | Marklund et al., 2015 |  |  |
| software,<br>algorithm | Data Analysis | Bruker<br>Daltonics |  |  |
| software,<br>algorithm | Isotope Pattern | Bruker<br>Daltonics |  |  |
| software,<br>algorithm | Prism 8 | GraphPad |  |  |
| software,<br>algorithm | Origin 2016 | OriginLab |  |  |
| software,<br>algorithm | Open Source PyMOL | Schroedinger<br>LLC |  |  |
| software,<br>algorithm | Phaser | McCoy et al.,<br>2007 |  |  |
| software,<br>algorithm | Phenix.autobuild | Adams et al.,<br>2010 |  |  |
| software,<br>algorithm | Phenix.refine | Afonine et al.,<br>2012 |  |  |
| software,<br>algorithm | Coot | Emsley et al.,<br>2010 |  |  |
| software,<br>algorithm | MolProbity | Chen et al.,<br>2010 |  |  |
| other | HiPrep Q HP; anion<br>exchange column | GE Healthcare | GE29-0181-82 |  |
| other | HiLoad 16/60 Superdex<br>200; size exclusion column;<br>S200 | GE Healthcare | GE28-9893-35 |  |
| other | HisTrap column | GE Healthcare | GE17-5255-01 | 5 mL column |
| other | Strep-Trap HP column | GE Healthcare | GE28-9075-48 | 5 mL column |

### Cloning

Expression constructs for the *R. rubrum* EncFtn protein (pACYCDuet-1-Rru_EncFtn) and Enc:EncFtn protein complex (pACYCDuet-1-Enc:EncFtn) were produced in the previous work (He et al., 2016) Site-directed mutagenesis was performed using Stratagene QuikChange kit following manufacturer instructions. Protein nomenclature used throughout the text is shown in Table 7. Primers used in this work for this purpose are listed in Table 8

**Table 7.** Protein nomenclature.

| Protein | Alternative nomenclature |
| --- | --- |
| Wild type EncFtn | EncFtn-WT |
|  | WT |
| E31A variant of EncFtn-WT | EncFtn-E31A |
|  | E31A |
| E34A variant of EncFtn-WT | EncFtn-E34A |
|  | E34A |
| W38A variant of EncFtn-WT | EncFtn-W38A |
|  | W38A |
| W38G variant of EncFtn-WT | EncFtn-W38G |
|  | W38G |
| E31A and E34A double variant of EncFtn-WT | EncFtn-E31A/E34A |
|  | E31A/E34A |
| Wild type Enc and EncFtn complex | Enc:EncFtn-WT |
|  | WT |
| Wild type Enc and E31A variant of EncFtn complex | Enc:EncFtn-E31A |
|  | E31A |
| Wild type Enc and E34A variant of EncFtn complex | Enc:EncFtn-E34A |
|  | E34A |
| Wild type Enc and W38A variant of EncFtn complex | Enc:EncFtn-W38A |
|  | W38A |
| Wild type Enc and W38G variant of EncFtn complex | Enc:EncFtn-W38G |
|  | W38G |
| His-tagged version | -sH |
| StreptII-tagged version | -strep |
| Isotopically depleted version | ID- |

Nomenclature adopted throughout this work to refer to the same species. The above is valid for each variant used in the study. When referring to tagged versions, the protein name acquires the specific suffix.

**Table 8.**
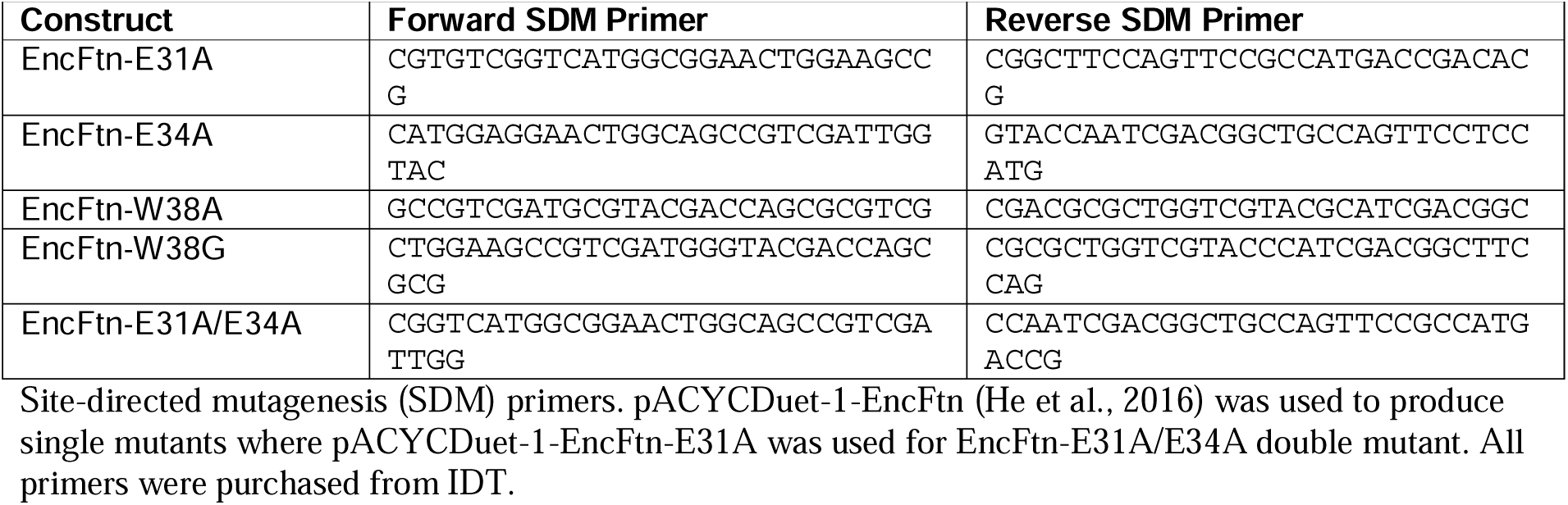
Primers used in this study.

### Protein production and purification

Expression plasmids for the *R. rubrum* EncFtn protein and Enc:EncFtn complex were transformed into competent *Escherichia coli* BL21(DE3) cells and grown in LB medium supplemented with appropriate antibiotics, and expressed as described previously (He et al., 2019, 2016). Protein sequences are listed in Table 9.

EncFtn and its variants, were purified as described in (He et al., 2019). Briefly, the proteins were purified by combining anion exchange chromatography (AXC) and size exclusion chromatography (SEC). Purity of protein samples was analyzed by 15 % SDS-PAGE.

Enc:EncFtn wild-type and variant complex cell pellets were resuspended in 10-times (v/w) of Lysis Buffer (20 mM HEPES pH 8.0, 2 mM MgCl_2_, 1 mg/ml lysozyme, 20 units/ml benzonase). Cells were sonicated on ice (5 minutes at 30 seconds on/off cycles, 35 % amplitude), followed by incubation for 30 minutes at 37 °C in a water bath. Cell lysate was clarified by centrifugation (35,000 x g, 30 minutes, 4 °C) and the supernatant was collected and incubated for 10 minutes at 80 °C in a water bath. After cooling the sample was centrifuged again (30 minutes, 35,000 x g, 4 °C), and the supernatant was collected and filtered using a 0.22 μm syringe filter (Millipore, UK). Enc:EncFtn protein complex sample was loaded onto a 20 ml anion exchange column (Hi-Prep Q HP, GE Healthcare) pre-equilibrated with Buffer QA (50 mM Tris, pH 8.0). Un-bound protein was washed with Buffer QA. A step gradient elution of 20 column volumes was performed by mixing Buffers QA and QB (50 mM Tris, pH 8.0, 1 M NaCl) to 100 % QB. Since Enc:EncFtn does not bind the anion exchange column, proteins of interest were found in flow-through sample, as confirmed by 15 % SDS-PAGE. Protein was concentrated using a Vivaspin centrifugal concentrator (MWCO 10 kDa, Sartorius) following the manufacturer’s instructions. Concentrated protein was then applied to a size exclusion column (HiLoad 16/60 Superdex 200, GE Healthcare) pre-equilibrated with Buffer GF (50 mM Tris, pH 8.0, 150 mM NaCl). Fractions of interest were run on a 15% SDS-PAGE to assess protein purity and oligomeric state, and further concentrated using a Vivaspin centrifugal concentrator as before. After concentration, protein samples were analyzed by 10% Tricine-SDS-PAGE (Schägger, 2006) in order to resolve bands around 30 kDa. Elution volumes of EncFtn and Enc:EncFtn complexes from the size exclusion column are listed in Table 2.

### His-tagged protein production and purification

Hexahistidine tagged EncFtn-E31A and -E34A variants (EncFtn-sH) were produced in *E. coli* BL21(DE3) as described above and purified following the same protocols as previously described (He et al., 2016). Briefly, a 5 ml HisTrap column (GE Healthcare, UK) was used for immobilized metal affinity chromatography (IMAC), followed by size exclusion chromatography (HiLoad 16/60 Superdex 200, GE Healthcare) using the protocol described above.

### Isotopically depleted strepII-tagged protein production and purification

To produce isotopically depleted EncFtn proteins, competent *E. coli* BL21 (DE3) cells were transformed with the required expression plasmid. A single colony was used to inoculate 10 ml of LB media supplemented with the appropriate antibiotic, before overnight incubation at 37 °C. This was then used to inoculate 500 ml of 2x YT media, incubated at 37 °C until an OD_600_ 0.6-0.8 was obtained, at which point the cell culture was centrifuged at 5000 x g at 4 °C for 20 minutes (200 ml per expression culture required). The pellet was washed with M9 salts solution (33.9 g/L Na_2_HPO_4_, 15 g/L KH_2_PO_4_, 2.5 g/L NaCl) and suspended in 100 ml M9 minimal media. Isotopically depleted M9 minimal media was supplemented with ^12^C (99.9%)–glucose (Cambridge Isotope Laboratories) and ^14^N (99.99%)-ammonium sulfate. The M9 minimal media cultures were further incubated at 37 °C for 1 hour, before protein expression was induced and stored as previously described (Gallagher et al., 2020).

Isotopically depleted strepII-tagged EncFtn (*ID*-EncFtn) was purified as reported in (He et al. 2019). Cells were suspended in 10x (v/w) Buffer W (100 mM Tris pH 8.0, 150 mM NaCl), sonicated and clarified by centrifugation. Cell lysate was loaded onto a Strep-Trap HP column (5 mL, GE Healthcare), equilibrated according to manufacturer’s instructions, and unbound protein was washed out with Buffer W (5 column volumes). *ID*-EncFtn was eluted by Buffer E (100 mM Tris pH 8.0, 150 mM NaCl, 2.5 mM desthiobiotin).

### Protein quantification

Both untagged and His-tagged EncFtn proteins were quantified using the Pierce BCA Assay kit (Thermo Fisher Scientific) following the manufacturer’s instructions for the Test-tube procedure. Concentration of Enc:EncFtn proteins was determined using the Beer-Lambert equation, measuring absorbance at 280 nm with a Nanodrop Spectrophotometer (Thermo Scientific) and using an extinction coefficient calculated by ProtParam tool on the Expasy platform entering a sequence composed of 3 x Enc and 2 x Enc-Ftn sequences. This ratio was determined by examination of SDS-PAGE gels of Enc:EncFtn proteins and densitometry of bands.

### Transmission electron microscopy

TEM imaging was performed as previously described (He et al., 2016) on purified Enc:EncFtn proteins. Proteins were diluted in Buffer GF to a final concentration of 0.1 mg/ml before being spotted on glow-discharged 300 mesh carbon-coated copper grids. Excess liquid was removed by blotting with filter paper (Whatman, UK) before the sample was washed three times with distilled water prior to being stained with 0.2 % uranyl acetate, blotted again, and air-dried. Grids were imaged using a Hitachi HT7800 transmission electron microscope and images were collected with an EMSIS Xarosa camera.

### Protein crystallization and X-ray data collection

EncFtn-sH variants (E31A or E34A) were concentrated to 10 mg/ml (based on extinction coefficient calculation) and subjected to crystallization under similar conditions to the wild-type protein (WT) (Table 10). Crystallization drops were set up in 24-well Linbro plates using the hanging drop vapor diffusion method at 292 K. Glass coverslips were set up with 1–2 μl protein mixed with 1 μl well solution and sealed over 1 ml of well solution. Crystals appeared after one week to two months and were mounted using a LithoLoop (Molecular Dimensions Limited, UK), transferred briefly to a cryoprotection solution containing well solution supplemented with 1 mM FeSO_4_ (in 0.1% (v/v) HCl) and 20% (v/v) PEG 200, and were subsequently flash cooled in liquid nitrogen.

**Table 10.** Crystallization conditions for EncFtn-sH variants.

| Variant | X-ray diffraction resolution | Space group | Crystallization conditions | Cryo-protectant |
| --- | --- | --- | --- | --- |
| EncFtn-E31A | 2.75 Å | *P* 1 2<sub>1</sub> 1 | 15% (w/v) PEG 3350 + 0.15 M Calcium Acetate | 15% (w/v) PEG 3350 + 0.15 M Calcium Acetate + 20% (v/v) PEG 200 |
| EncFtn-E34A | 2.19 Å | *P* 1 2<sub>1</sub> 1 | 14% (w/v) PEG 3350 + 0.15 M Calcium Acetate | 15% (w/v) PEG 3350 + 0.14 M Calcium Acetate + 20% (v/v) PEG 200 + 50 mM Tris-HCl (pH 8.5) + 1 mM acidic FeSO<sub>4</sub> |

The crystallization solutions were diluted from stock solutions prepared as follows: 50% (w/v) PEG 3350, 100% (v/v) PEG 200, 1 M calcium acetate, 1 M magnesium formate, 1 M Tris-HCl (pH 8.5). 100 mM FeSO<sub>4</sub> was freshly dissolved in 0.1% (v/v) HCl before diluting in the cryo-protectant solutions.

All crystallographic datasets were collected at 10–100 eV above the experimentally determined Fe-*K*α edge on the macromolecular crystallography beamlines at Diamond Light Source (Didcot, UK) at 100 K using Pilatus 6M detectors. Diffraction data were integrated and scaled using XDS (Kabsch, 2010) and symmetry related reflections were merged with Aimless (Evans, 2011). Data collection statistics are shown in Table 11. The resolution cut-off used for structure determination and refinement was determined based on the CC_1/2_ criterion proposed by (Karplus and Diederichs, 2012).

**Table 11.**
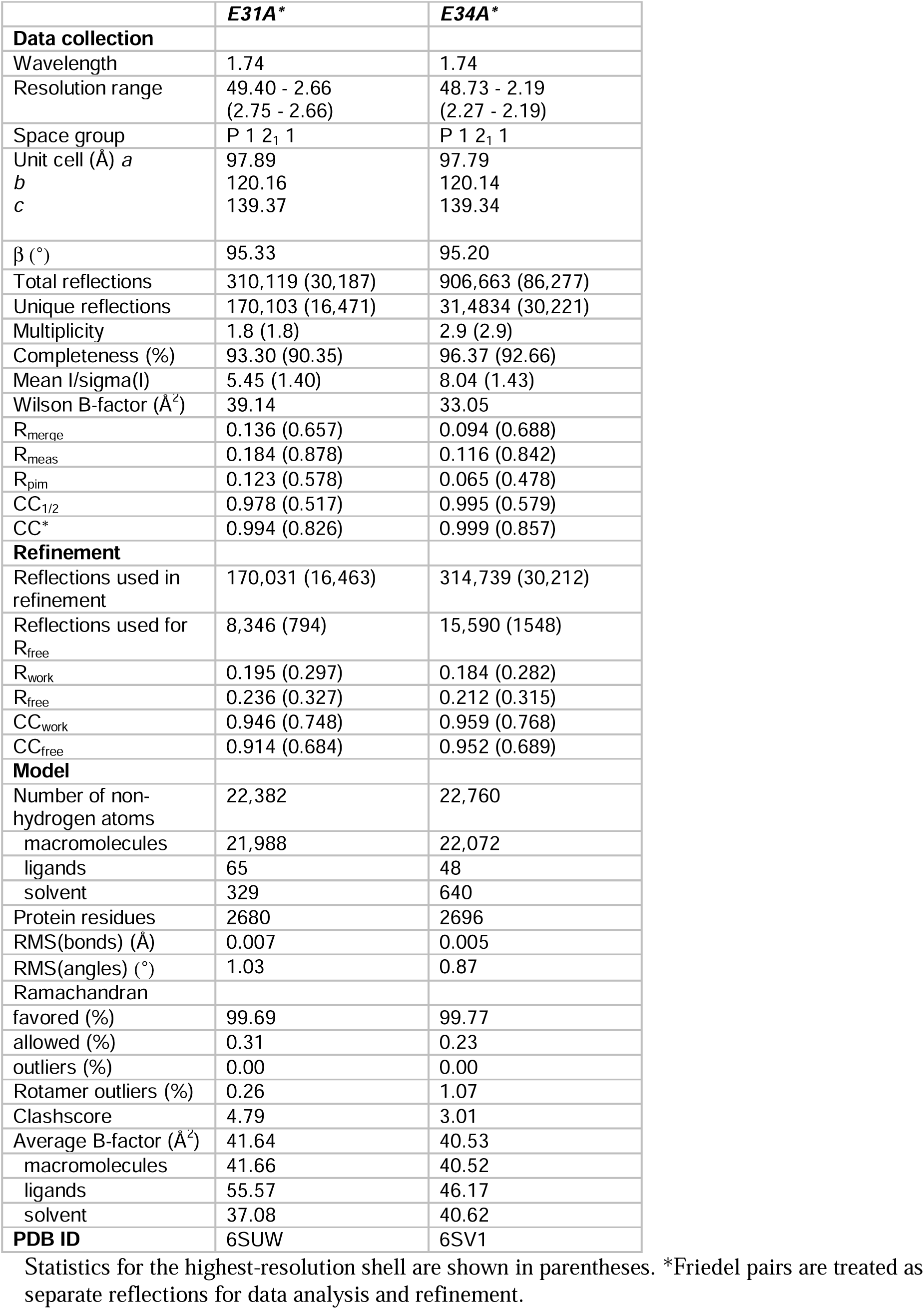
Data collection and refinement statistics.

|  | **E31A\*** | **E34A\*** |
| --- | --- | --- |
| **Data collection** |  |  |
| Wavelength | 1.74 | 1.74 |
| Resolution range | 49.40 - 2.66<br>(2.75 - 2.66) | 48.73 - 2.19<br>(2.27 - 2.19) |
| Space group | P 1 2<sub>1</sub> 1 | P 1 2<sub>1</sub> 1 |
| Unit cell (Å) *a* | 97.89 | 97.79 |
| *b* | 120.16 | 120.14 |
| *c* | 139.37 | 139.34 |
| β (°) | 95.33 | 95.20 |
| Total reflections | 310,119 (30,187) | 906,663 (86,277) |
| Unique reflections | 170,103 (16,471) | 31,4834 (30,221) |
| Multiplicity | 1.8 (1.8) | 2.9 (2.9) |
| Completeness (%) | 93.30 (90.35) | 96.37 (92.66) |
| Mean I/sigma(I) | 5.45 (1.40) | 8.04 (1.43) |
| Wilson B-factor (Å<sup>2</sup>) | 39.14 | 33.05 |
| R<sub>merge</sub> | 0.136 (0.657) | 0.094 (0.688) |
| R<sub>meas</sub> | 0.184 (0.878) | 0.116 (0.842) |
| R<sub>pim</sub> | 0.123 (0.578) | 0.065 (0.478) |
| CC<sub>1/2</sub> | 0.978 (0.517) | 0.995 (0.579) |
| CC* | 0.994 (0.826) | 0.999 (0.857) |
| **Refinement** |  |  |
| Reflections used in refinement | 170,031 (16,463) | 314,739 (30,212) |
| Reflections used for R<sub>free</sub> | 8,346 (794) | 15,590 (1548) |
| R<sub>work</sub> | 0.195 (0.297) | 0.184 (0.282) |
| R<sub>free</sub> | 0.236 (0.327) | 0.212 (0.315) |
| CC<sub>work</sub> | 0.946 (0.748) | 0.959 (0.768) |
| CC<sub>free</sub> | 0.914 (0.684) | 0.952 (0.689) |
| **Model** |  |  |
| Number of non-hydrogen atoms | 22,382 | 22,760 |
| macromolecules | 21,988 | 22,072 |
| ligands | 65 | 48 |
| solvent | 329 | 640 |
| Protein residues | 2680 | 2696 |
| RMS(bonds) (Å) | 0.007 | 0.005 |
| RMS(angles) (°) | 1.03 | 0.87 |
| Ramachandran |  |  |
| favored (%) | 99.69 | 99.77 |
| allowed (%) | 0.31 | 0.23 |
| outliers (%) | 0.00 | 0.00 |
| Rotamer outliers (%) | 0.26 | 1.07 |
| Clashscore | 4.79 | 3.01 |
| Average B-factor (Å<sup>2</sup>) | 41.64 | 40.53 |
| macromolecules | 41.66 | 40.52 |
| ligands | 55.57 | 46.17 |
| solvent | 37.08 | 40.62 |
| **PDB ID** | 6SUW | 6SV1 |

Statistics for the highest-resolution shell are shown in parentheses. \*Friedel pairs are treated as separate reflections for data analysis and refinement.

The structures of the EncFtn-sH variants were determined by molecular replacement using decameric wild-type protein (PDB ID: 5DA5) as the search model (He et al., 2016). A single solution comprising three decamers in the asymmetric unit was found by molecular replacement using Phaser (McCoy et al., 2007). The initial model was rebuilt using Phenix.autobuild (Adams et al., 2010) followed by cycles of refinement with Phenix.refine (Afonine et al., 2012), with manual rebuilding and model inspection in Coot (Emsley et al., 2010). The final model was refined with isotropic B-factors, torsional NCS restraints, and with anomalous group refinement to correctly model the iron ions in the ferroxidase center. The model was validated using MolProbity (Chen et al., 2010). Structural superimpositions were calculated using Coot (Emsley et al., 2010). Crystallographic figures were generated with PyMOL. Model refinement statistics are shown in Table 11. The final models and experimental data are deposited in the PDB.

### Mass spectrometry analysis

Sample preparation, instrumentation, native MS conditions, ion mobility-MS conditions and the method of collision cross section (CCS) determination have all been reported in previous works by our group (He et al., 2016; Ross et al., 2020). For collision induced unfolding (CIU) experiments, protein samples were prepared and the Synapt G2 was set up as for IM-MS. Samples were analyzed with incremental increasing trap voltages between 10 V to 60 V, using 2 V increments. Data was processed using Spyder CIUSuite (Dixit et al., 2018).

### Ferroxidase assay of EncFtn proteins

Enzymatic activity of EncFtn proteins was tested by ferroxidase assay, as previously described (He et al., 2019). Fe(II) aliquots were prepared under anaerobic conditions by dissolving FeSO_4_·7H_2_O in 0.1% (v/v) HCl. Purified protein was diluted anaerobically in Buffer H (10 mM HEPES, pH 8.0, 150 mM NaCl), previously purged with gaseous nitrogen, to a final concentration of 10 µM (EncFtn).

Protein and ferrous iron samples were added to a quartz cuvette (Hellma) under aerobic conditions at a final concentration of 10 µM and 50 µM, respectively, corresponding to a FOC:Fe(II) ratio of 1:10.

Absorbance at 315 nm was recorded every second for 1500 s by a UV-visible spectrophotometer (PerkinElmer Lambda 35), using the TimeDrive software. The same experiment was performed in the absence of the enzyme to determine the oxidation of ferrous salts by atmospheric oxygen. The same setup was adopted for recording the activity of EncFtn-E34A and EncFtn-E31A/E34A in a separate assay at higher concentration (20 µM EncFtn and 100 µM Fe(II)) for comparison purposes. Data presented here are the mean of three technical replicates of time zero-subtracted progress curves with standard deviations calculated from the mean.

Calculation of the EncFtn enzymatic reaction initial rate (*v*_0_) was made by applying the Linear Regression tool in GraphPad (Prism8) on the absorbance at 315 nm measured for the first 200 s, when curves are still linear and following an order zero kinetics (Figure 6-supplement 2, Table 4). Slopes obtained from these curves correspond to initial reaction rate. *v*_0_^variant^/ *v*_0_^wild-type^ factors were calculated by dividing the initial rates of the variant enzymes by the initial rate of the wild-type protein (Table 6).

### Ferroxidase assay of Enc:EncFtn protein complexes

Ferroxidase assay of Enc:EncFtn protein complexes was performed in the same conditions as those used for EncFtn, but with different protein and iron salt concentration. Enc:EncFtn was diluted to a final concentration of 25 µM which corresponds to 10 µM EncFtn based on a 3:2 Enc:EncFtn ratio, as previously observed by SDS-PAGE by our group (Figure 1A in (He et al., 2016). Fe(II) sample was diluted to 50 µM in the reaction system to maintain a final ratio of 1:10 FOC:Fe(II).

Calculation of the Enc-EncFtn enzymatic reaction initial rate (*v*_0_) and *v*_0_^variant^/ *v*_0_^wild-type^ factors was carried out as described in the above section (Figure 6-supplement 2, Tables 5 and 6).

### Zinc inhibition of ferroxidase activity of EncFtn proteins

Ferroxidase assay was performed as above with EncFtn (10 µM) and Fe(II) samples (50 µM) in the presence of 34 µM ZnSO_4_·7H_2_O. The chosen concentration corresponds to the IC_50_ value previously determined for EncFtn-StrepII, under the same experimental conditions by our group (He et al., 2019). Oxygen-free metal samples were added to the quartz cuvette under aerobic conditions, followed by the protein sample. Data were replicated three times and means and standard deviation of time zero-subtracted progress curves were calculated. A negative control was performed by monitoring A_315_ of Zn(II) and Fe(II) salts mixed in the absence of enzyme.

### Monitoring quenching of protein intrinsic fluorescence upon metal-binding

Experiments were carried out aerobically in a quartz cuvette (Hellma), using a Cary Eclipse Fluorescence spectrophotometer. Protein was diluted in Buffer GF to a final concentration of 20 µM, corresponding to 10 µM for the entry site. ZnSO_4_•7H_2_O stock was prepared in deionized water. The instrument excitation wavelength was set to 280 nm, corresponding to maximum absorption for tryptophan residues.

Two distinct modes of detection were used; Kinetic option on the Analysis software package was chosen to record protein fluorescence emission signal at specific wavelengths over time. Pre-scans were carried out to find the optimal emission wavelength (ranging from 322 nm to 355 nm) for each protein variant. Metal aliquots (0.15 or 0.3 molar equivalents) were added to the cuvette while pausing the data collection for fewer than five seconds. The final data represent average of 10 data points at equilibrium upon each metal addition.

The Scan option of the Analysis software package was chosen to record emission in the 290-400 nm range, allowing detection of possible shifts in emission peak maximum upon metal addition. Spectra were recorded in triplicate at equilibrium.

### Native FT-ICR mass spectrometry

High resolution native mass spectrometry was performed on a 12T SolariX 2XR FT-ICR MS (Bruker Daltonics) equipped with a nanoelectrospray source. Protein samples were buffer exchanged into ammonium acetate (100 mM; pH 8.0) prior to direct infusion. Source conditions and ion optics were optimized to transmit native proteins ions and when required, Continual Accumulation of Selected Ions (CASI) was employed to isolate charge states of interest. Typically, 2 Megaword data was collected in QPD (2ω) mode to produce a 6 second FID, which resulted in a typical mass resolving power of ca. 300,000. The resulting data was processed using Data Analysis (Bruker Daltonics) and theoretical isotope patterns were calculated using IsotopePattern (Bruker Daltonics). For metal titrations, fresh iron(II) chloride tetrahydrate or zinc acetate in 0.1% (v/v) HCl was added to *ID*-EncFtn (1:1 metal:protein concentration) prior to buffer exchange into ammonium acetate (100 mM; pH 8.0).

### Accession codes and datasets

Data sets supporting this paper have been deposited in appropriate public data repositories. Please see figure legends and tables for links to these.

---

**Table S9.**
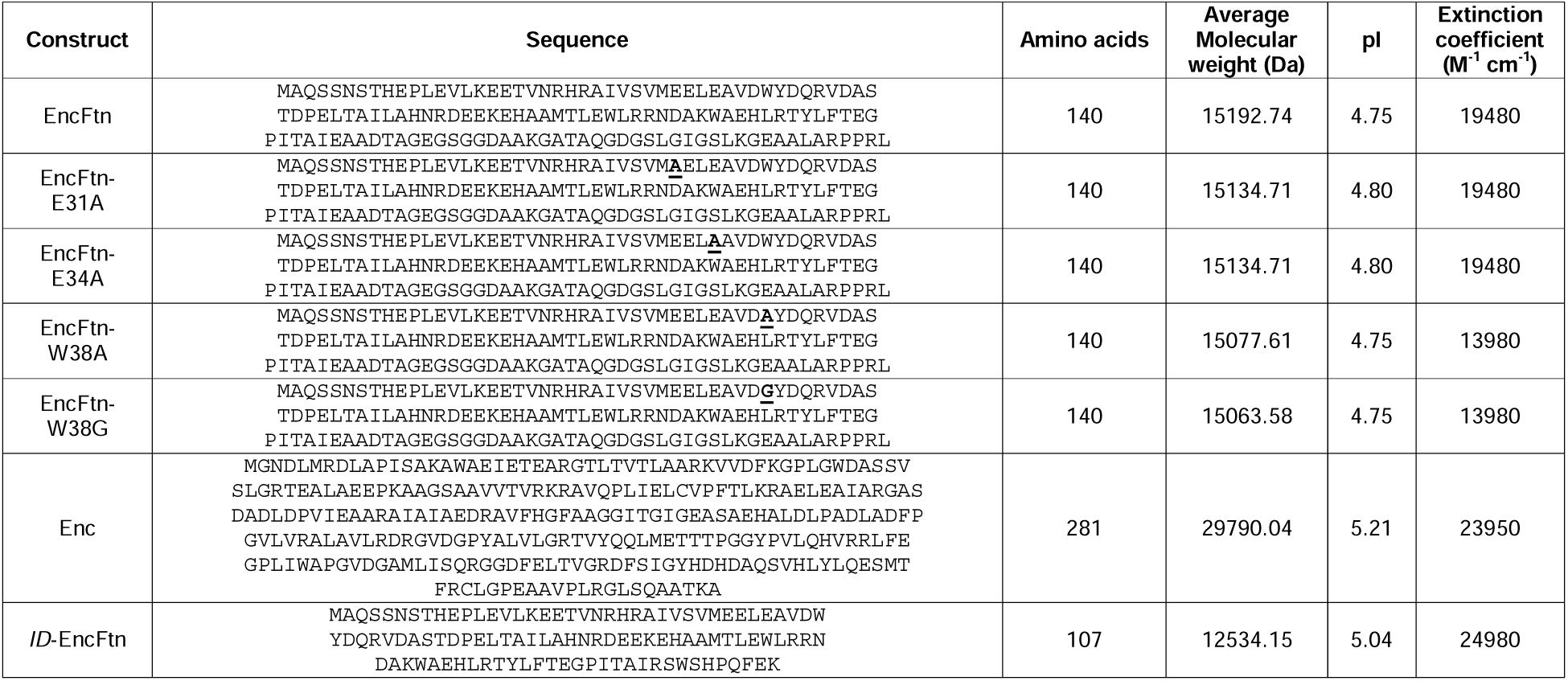
Constructs used in this study.

## Author contributions

**CP**, Conception and design, Acquisition of data, Analysis and interpretation of data, Drafting or revising the article.

**JR**, Conception and design, Acquisition of data, Analysis and interpretation of data. Drafting or revising the article.

**DH**, Conception and design, Acquisition of data, Analysis and interpretation of data.

**KJG**, ID-MS methods development and acquisition of data, Analysis and interpretation of data

**WAS**, Acquisition of data.

**LA**, Acquisition of data.

**CLM**, Acquisition of data.

**KJW**, Acquisition of data, Interpretation of data.

**DJC**, Conception and design, Acquisition of data, Analysis and interpretation of data, Drafting or revising the article.

**JMW**, Conception and design, Acquisition of data, Analysis and interpretation of data, Drafting or revising the article.

## Abbreviations

Enc: encapsulin;
EncFtn: encapsulated ferritin;
FOC: ferroxidase center;
MS: mass spectrometry;
nESI: Native nanoelectrospray ionization;
CIU: collision induced unfolding;
FT-ICR: Fourier-transform ion cyclotron resonance;
ID: isotope depletion.

## Acknowledgements

We would like to thank staff at the Newcastle University Electron Microscopy Research Services for assistance with TEM. We would like to thank Diamond Light Source for beamtime (proposal mx9487), and the staff of beamlines I02, I03 and I24 for assistance with crystal testing and data collection. We would like to thank Dr Arnaud Baslé, Prof. Dominic Campopiano and Gregor Skeldon for their constructive advice throughout this project.

## Figure Supplements

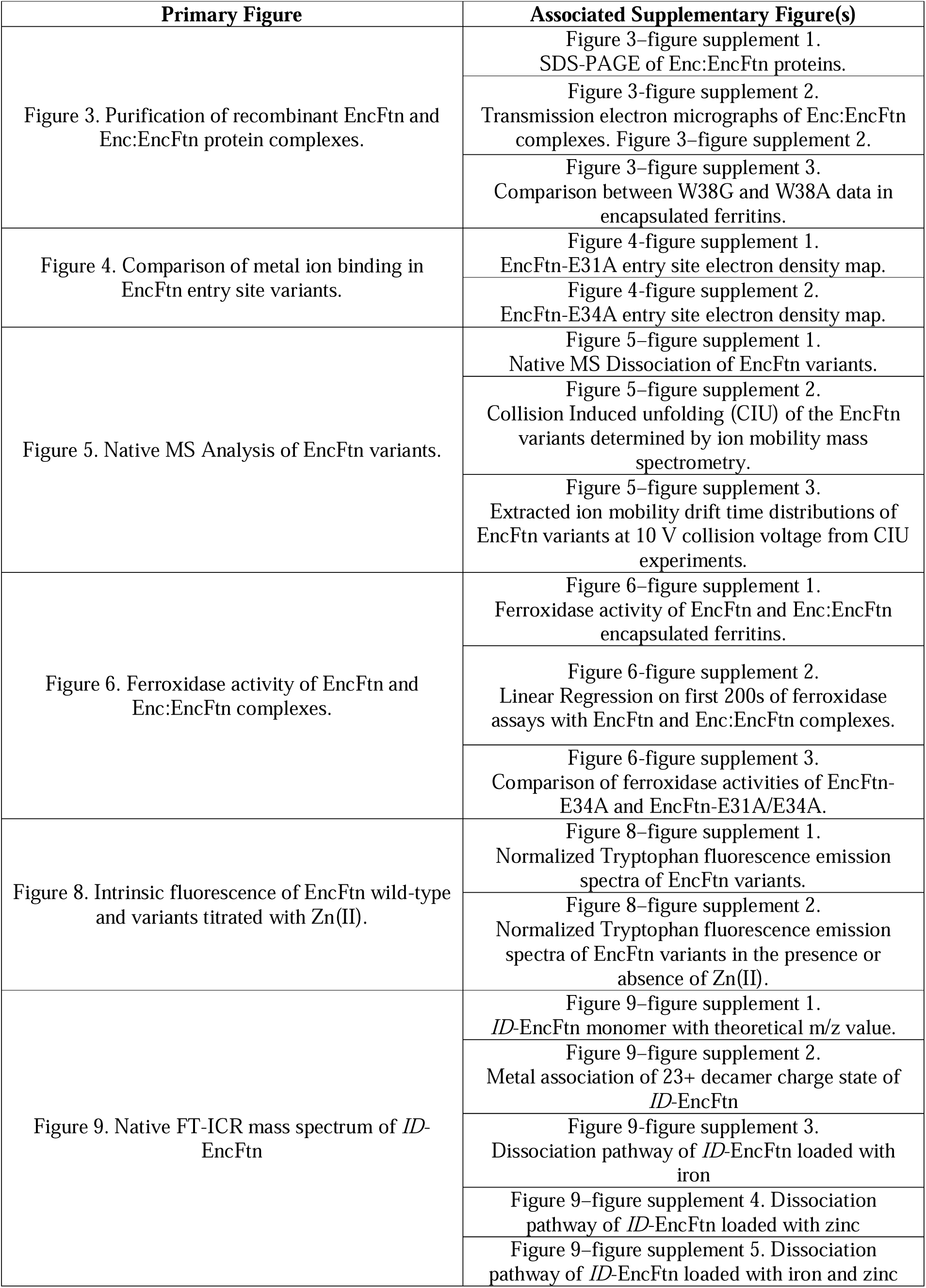

## References

Adams PD, Afonine P V., Bunkóczi G, Chen VB, Davis IW, Echols N, Headd JJ, Hung L-W, Kapral GJ, Grosse-Kunstleve RW, McCoy AJ, Moriarty NW, Oeffner R, Read RJ, Richardson DC, Richardson JS, Terwilliger TC, Zwart PH. 2010. PHENIX□: a comprehensive Python-based system for macromolecular structure solution. Acta Crystallogr Sect D Biol Crystallogr 66:213–221. doi:10.1107/S0907444909052925

Afonine P V., Grosse-Kunstleve RW, Echols N, Headd JJ, Moriarty NW, Mustyakimov M, Terwilliger TC, Urzhumtsev A, Zwart PH, Adams PD. 2012. Towards automated crystallographic structure refinement with phenix.refine. Acta Crystallogr Sect D Biol Crystallogr 68:352–367. doi:10.1107/S0907444912001308

Almiron M, Link AJ, Furlong D, Kolter R. 1992. A novel DNA-binding protein with regulatory and protective roles in starved Escherichia coli. Genes Dev 6:2646–2654. doi:10.1101/gad.6.12b.2646

Andrews S. 1998. Iron storage in bacteria. Adv Microb Physiol 40:281–351.

Andrews SC. 2010. The Ferritin-like superfamily: Evolution of the biological iron storeman from a rubrerythrin-like ancestor. Biochim Biophys Acta - Gen Subj 1800:691–705. doi:10.1016/j.bbagen.2010.05.010

Baaghil S, Lewin A, Moore GR, Le Brun NE. 2003. Core Formation in Escherichia coli Bacterioferritin Requires a Functional Ferroxidase Center †. Biochemistry 42:14047–14056. doi:10.1021/bi035253u

Behera RK, Torres R, Tosha T, Bradley JM, Goulding CW, Theil EC. 2015. Fe2+ substrate transport through ferritin protein cage ion channels influences enzyme activity and biomineralization. JBIC J Biol Inorg Chem 20:957–969. doi:10.1007/s00775-015-1279-x

Bellapadrona G, Stefanini S, Zamparelli C, Theil EC, Chiancone E. 2009. Iron Translocation into and out of Listeria innocua Dps and Size Distribution of the Protein-enclosed Nanomineral Are Modulated by the Electrostatic Gradient at the 3-fold “Ferritin-like” Pores. J Biol Chem 284:19101–19109. doi:10.1074/jbc.M109.014670

Cardenas JP, Quatrini R, Holmes DS. 2016. Aerobic Lineage of the Oxidative Stress Response Protein Rubrerythrin Emerged in an Ancient Microaerobic, (Hyper)Thermophilic Environment. Front Microbiol 7. doi:10.3389/fmicb.2016.01822

Chasteen ND, Harrison PM. 1999. Mineralization in Ferritin: An Efficient Means of Iron Storage. J Struct Biol 126:182–194. doi:10.1006/jsbi.1999.4118

Chen V, Arendall W, Headd J, Keedy D, Immormino R, Kapral G, Murray L, Richardson J, Richardson D. 2010. MolProbity: all-atom structure validation for macromolecular crystallography. Acta Crystallogr Sect D Biol Crystallogr 66:12–21. doi:10.1107/S0907444909042073

Chiancone E, Ceci P. 2010. The multifaceted capacity of Dps proteins to combat bacterial stress conditions: Detoxification of iron and hydrogen peroxide and DNA binding. Biochim Biophys Acta - Gen Subj 1800:798–805. doi:10.1016/j.bbagen.2010.01.013

Choi SH, Baumler DJ, Kaspar CW. 2000. Contribution of dps to Acid Stress Tolerance and Oxidative Stress Tolerance in Escherichia coli O157:H7. Appl Environ Microbiol 66:3911–3916. doi:10.1128/AEM.66.9.3911-3916.2000

De Martino M, Ershov D, van den Berg PJ, Tans SJ, Meyer AS. 2016. Single-Cell Analysis of the Dps Response to Oxidative Stress. J Bacteriol 198:1662–1674. doi:10.1128/JB.00239-16

Dillard BD, Demick JM, Adams MWW, Lanzilotta WN. 2011. A cryo-crystallographic time course for peroxide reduction by rubrerythrin from Pyrococcus furiosus. JBIC J Biol Inorg Chem 16:949–959. doi:10.1007/s00775-011-0795-6

Dixit SM, Polasky DA, Ruotolo BT. 2018. Collision induced unfolding of isolated proteins in the gas phase: past, present, and future. Curr Opin Chem Biol 42:93–100. doi:10.1016/j.cbpa.2017.11.010

Ebrahimi KH, Bill E, Hagedoorn P-L, Hagen WR. 2016. Spectroscopic evidence for the role of a site of the di-iron catalytic center of ferritins in tuning the kinetics of Fe(<scp>ii</scp>) oxidation. Mol Biosyst 12:3576–3588. doi:10.1039/C6MB00235H

Ebrahimi KH, Hagedoorn P-L, Jongejan J a, Hagen WR. 2009. Catalysis of iron core formation in Pyrococcus furiosus ferritin. J Biol Inorg Chem 14:1265–74. doi:10.1007/s00775-009-0571-z

Emsley P, Lohkamp B, Scott WG, Cowtan K. 2010. Features and development of Coot. Acta Crystallogr Sect D Biol Crystallogr 66:486–501. doi:10.1107/S0907444910007493

Eschweiler JD, Rabuck-Gibbons JN, Tian Y, Ruotolo BT. 2015. CIUSuite: A Quantitative Analysis Package for Collision Induced Unfolding Measurements of Gas-Phase Protein Ions. Anal Chem 87:11516–11522. doi:10.1021/acs.analchem.5b03292

Evans PR. 2011. An introduction to data reduction: space-group determination, scaling and intensity statistics. Acta Crystallogr Sect D Biol Crystallogr 67:282–292. doi:10.1107/S090744491003982X

Foster AW, Osman D, Robinson NJ. 2014. Metal Preferences and Metallation. J Biol Chem 289:28095–28103. doi:10.1074/jbc.R114.588145

Frausto da Silva J, Williams R. 1991. The biological chemistry of the elements: the inorganic chemistry of life. Oxford Univ Press 20:62–63. doi:10.1016/0307-4412(92)90039-O

Frolow F, Kalb A, Yariv J. 1994. Structure of a unique twofold symmetric haem-binding site. Nat Struct Biol 453–460.

Gallagher KJ, Palasser M, Hughes S, Mackay CL, Kilgour DPA, Clarke DJ. 2020. Isotope Depletion Mass Spectrometry (ID-MS) for Accurate Mass Determination and Improved Top-Down Sequence Coverage of Intact Proteins. J Am Soc Mass Spectrom 31:700–710. doi:10.1021/jasms.9b00119

Giessen TW, Orlando BJ, Verdegaal AA, Chambers MG, Gardener J, Bell DC, Birrane G, Liao M, Silver PA. 2019. Large protein organelles form a new iron sequestration system with high storage capacity. Elife 8. doi:10.7554/eLife.46070

Grant R, Filman D, Finkel S, Kolter R, Hogle J. 1998. The crystal structure of Dps, a ferritin homolog that binds and protects DNA. Nat Struct Biol 5:294–303.

Hagen W, Hagedoorn P-L, Ebrahimi K. 2017. The working of ferritin: a crossroad of options. Metallomics 9:595–605.

Hagen WR, Hagedoorn PL, Honarmand Ebrahimi K. 2017. The workings of ferritin: A crossroad of opinions. Metallomics. doi:10.1039/c7mt00124j

Hall Z, Hernández H, Marsh JA, Teichmann SA, Robinson C V. 2013. The Role of Salt Bridges, Charge Density, and Subunit Flexibility in Determining Disassembly Routes of Protein Complexes. Structure 21:1325–1337. doi:10.1016/j.str.2013.06.004

Harrison PM, Arosio P. 1996. The ferritins: molecular properties, iron storage function and cellular regulation. Biochim Biophys Acta - Bioenerg 1275:161–203. doi:10.1016/0005-2728(96)00022-9

He D, Hughes S, Vanden-Hehir S, Georgiev A, Altenbach K, Tarrant E, Mackay CL, Waldron KJ, Clarke DJ, Marles-Wright J. 2016. Structural characterization of encapsulated ferritin provides insight into iron storage in bacterial nanocompartments. Elife 5. doi:10.7554/eLife.18972

He D, Piergentili C, Ross J, Tarrant E, Tuck LR, Mackay CL, McIver Z, Waldron KJ, Clarke DJ, Marles-Wright J. 2019. Conservation of the structural and functional architecture of encapsulated ferritins in bacteria and archaea. Biochem J 476:975–989. doi:10.1042/bcj20180922

Hempstead PD, Hudson AJ, Artymiuk PJ, Andrews SC, Banfield MJ, Guest JR, Harrison PM. 1994. Direct observation of the iron binding sites in a ferritin. FEBS Lett 350:258–62. doi:10.1016/0014-5793(94)00781-0

Kabsch W. 2010. Integration, scaling, space-group assignment and post-refinement. Acta Crystallogr Sect D Biol Crystallogr 66:133–144. doi:10.1107/S0907444909047374

Karplus PA, Diederichs K. 2012. Linking Crystallographic Model and Data Quality. Science (80-) 336:1030–1033. doi:10.1126/science.1218231

Le Brun NE, Andrews SC, Guest JR, Harrison PM, Moore GR, Thomson AJ. 1995. Identification of the ferroxidase centre of Escherichia coli bacterioferritin. Biochem J 312:385–392. doi:10.1042/bj3120385

Liu XS, Patterson LD, Miller MJ, Theil EC. 2007. Peptides Selected for the Protein Nanocage Pores Change the Rate of Iron Recovery from the Ferritin Mineral. J Biol Chem 282:31821–31825. doi:10.1074/jbc.C700153200

Marklund EG, Degiacomi MT, Robinson C V., Baldwin AJ, Benesch JLP. 2015. Collision Cross Sections for Structural Proteomics. Structure 23:791–799. doi:10.1016/j.str.2015.02.010

Martinez A, Kolter R. 1997. Protection of DNA during oxidative stress by the nonspecific DNA-binding protein Dps. J Bacteriol 179:5188–5194. doi:10.1128/JB.179.16.5188-5194.1997

Masuda T, Goto F, Yoshihara T, Mikami B. 2010. Crystal structure of plant ferritin reveals a novel metal binding site that functions as a transit site for metal transfer in ferritin. J Biol Chem. doi:10.1074/jbc.M109.059790

McCoy AJ, Grosse-Kunstleve RW, Adams PD, Winn MD, Storoni LC, Read RJ. 2007. Phaser crystallographic software. J Appl Crystallogr 40:658–674. doi:10.1107/S0021889807021206

Nair S, Finkel SE. 2004. Dps Protects Cells against Multiple Stresses during Stationary Phase. J Bacteriol 186:4192–4198. doi:10.1128/JB.186.13.4192-4198.2004

Pesek J, Büchler R, Albrecht R, Boland W, Zeth K. 2011. Structure and Mechanism of Iron Translocation by a Dps Protein from Microbacterium arborescens. J Biol Chem 286:34872–34882. doi:10.1074/jbc.M111.246108

Pfaffen S, Abdulqadir R, Le Brun NE, Murphy MEP. 2013. Mechanism of Ferrous Iron Binding and Oxidation by Ferritin from a Pennate Diatom. J Biol Chem 288:14917–14925. doi:10.1074/jbc.M113.454496

Proulx-Curry PM, Chasteen ND. 1995. Molecular aspects of iron uptake and storage in ferritin. Coord Chem Rev 144:347–368. doi:10.1016/0010-8545(95)01148-I

Ratnayake DB, Wai SN, Shi Y, Amako K, Nakayama H, Nakayama K. 2000. Ferritin from the obligate anaerobe Porphyromonas gingivalis: purification, gene cloning and mutant studies The GenBank accession number for the sequence reported in this paper is AB016086. Microbiology 146:1119–1127. doi:10.1099/00221287-146-5-1119

Reyes-Caballero H, Campanello GC, Giedroc DP. 2011. Metalloregulatory proteins: Metal selectivity and allosteric switching. Biophys Chem 156:103–114. doi:10.1016/j.bpc.2011.03.010

Ross J, Lambert T, Piergentili C, He D, Waldron KJ, Mackay CL, Marles-Wright J, Clarke DJ. 2020. Mass spectrometry reveals the assembly pathway of encapsulated ferritins and highlights a dynamic ferroxidase interface. Chem Commun. doi:10.1039/C9CC08130E

Schägger H. 2006. Tricine–SDS-PAGE. Nat Protoc 1:16–22. doi:10.1038/nprot.2006.4

Stillman TJ, Hempstead PD, Artymiuk PJ, Andrews SC, Hudson AJ, Treffry A, Guest JR, Harrison PM. 2001. The high-resolution X-ray crystallographic structure of the ferritin (EcFtnA) of Escherichia coli; comparison with human H ferritin (HuHF) and the structures of the Fe(3+) and Zn(2+) derivatives. J Mol Biol 307:587–603. doi:10.1006/jmbi.2001.4475

Sztukowska M, Bugno M, Potempa J, Travis J, Kurtz Jr DM. 2002. Role of rubrerythrin in the oxidative stress response of Porphyromonas gingivalis. Mol Microbiol 44:479–488. doi:10.1046/j.1365-2958.2002.02892.x

Teale FWJ, Weber G. 1957. Ultraviolet fluorescence of the aromatic amino acids. Biochem J 65:476–482. doi:10.1042/bj0650476

Theil EC, Behera RK, Tosha T. 2013. Ferritins for chemistry and for life. Coord Chem Rev 257:579–586. doi:10.1016/j.ccr.2012.05.013

Tottey S, Waldron KJ, Firbank SJ, Reale B, Bessant C, Sato K, Cheek TR, Gray J, Banfield MJ, Dennison C, Robinson NJ. 2008. Protein-folding location can regulate manganese-binding versus copper- or zinc-binding. Nature 455:1138–1142. doi:10.1038/nature07340

Treffry A, Zhao Z, Quail MA, Guest JR, Harrison PM. 1998. How the presence of three iron binding sites affects the iron storage function of the ferritin (EcFtnA) of Escherichia coli. FEBS Lett 432:213–218. doi:10.1016/S0014-5793(98)00867-9

Trefry A, Harrison PM. 1978. Incorporation and release of inorganic phosphate in horse spleen ferritin. Biochem J 171:313–320. doi:10.1042/bj1710313

Xu B, Chasteen N. 1991. Iron oxidation chemistry in ferritin. Increasing Fe/O2 stoichiometry during core formation. J Biol Chem 266:19965–70.

Yang X, Chen-Barrett Y, Arosio P, Chasteen ND. 1998. Reaction Paths of Iron Oxidation and Hydrolysis in Horse Spleen and Recombinant Human Ferritins †. Biochemistry 37:9743–9750. doi:10.1021/bi973128a

Yao H, Wang Y, Lovell S, Kumar R, Ruvinsky AM, Battaile KP, Vakser IA, Rivera M. 2012. The Structure of the BfrB–Bfd Complex Reveals Protein–Protein Interactions Enabling Iron Release from Bacterioferritin. J Am Chem Soc 134:13470–13481. doi:10.1021/ja305180n

