## Supplemental figures and tables for "Dissecting the structural and functional roles of a vicinal iron-binding site in encapsulated ferritins"

**Supplementary Figures**

**Figure 3–figure supplement 1.** SDS-PAGE of Enc:EncFtn proteins.

**Figure 3-figure supplement 2.** Transmission electron micrographs of Enc:EncFtn complexes

**Figure 3–figure supplement** **3.** Comparison between W38G and W38A data in encapsulated ferritins.

**Figure 4-figure supplement 1.** EncFtn-E31A entry site electron density map

**Figure 4-figure supplement 2.** EncFtn-E34A entry site electron density map

**Figure 5–figure supplement** **1.** Native MS Dissociation of EncFtn variants.

**Figure 5–figure supplement** **2.** Collision Induced unfolding (CIU) of the EncFtn variants determined by ion mobility mass spectrometry

**Figure 5–figure supplement** **3.** Extracted ion mobility drift time distributions of EncFtn variants at 10 V collision voltage from CIU experiments.

**Figure 6–figure supplement** **1.** Ferroxidase activity of EncFtn and Enc:EncFtn encapsulated ferritins.

**Figure 6–figure supplement** **2.** Linear Regression on first 200s of ferroxidase assays with EncFtn and Enc:EncFtn complexes.

**Figure 6–figure supplement** **3.** Comparison of ferroxidase activities of EncFtn-E34A and EncFtn-E31A/E34A.

**Figure 8–figure supplement** **1.** Normalized Tryptophan fluorescence emission spectra of EncFtn variants.

**Figure 8–figure supplement** **2.** Normalized Tryptophan fluorescence emission spectra of EncFtn variants in the presence or absence of Zn(II).

**Figure 9–figure supplement** **1.** *ID*-EncFtn monomer with theoretical *m/z* value

**Figure 9–figure supplement** **2.** Metal association of 23+ decamer charge state of *ID*-EncFtn

**Figure 9–figure supplement** **3.** Dissociation pathway of *ID*-EncFtn loaded with iron

**Figure 9–figure supplement** **4.** Dissociation pathway of *ID*-EncFtn loaded with zinc

**Figure 9–figure supplement** **5.** Dissociation pathway of *ID*-EncFtn loaded with iron and zinc

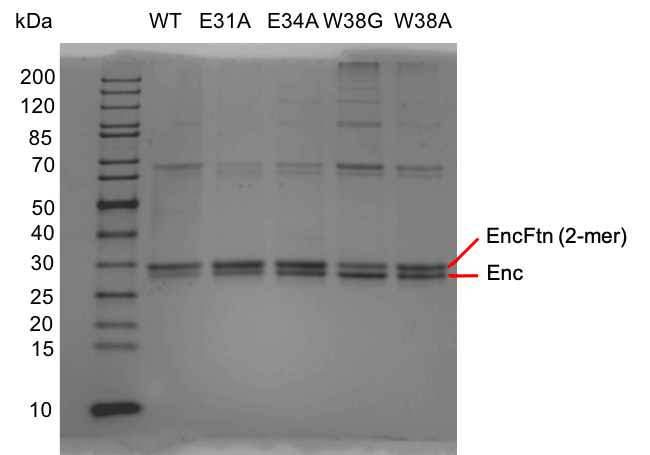

**Figure 3–figure supplement 1.**

**SDS-PAGE of Enc:EncFtn proteins.**

Samples of Enc:EncFtn protein complexes were subjected to 10 % Tris-Tricine SDS-PAGE analysis (Schägger, 2006) to assess the presence of both proteins in the complex. The EncFtn protein is primarily present as a dimeric species of 30 kDa, while the Enc is present as the 29 kDa monomer. Some higher order complex formation at multiples of two and four times the size of the major bands for the Enc and EncFtn proteins are seen in all of the variant Enc:EncFtn complexes, with varying ratios of these higher-order complexes present for the different variants.

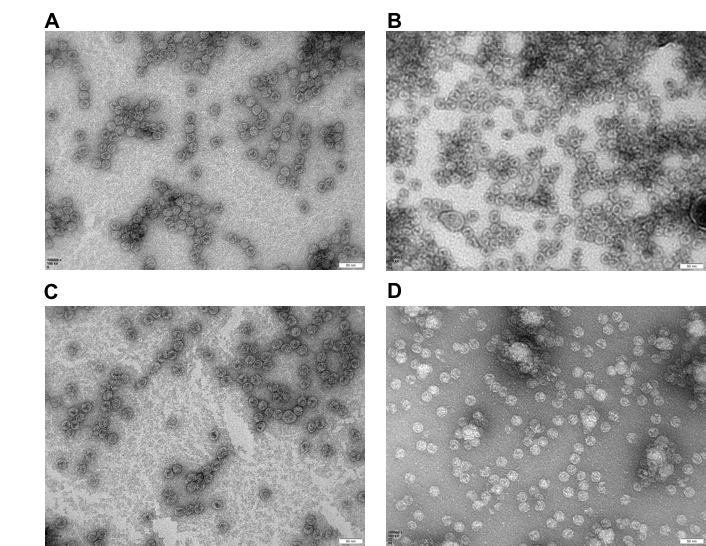

**Figure 3 – figure supplement 2**

**Transmission electron micrographs of Enc:EncFtn complexes.**

Negative stain TEM images of Enc:EncFtn nanocompartments stained with uranyl acetate. Enc:EncFtn-WT (**A**), Enc:EncFtn-E34A (**B**), Enc:EncFtn-W38A (**C**), and Enc:EncFtn-E31A (**D**). Samples were imaged at a magnification of 100,000, scale bars correspond to 50 nm. Proteins were diluted in Buffer GF to 0.1 mg/ml.

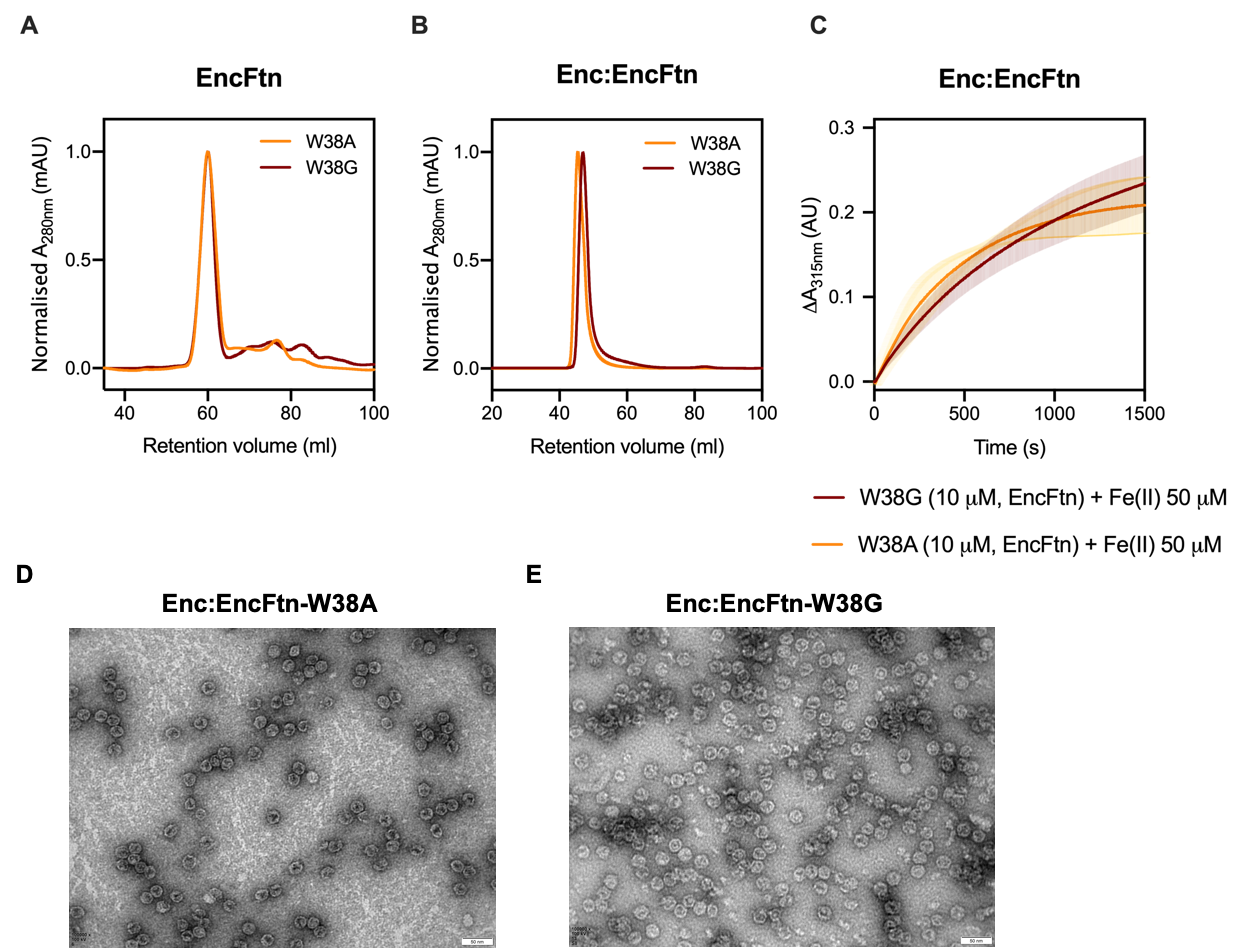

**Figure 3–figure supplement** **3.**

**Comparison between W38G and W38A data in encapsulated ferritins.**

(**A/B**) Size exclusion elution profiles of EncFtn (**A**) and Enc:EncFtn (**B**) W38G (chestnut line) and W38A (orange line) variants exhibit a similar behavior in liquid phase. doi. 10.6084/m9.figshare.9885557

(**C**) Enzymatic activities of Enc:EncFtn-W38A (orange line) and W38G (chestnut line) were tested by ferroxidase assay, as already shown for the other Enc:EncFtn proteins (Figure 6B), showing comparable activity curves. doi.10.6084/m9.figshare.9885575. (**D/E**) Negative stain TEM images of Enc:EncFtn-W38A and -W38G (100,000 magnification; 50 nm scale bar) confirm same morphology.

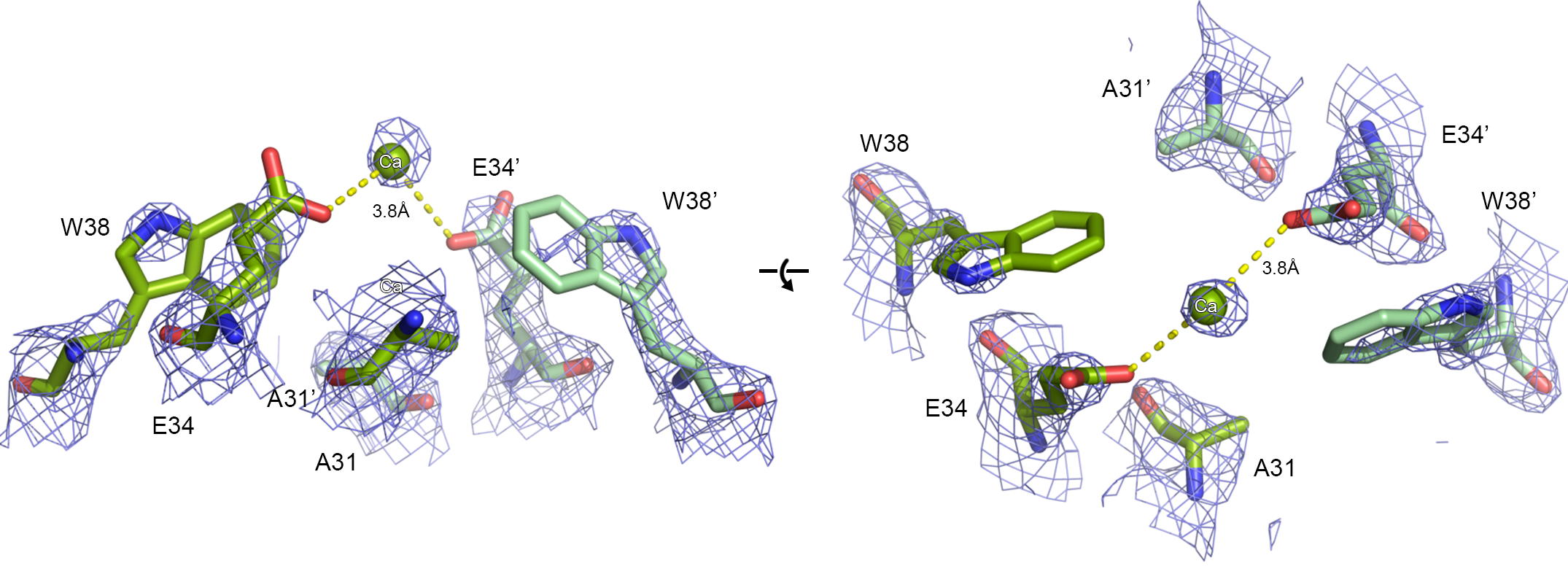

**Figure 4-figure supplement 1.** **EncFtn-E31A entry site electron density map.**

Orthogonal views of the EncFtn-E31A variant X-ray crystal structure are shown with the final experimental 2mFo-DFc electron density map contoured at 1.5 σ as a blue mesh. Entry site residues are shown as green sticks, with oxygen atoms in red and nitrogen in blue. The single coordinated calcium ion is shown as a green sphere. Residues in the entry site are labelled, with residues from the second monomer indicated with prime symbols.

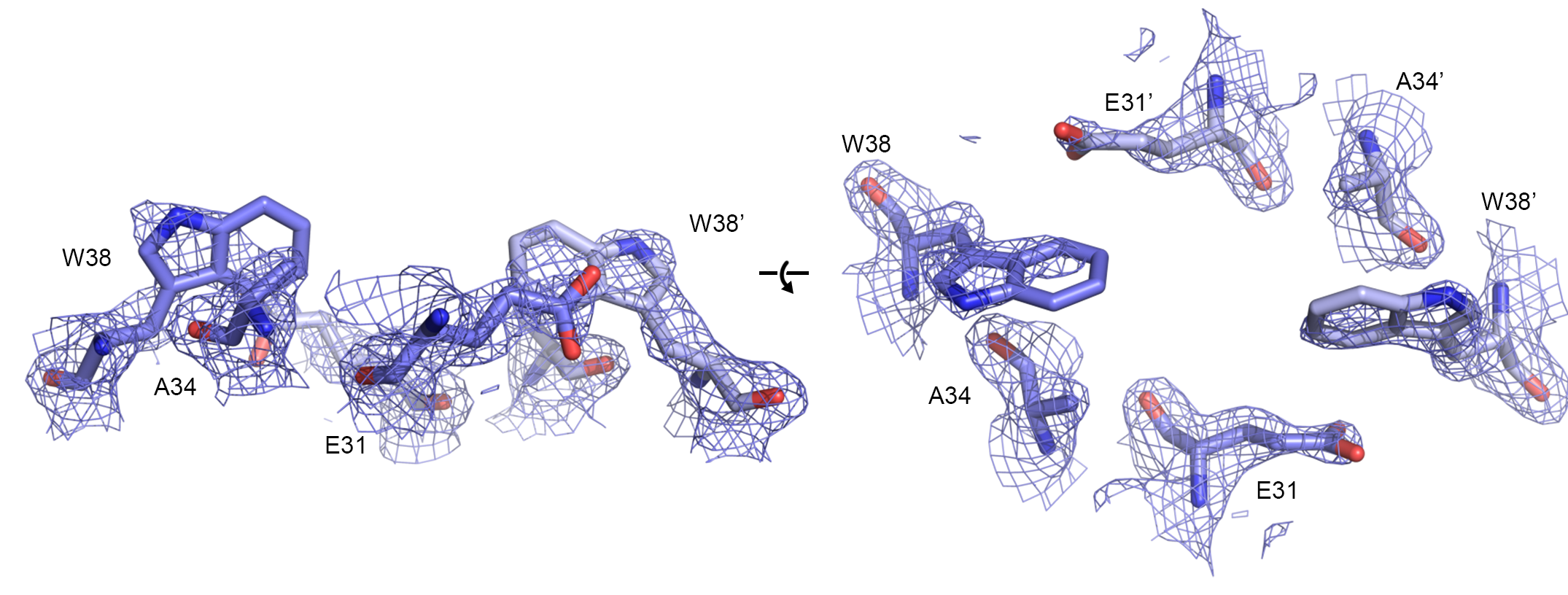

**Figure 4-figure supplement 2.** **EncFtn-E34A entry site electron density map**

Orthogonal views of the EncFtn-E34A variant X-ray crystal structure are shown with the final experimental 2mFo-DFc electron density map contoured at 1.5 σ as a blue mesh. Entry site residues are shown as blue sticks, with oxygen atoms in red and nitrogen in blue. Residues in the entry site are labelled, with residues from the second monomer indicated with prime symbols.

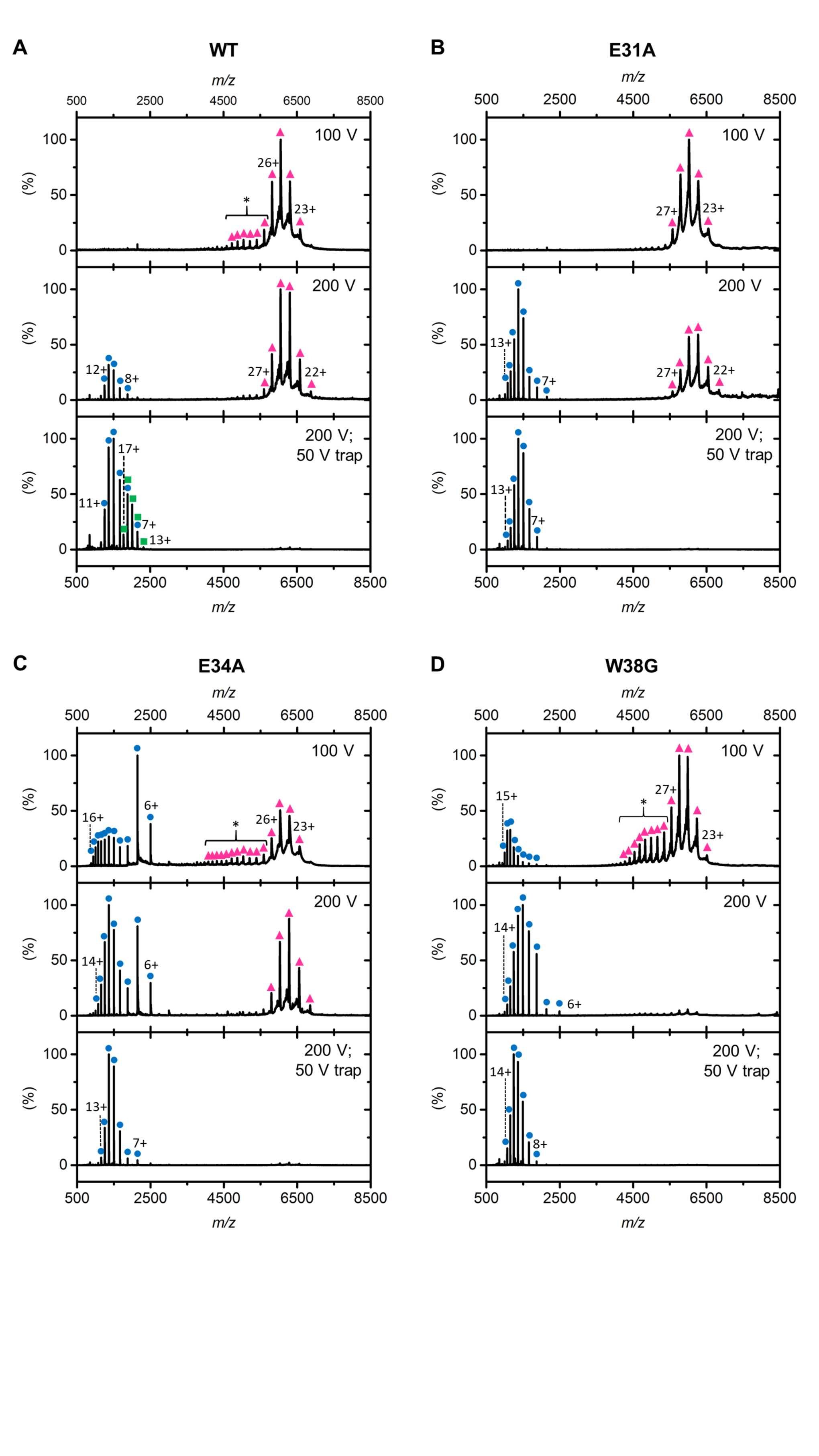

**Figure 5–figure supplement 1.**

**Native MS Dissociation of EncFtn variants.**

nESI spectra of EncFtn-WT (**A**), EncFtn-E31A (**B**), EncFtn-E34A (**C**) and EncFtn-W38G (**D**) displaying dissociation of decamer charge state distribution into a charge state distribution consistent with the respective EncFtn monomer. Oligomerization states are represented by the following colored shapes: decamer (pink triangles); monomer (blue circles); and dimer (green squares). * denotes the extended, decameric charge state observed in EncFtn-WT, EncFtn-E34A and EncFtn-W38G. The sampling cone voltage was increased from 100 V to 200 V and finally a 50 V trap voltage was applied to achieve maximum dissociation of the decamer charge state.

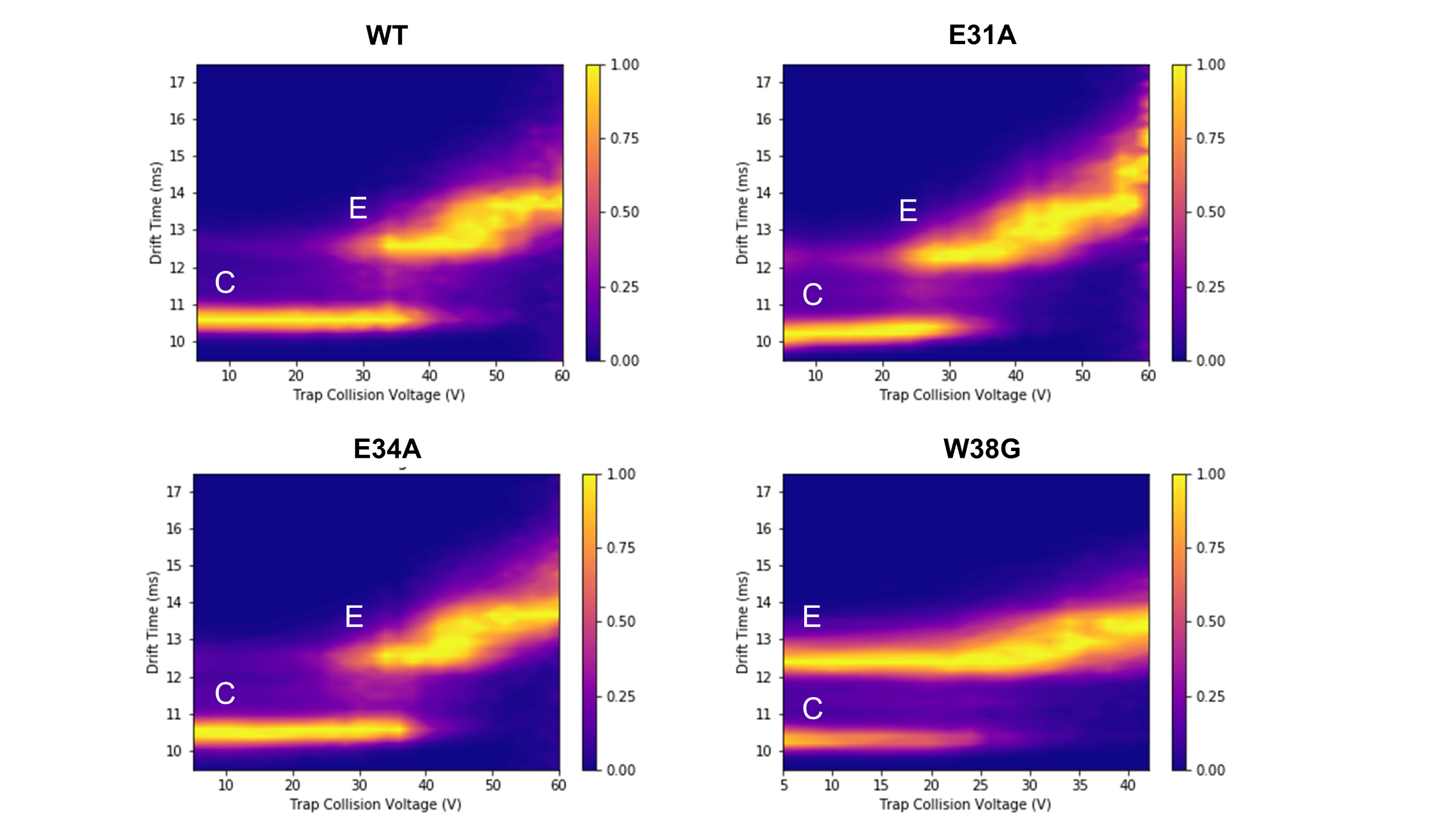

**Figure 5-figure supplement 2.**

**Collision Induced unfolding (CIU) of the EncFtn variants determined by ion mobility mass spectrometry.**

The ion mobility drift time of the 26+ charge state of each EncFtn variant is displayed as a function of increasing collision voltage. All variants display similar CIU profiles, with a compact structural conformation (10.5 ms, highlighted by the letter C) present at low activation which transitions to a more extended conformation (drift time 12.5-13.5 ms, highlighted by the letter E) as activation increases. The onset of this transition occurs at around 32 V in WT, E31A, and E34A; However, the transition occurs at significantly lower activation energy in W38G and the extended structural conformation (12.5 ms) dominates the profile, even at low activation voltages.

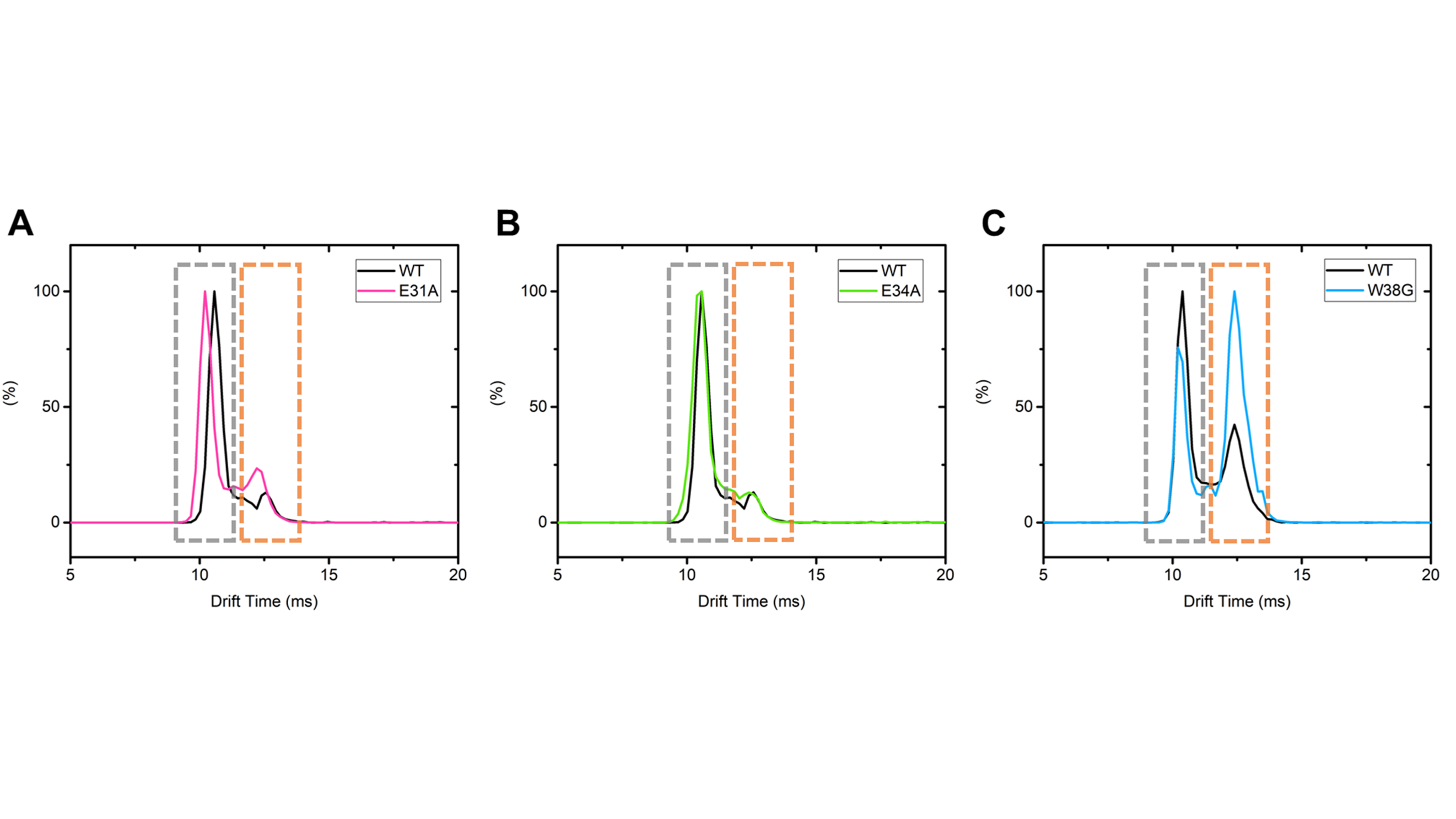

**Figure 5–figure supplement** **3.**

**The extracted ion mobility arrival time distributions of EncFtn variants at 10 V collision voltage from CIU IM-MS experiments.**
In each plot, the decamer form of an EncFtn variant (colored trace) is compared to EncFtn-WT (black trace). (**A**) EncFtn-E31A, pink; (**B**) EncFtn-E34, green; (**C**) EncFtn-W38G, blue. Each arrival time distribution consists of peaks corresponding to a compact structural conformation (10.5 ms, stressed with grey boxes) and extended structural conformation (12.5 ms, highlighted by orange boxes). In WT, E31A, and E34A the compact conformation dominates. However, we note an increase in the extended conformation in the W38G variant. For each panel, comparative data acquired in parallel to control for day-to-day instrument variability.

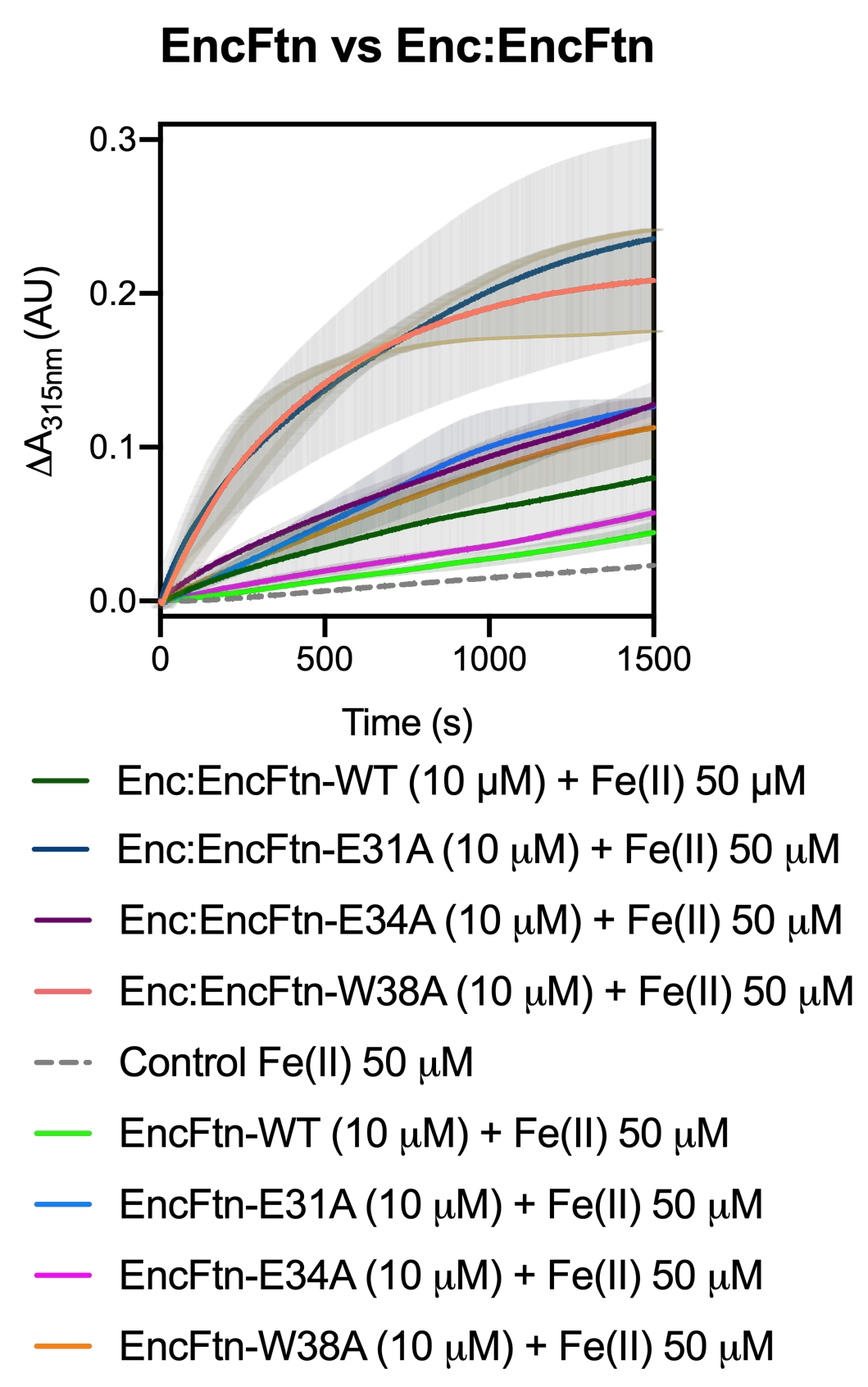

**Figure 6–figure supplement** **1.**

**Ferroxidase activity of EncFtn and Enc:EncFtn encapsulated ferritins.**

Data shown in Figure 6A and B were plotted in the same frame to emphasize the increase in activity induced by the encapsulation of EncFtn proteins by Enc. doi.10.6084/m9.figshare.9885575

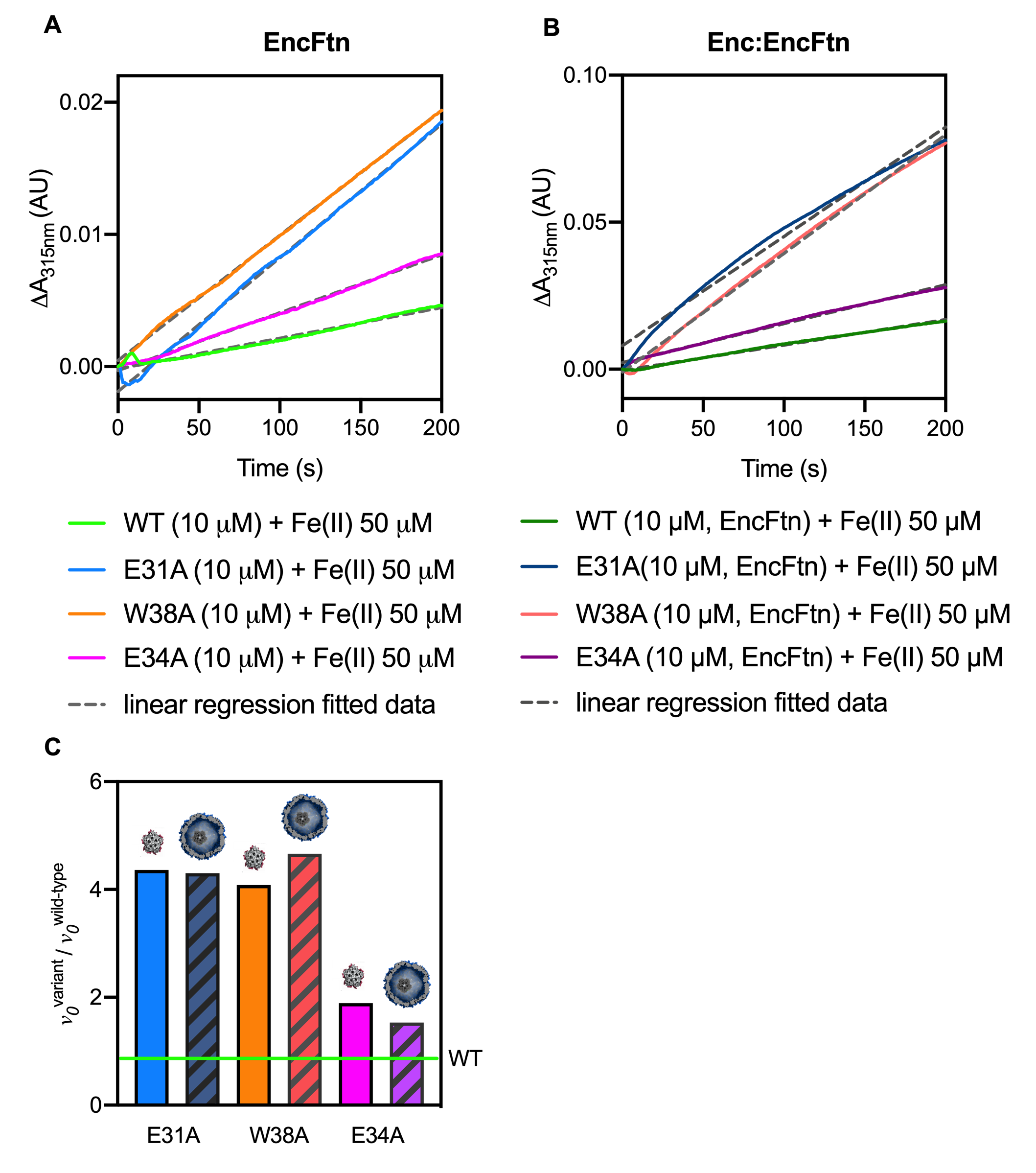

**Figure 6-supplement figure 2.**

**Linear Regression on first 200s of ferroxidase assays with EncFtn and Enc:EncFtn complexes.**

(**A**) Data from first 200 s of the assays shown in Figure 6A were processed using the Linear Regression tool in GraphPad (Prism8) to determine v_0_, initial enzymatic rate, from slope of each of the linear curves (see Table 4). (**B**) Data from first 200 s of the assays shown in Figure 6B were processed using the Linear Regression tool in GraphPad (Prism8) to determine v_0_, initial enzymatic rate, from slope of each of the linear curves (see Table 5). (**C**) Variants initial enzymatic rate (v_0_^variant^) calculated from linear regression curves in the first 200 s of ferroxidase assay experiments were divided by the slope calculated for the corresponding wild-type species (v_0_^wild-type^). Evaluation of the resulting v_0_^variant^/ v_0_^wild-type^ factors allow comparison between enzymatic kinetics of EncFtn (solid colored columns) and Enc:EncFtn (striped columns) proteins. Color coding is consistent with Figures 6A and B and Figure 6-supplement figure 3A and B, and columns are labelled with corresponding mutation. Horizontal green line represents EncFtn-WT’s v_0_^variant^/ *v*_0_^wild-type^ ratio = 1. Representations of EncFtn or Enc:EncFtn systems (non-descriptive of true size ratios) are shown above each column for clarity. doi.10.6084/m9.figshare.9885575

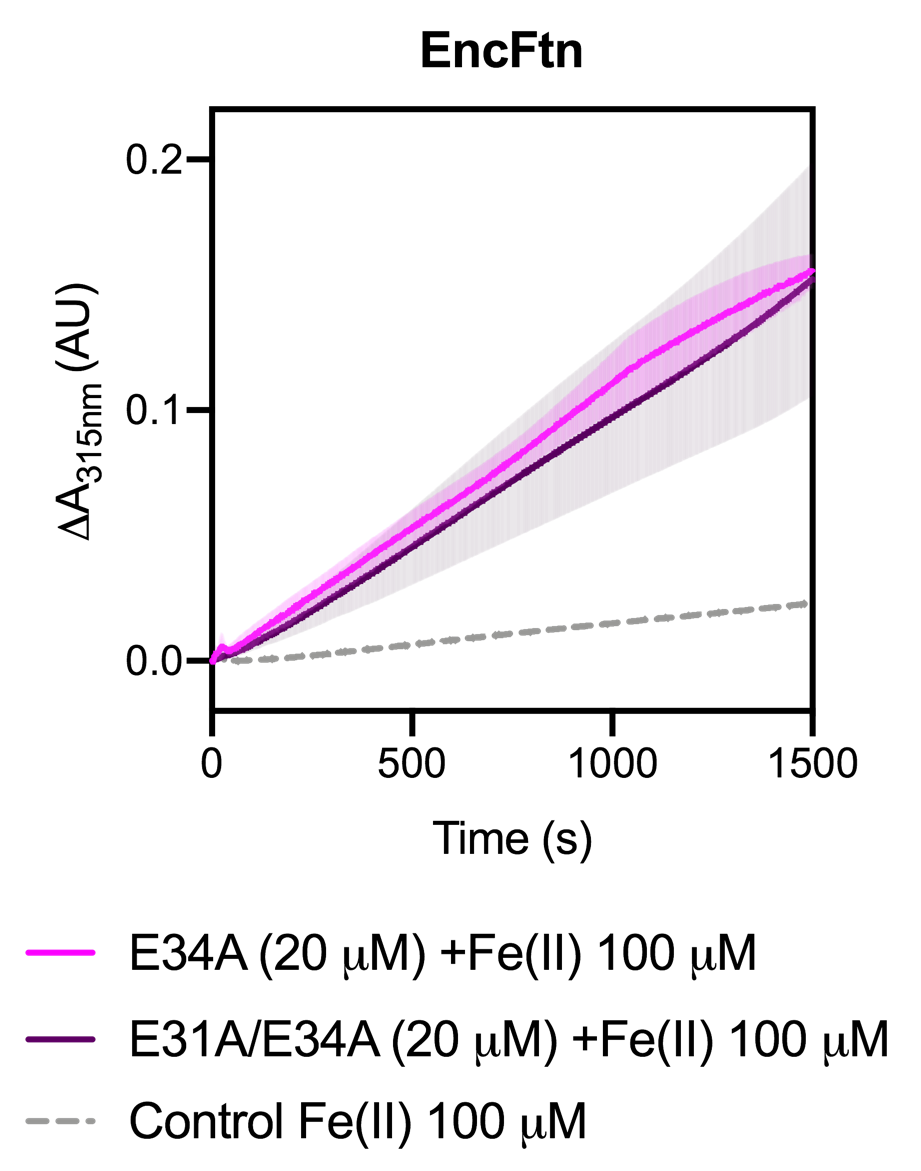

**Figure 6-figure supplement 3.**

**Comparison of ferroxidase activities of EncFtn-E34A and EncFtn-E31A/E34A.**

EncFtn-E34A (pink line) and EncFtn-E31A/E34A (dark purple line), (20 μM, monomer) were incubated with 100 μM FeSO_4_.7H_2_O (10 times molar equivalent Fe(II) per FOC) and progress curves of the oxidation of Fe(II) to Fe(III) was monitored at 315 nm at room temperature. The background oxidation of iron at 100 μM in enzyme-free control is shown for reference (dotted grey line). Solid lines represent the average (n = 3) of technical replicates, shaded areas represent standard deviation from the mean. Protein and iron samples were prepared anaerobically in Buffer H (10 mM HEPES pH 8.0, 150 mM NaCl), and 0.1 % (v/v) HCl, respectively. doi.10.6084/m9.figshare.9885575

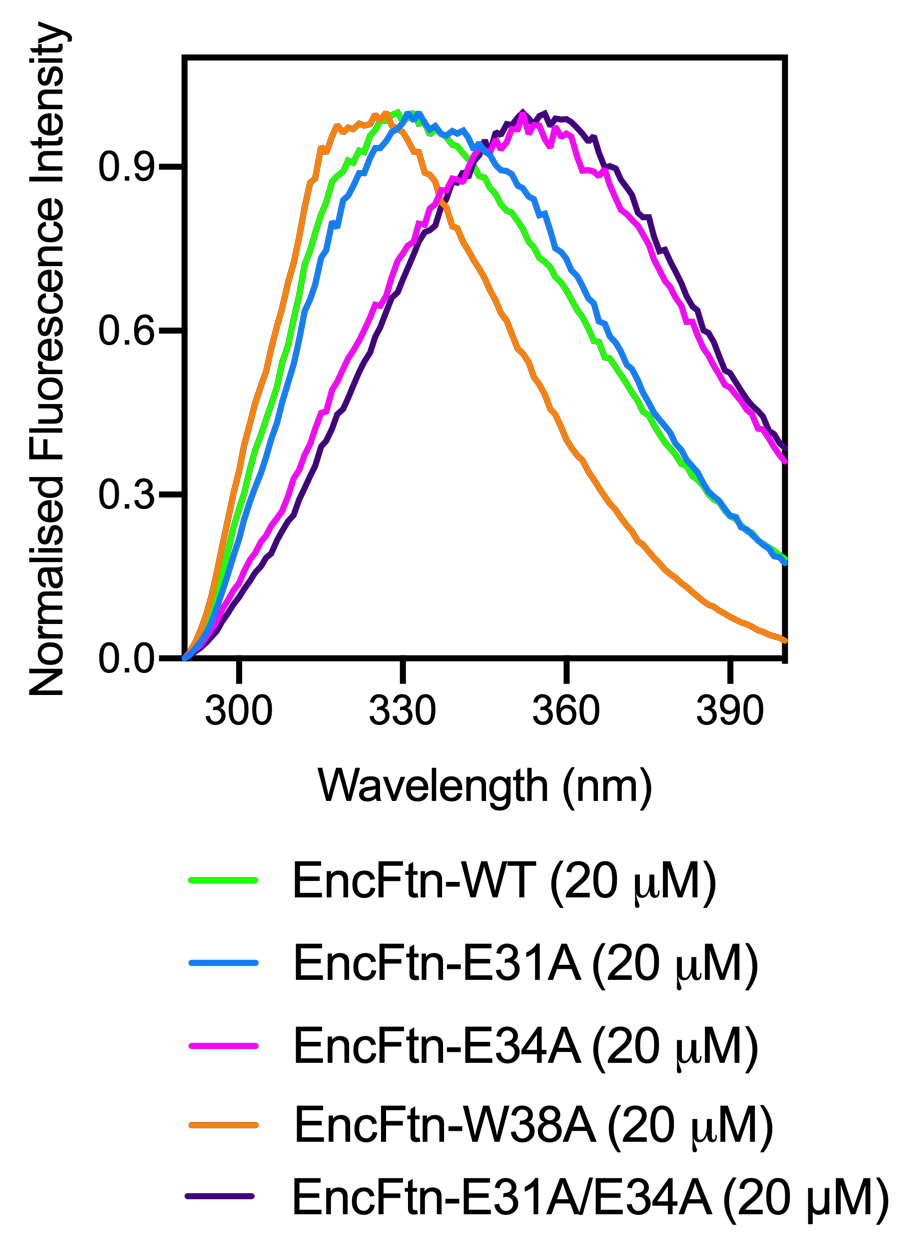

**Figure 8–figure supplement** **1.**

**Normalized intrinsic tryptophan fluorescence emission spectra of EncFtn variants.**

Intrinsic tryptophan fluorescence emission spectra (emission = 290-400 nm, excitation = 280 nm) of EncFtn-WT (green line), EncFtn-E31A (blue line), EncFtn-E34A (pink line), EncFtn-W38A (orange line), EncFtn-E31A/E34A (dark purple line). Proteins were diluted to 20 µM in Buffer GF. Data were normalized to allow comparison. doi.[10.6084/m9.figshare.11920512](https://doi.org/10.6084/m9.figshare.11920512)

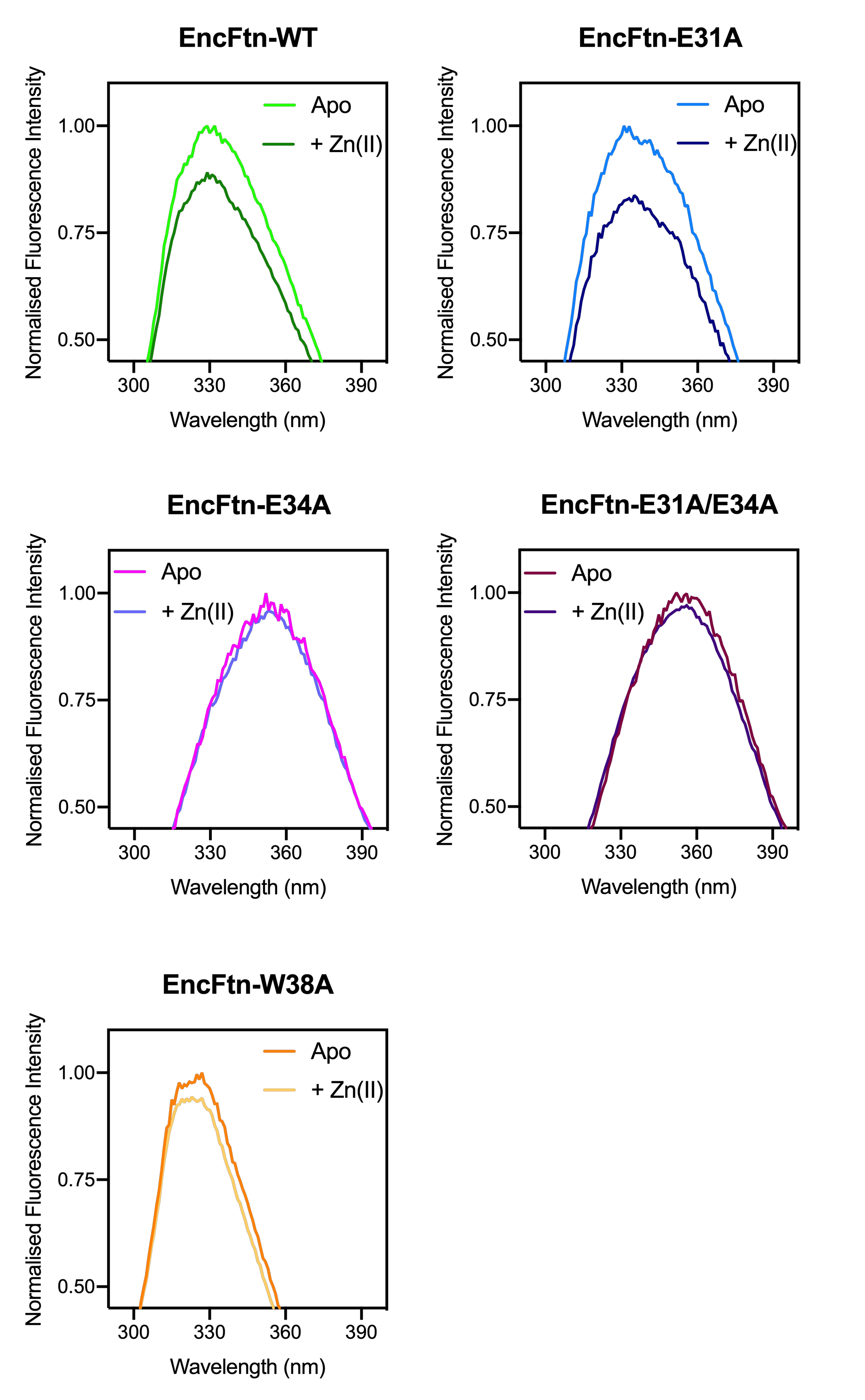

**Figure 8–figure supplement** **2.**

**Normalized intrinsic tryptophan fluorescence emission spectra of EncFtn variants in the presence or absence of Zn(II).**

Intrinsic tryptophan fluorescence emission spectra (emission = 290-400 nm, excitation = 280 nm) shown in Figure 8-figure supplement 1 are presented here alongside corresponding spectra recorded upon addition of 1.5 molecular equivalents of ZnSO_4_.7H_2_O. Final protein concentration in the reaction sample is 20 µM (diluted in Buffer GF), while molecular equivalents of Zn(II) were calculated against concentration of EncFtn entry site (10 µM), with 1.5 equivalents corresponding to 15 µM Zn(II). doi.[10.6084/m9.figshare.11920512](https://doi.org/10.6084/m9.figshare.11920512)

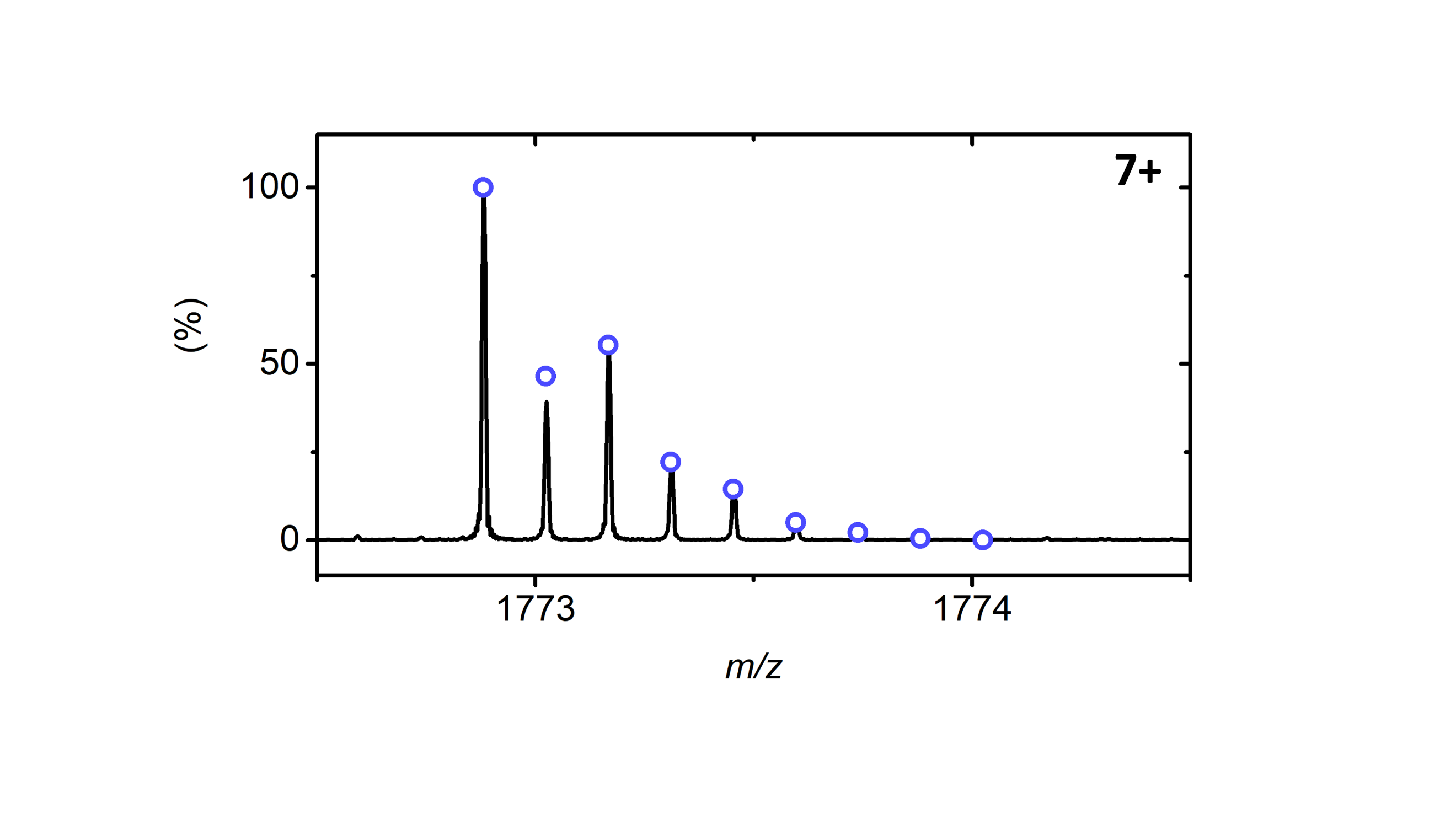

**Figure 9–figure supplement** **1.**

***ID*-EncFtn monomer with theoretical *m/z* value**

Native MS spectrum of the 7+ monomer charge state of apo-*ID*-EncFtn with the theoretical value of the 7+ monomer charge state shown in blue circles.

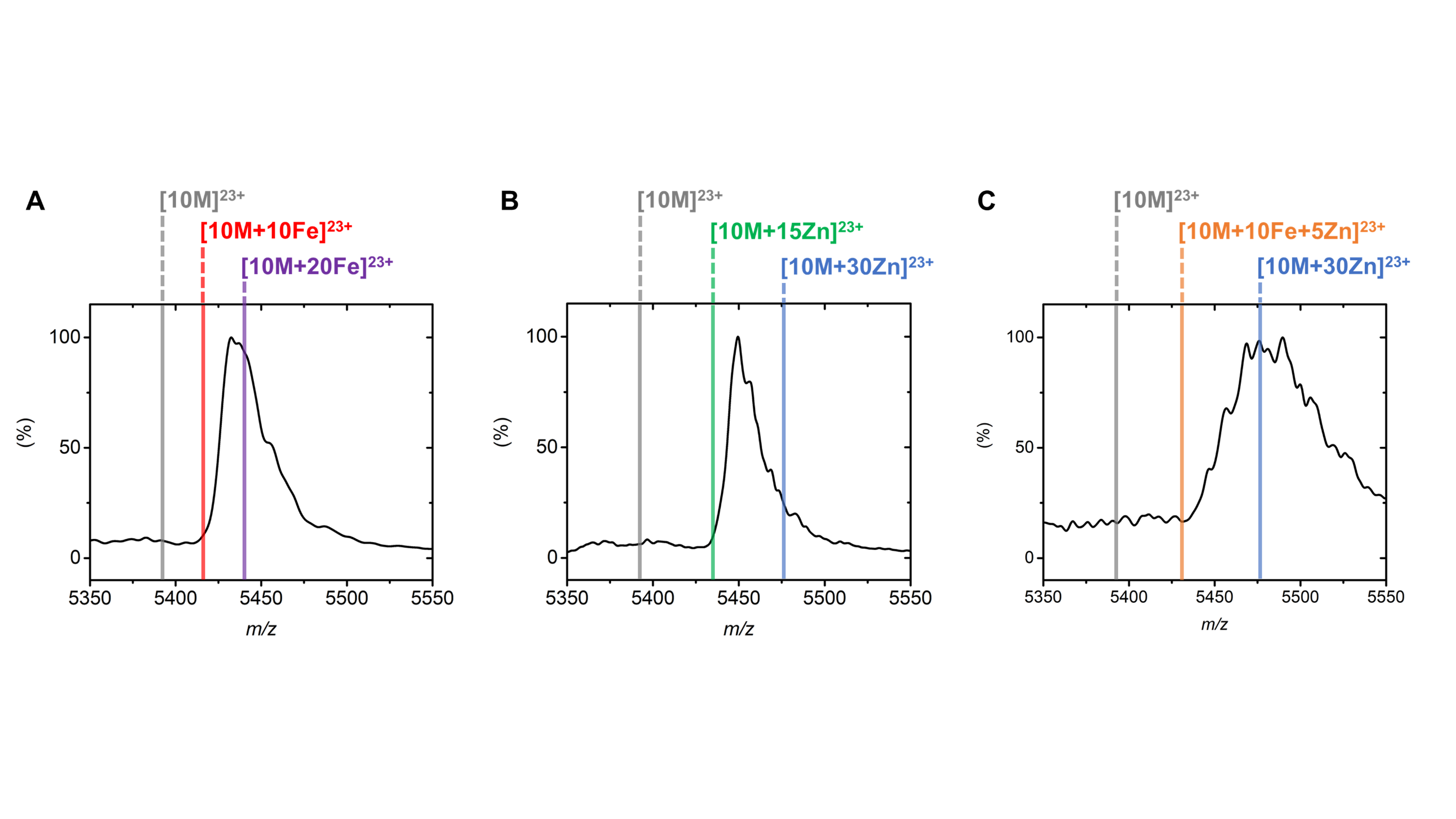

**Figure 9–figure supplement** **2.**

**Metal association of 23+ decamer charge state of *ID*-EncFtn**

(**A**) Iron loading of the 23+ decameric charge state of *ID*-EncFtn showing each decamer binds between 10 and 20 Fe(2+) ions. The grey line represents the theoretical mass of the apo-decamer (*m/z*: 5393.66); the red line indicates the mass of decameric *ID*-EncFtn with 10 Fe(2+) ions bound (*m/z*: 5417.14); and the purple line corresponds to 20 associated Fe(2+) ions (*m/z*: 5440.61). (**B**) Zinc association of *ID*-EncFtn 23+ decamer charge state. Grey line as in (**A**), green line is consistent with the decamer *ID*-EncFtn with 15 Zn(2+) coordinated (*m/z*: 5435.54), and the blue line corresponds to 30 bound Zn(2+) irons (*m/z*: 5476.35). (**C**) Metal association of *ID*-EncFtn titrated with zinc and iron. Grey and blue lines as shown in (**B**). A fully loaded FOC with iron and entry site with bound zinc is shown with an orange line (*m/z*: 5430.92).

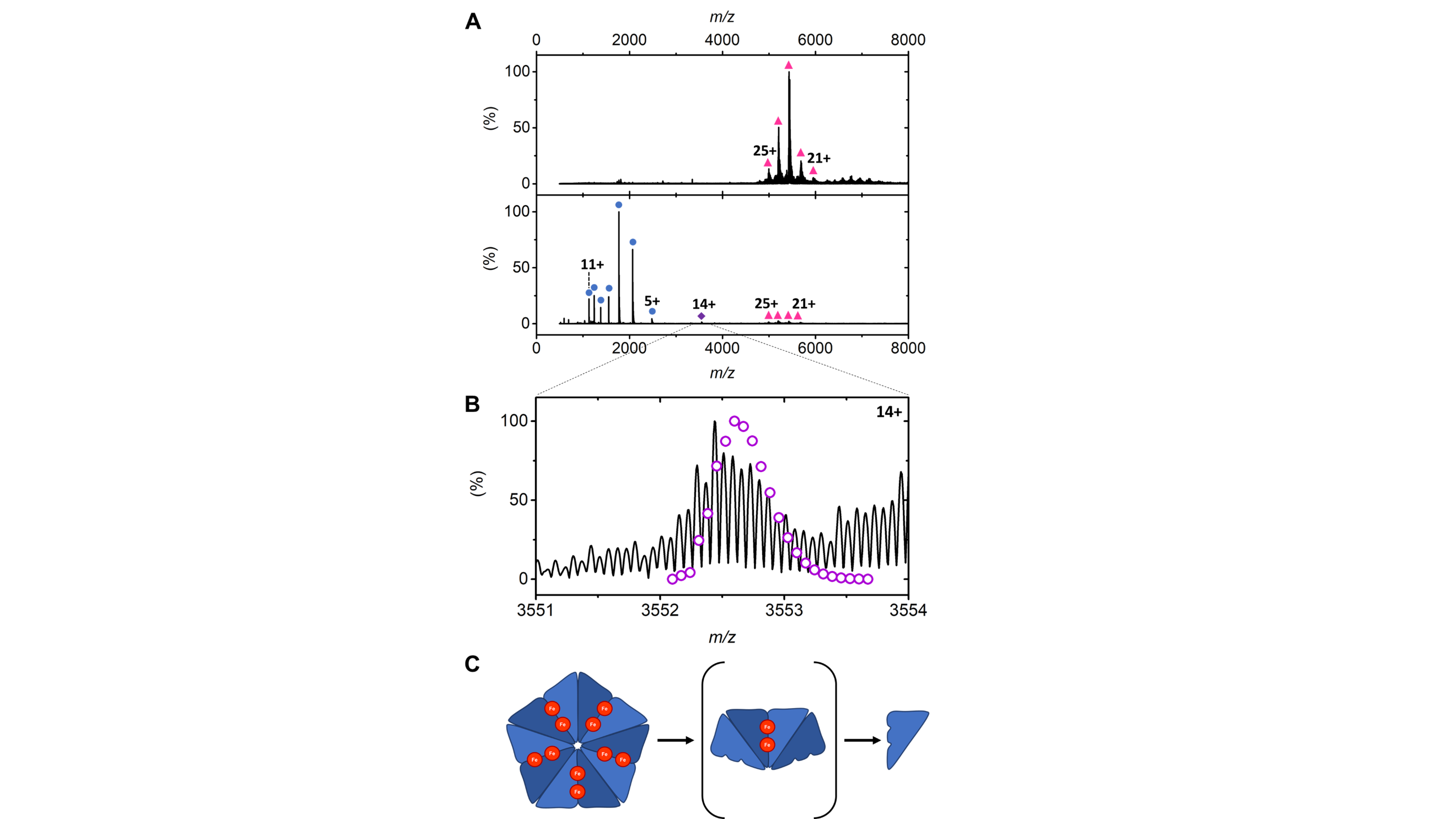

**Figure 9-figure supplement 3.**

**Dissociation pathway of *ID*-EncFtn loaded with iron**

(**A**) Top panel: native mass spectrum of *ID*-EncFtn titrated with Fe(II) in minimally activating MS conditions. Only a decameric oligomerization state is observed (pink triangles). Lower panel: mass spectrum of ID-EncFtn under activating conditions displaying monomer (blue circles), tetramer (purple dimer) and decamer (pink triangles) charge state distributions. (**B**) The 14+ tetramer charge state with a theoretical model of *m/z* values for EncFtn_4_Fe(III)_2_ shown in purple circles. (**C**) Cartoon representation of the dissociation of iron-loaded EncFtn from decamer to monomer or via a minor tetramer species.

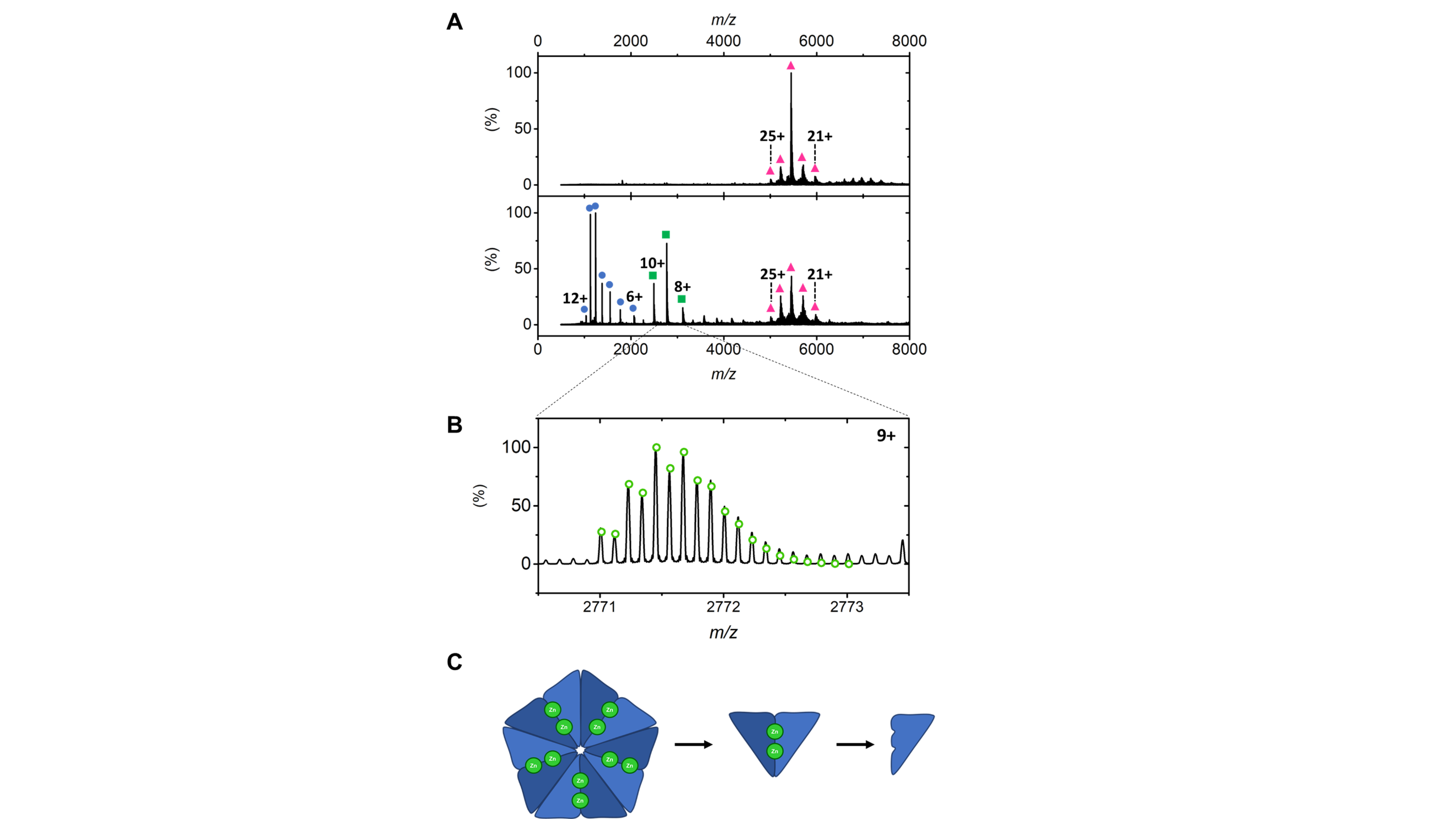

**Figure 9–figure supplement** **4.**

**Dissociation pathway of *ID*-EncFtn loaded with zinc**

(**A**) Dissociation of zinc-loaded *ID*-EncFtn by CID. Top panel displays exclusively decameric species (pink triangles) in minimally activating conditions. Lower panel shows the dissociated species with increased activation of the complex. Gas phase oligomerization states are highlighted by blue circles (monomer), green squares (dimer) and pink triangles (decamer). (**B**) 9+ dimer charge state distribution with theoretical model of *ID*-EncFtn_2_Zn(II)_2_ shown in green circles (**C**) Representation of the dissociation pathway of zinc bound *ID*-EncFtn showing the intermediate FOC dimer species between decamer and monomer.

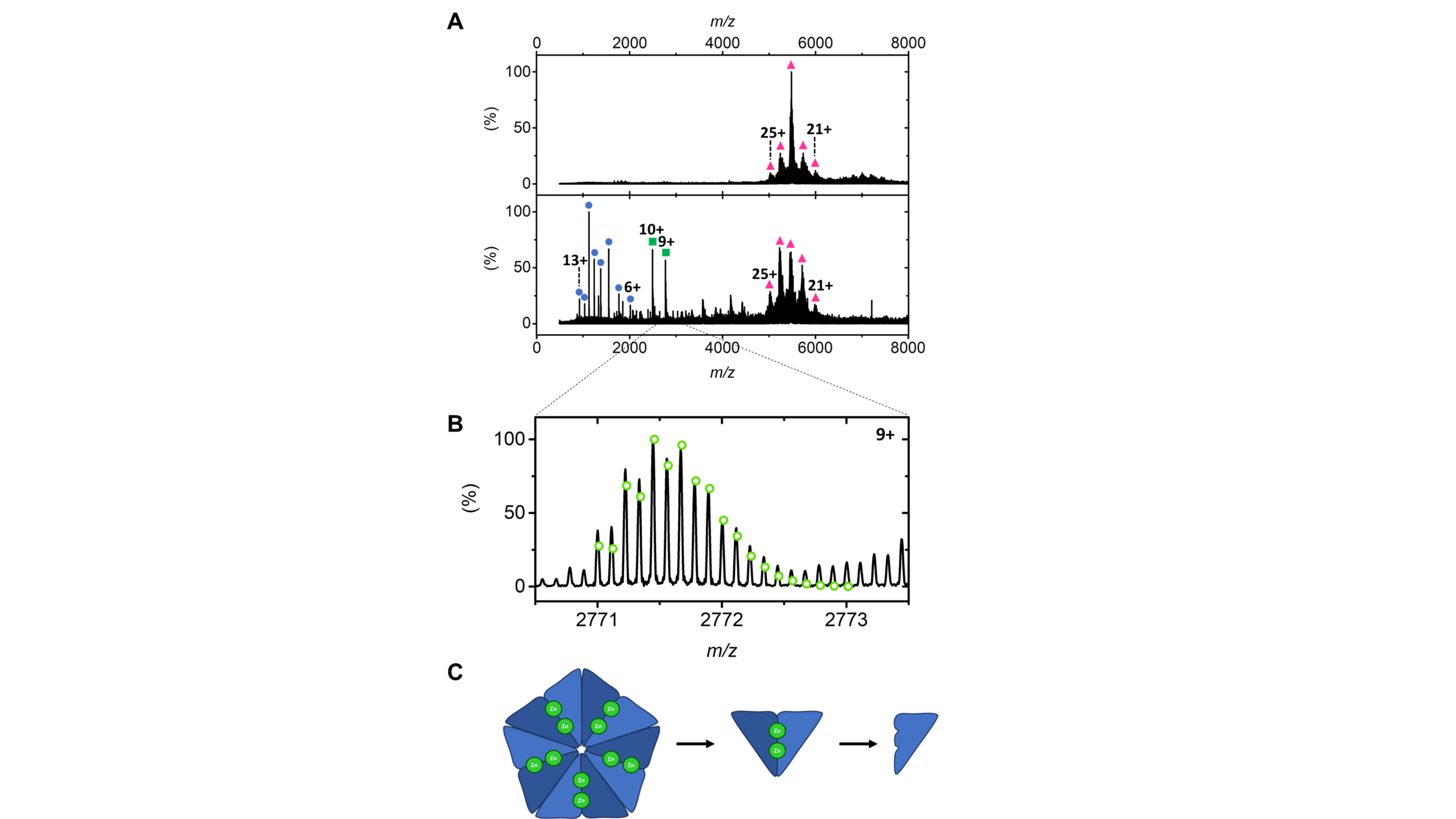

**Figure 9–figure supplement** **5.**

**Dissociation pathway of *ID*-EncFtn loaded with iron and zinc**

(**A**) *ID*-EncFtn titrated with iron then zinc is observed as a decamer in mild MS conditions (top panel). Activation of the complex results in monomer, dimer and decamer species being observed (blue circles, green squares, and pink triangles respectively). (**B**) *ID*-EncFtn_2_Zn_2_ dimer species with corresponding theoretical model in green circles. (**C**) Dissociation pathway of *ID*-EncFtn titrated with iron and zinc
